# Diplopoda in the world fossil record

**DOI:** 10.1101/2024.02.21.581465

**Authors:** Michelle Álvarez-Rodríguez, Francisco Riquelme, Miguel Hernández-Patricio, Fabio Cupul-Magaña

## Abstract

We present a comprehensive catalog with an updated database of the fossil record of Diplopoda in the world. Taxonomic data was collected from descriptions and reports published from 1854 to the present. We also include new records from the Lower Miocene Mexican amber, counting 83 unknown fossil inclusions, with the first records of the orders Polyxenida, Platydesmida, and Julida, as well as the families Sphaeriodesmidae and Trichopolydesmidae within Polydesmida. According to our results, Diplopoda comprises 217 fossil records from the Middle Silurian to Upper Pleistocene, representing three subclasses, six superorders, 25 orders, one superfamily, 54 families, 90 genera, and 156 fossil species. To date, no fossils of the order Siphonocryptida have been reported. The fossil record extends over three geological eras: the Paleozoic, with 156 records; the Mesozoic, with 51; and the Cenozoic, with 77. The fossil preservation includes 87 impressions, 68 compressions, 108 amber inclusions, and 19 ichnites. Thus, this catalog allows us to estimate the size and taxonomic composition of Diplopoda in the fossil record worldwide.

## 1 Introduction

Millipedes (Arthropoda: Myriapoda: Diplopoda) are one of the most successful animal forms in terrestrial environments. Diplopoda in the fossil record extends from the Mid-Paleozoic to the Upper Cenozoic (Wilson & Anderson, 2004; Shear & Edgecombe, 2010). Paleozoic forms include the oldest known land animals from the Middle to Upper Silurian, such as *Casiogrammus ichthyeros* (Wilson, 2005b)*, Kampecaris obanensis* (Brookfield *et al.,* 2020), *Cowiedesmus eroticopodus, Albadesmus almondi*, *Pneumodesmus newmani* (Wilson & Anderson, 2004), and *Eoarthropleura ludfordensis* (Shear & Selden, 1995). Other Paleozoic millipedes of the genus *Arthropleura* from the Upper Carboniferous are considered one of the largest known terrestrial arthropods (Davies *et al*., 2021). In the Permian, at the end of the Paleozoic, a single fossil record is found in North America (Hannibal, 2006; Hannibal & May, 2020). Mesozoic forms are often assigned within existing orders and families (Sierwald & Bond, 2007; Shear *et al*., 2009; Shear & Edgecombe, 2010; Edgecombe, 2015). The Triassic is represented by *Tomiulus angulatus* (Dzik, 1981) and *Hannibaliulus wilsonae* (Shear *et al*., 2009). Little is known about the Jurassic fossil record, there are a few poorly preserved specimens, and their taxonomic identity still needs to be determined (Shear & Edgecombe, 2010). Only the species *Decorotergum warrenae* Jell, 1983 from the Lower Jurassic of Australia is known. From the Upper Mesozoic (Cretaceous) and through the Cenozoic, fossil millipedes are predominantly modern forms preserved as amber inclusions (Shear & Edgecombe, 2010; Edgecombe, 2015). Cretaceous amber inclusions come from different strata, including the most emblematic amber sites, such as Lebanon (Nguyen Duy-Jacquemin & Azar, 2004), France (Nguyen Duy-Jacquemin & Azar, 2004), and Myanmar (Rasnitsyn & Ross, 2000; Grimaldi *et al*., 2002; Ross & Sheridan, 2013; Liu *et al*., 2017; Zhang, 2017; Ross, 2018; Wesener & Moritz, 2018; Jiang *et al*., 2019; Moritz & Wesener, 2019, 2021; Stoev *et al*., 2019; Su *et al*., 2019, 2020). In the Cenozoic, most fossil materials are also preserved as amber inclusions from exceptional conservation sites, such as those in the Baltic region (Koch & Berendt, 1854; Menge, 1854; Hoffman, 1969; Wesener, 2019), India (Srivastava *et al*., 2006), Dominican Republic (Shear, 1981; Santiago-Blay & Poinar, 1992), and Mexico (Riquelme *et al*., 2013; 2014; 2021; Riquelme & Hernández-Patricio, 2018).

Several authors have previously reviewed the taxonomic composition of the Diplopoda fossil record (Almond, 1985; Shear, 1997; Wilson, 2006; Shear & Edgecombe, 2010; Edgecombe, 2015). Initially, Almond (1985) listed the fossil record of Myriapoda, including Diplopoda, establishing a geochronological sequence from the Silurian to the Devonian. Subsequently, Shear (1997) documented the fossil record of Diplopoda, focusing on the most representative higher taxa, also including a geochronological sequence. In addition, Wilson (2006) performed a stratocladogram of Myriapoda, including Diplopoda, combining data from the fossil record with a cladistic analysis (Sierwald *et al*., 2003). Shear & Edgecombe (2010) then chronologically reviewed the fossil record of Diplopoda. Finally, a summary taxonomic list has been presented by Edgecombe (2015).

Other authors have also listed millipedes or have published inventories of specific geological sites. Santiago-Blay & Poinar (1992) reported the orders Polyxenida, Glomeridesmida, Stemmiulida, Spirostreptida, Polyzonida, Siphonophorida, and Polydesmida from Miocene Dominican amber. Štamberg & Zajíc (2008) recorded the Permian-Carboniferous fossil fauna of the Czech Republic, including millipedes. Wesener & Moritz (2018) published an inventory of the Myriapoda, including Diplopoda, from the lowermost Upper Cretaceous amber of Myanmar. Riquelme & Hernández-Patricio (2018) published the first inventory of Diplopoda from the Lower Miocene Chiapas amber. They initially listed 34 records in four orders, six families, five genera, and three species.

This work aims to critically review the Diplopoda fossil record literature to provide a global catalog with an up-to-date database. Taxonomic data were compiled from descriptions and reports published since 1854. We have new data on the fossil record of Diplopoda in Mexican amber from the Miocene, which is presented here as new records added to the catalog. We also include a revised stratocladogram and summary taxonomic list of the worldwide fossil record.

## 2 Materials and methods

The fossil species (valid/named) were compiled from literature published up to November 2023. Also included here are other fossil records not determined to the species level or considered *Incertae sedis*. Each record provides the data of the original description with the author, year, pages, figures, referred material, preservation, repositories, and comments associated with their current status. Referred material identified as adult female ♀, male ♂, and juvenile is indicated in the catalog and database. Undetermined adults and no declared status are shown only in the database. Taxonomic treatment follows the expanded traditional Linnaean classification ranks. The nomenclature and taxonomy follow Sierwald & Bond (2007), Shear (2011), Golovatch (2013); Enghoff et al., 2015; and Edgecombe (2015). Nomenclature and terminology of geological ages follows the International Chronostratigraphic Chart. Additional information about the fossiliferous localities is also presented, including strata, quarries, mines, and geological horizons. Taxonomic references are listed in chronological order. The catalog database is continuously updated on a website (www.riquelmelab.org.mx), which will be helpful to different users and provide a nested system as a basis for future consultation on any taxa of interest within Diplopoda. Therefore, we would appreciate readers’ corrections and additions to this online database. Microsoft Office Excel 2010 was used to analyze the information in the database. Photomicrographs of Mexican amber inclusions were collected by applying multiple image stacking (Z ≥ 30) on a CARL ZEISS® AXIO ZOOM.V16 microscope coupled to an Axiocam MRc5 camera. The ZEN 2012® software and Corel PHOTO-PAINT® were used for image processing and editing (Riquelme *et al*., 2014).

Institutional acronyms and other abbreviations are as follows:

AM.CH—Ámbar de Chiapas, Mexico;

AMNH—American Museum of Natural History, New York, USA;

AMS—Australian Museum, Sydney, Australia;

ANSP—Academy of Natural Sciences of Drexel University, Pennsylvania, USA;

BGS—British Geological Survey, England, UK;

BIRUG—Lapworth Museum of Geology University of Birmingham, England, UK;

BMB—Collection of Writhlington Geological Nature Reserve, England, UK;

BMNH—British Museum Natural History, England, UK;

BNiel—Private collection of Bente Nielsen, Denmark;

BRSMG—Bristol Museum Collection, England, UK;

BSC—Bergschule Saarbrucken Collection, Saarland, Germany;

BSIP—Birbal Sahni Institute of Paleobotany, Uttar Pradesh, India;

BuB—Private Collection of Patrick Müller, Rhineland-Palatinate, Germany;

CaMNH—Carnegie Museum of Natural History, Pennsylvania, USA;

CAMSM—Sedgwick Museum of Earth Sciences, University of Cambridge, England, UK;

CDCGS—Chengdu Center of China Geological Survey, Sichuan, China;

CGH—National Museum of Prague, Prague, Czech Republic;

CPAL-UAEM—Colección de Paleontología, Universidad Autónoma del Estado de Morelos, Morelos, Mexico;

FMNH—Field Museum of Natural History Chicago, Illinois, USA;

GCPE—Colección Museográfica del Grupo Cultural Paleontológico de Elche, Alicante, España;

GC—Private Collection of Carsten Gröhn, Schleswig-Holstein, Germany;

GOP—Personal Collection of George O. Poinar, Jr. at the University of California, Berkeley, California, USA;

GPIH—Geological-Paleontological Institute and Museum, University of Hamburg, currently Centrum of Natural

History (CeNak), Hamburg, Germany;

GSC—Geological Survey of Canada, Ontario, Canada;

HM—Hunterian Museum, University of Glasgow, Scotland, UK;

HNHM—Hungarian Natural History Museum, Budapest, Hungary;

IBGAS—Institute of Biology, Guizhou Academy of Sciences, Guiyang, China;

IGCSE—Institute of Geological Sciences, Edinburgh, Scotland, UK;

IGL-UNAM—Colección Nacional de Paleontología, Instituto de Geología, Universidad Nacional Autónoma de México, Mexico City, Mexico;

IRSNB—Institut Royal des Sciences Naturelles de Belgique, Bruxelles, Belgique;

IUT—Institut Universitaire de Technologie Louis Pasteur, Strasbourg, France;

JB—Private Collection of Jake Brodzinsky at the Smithsonian Institution, Washington DC, USA;

KUMIP—Division of Invertebrate Paleontology, University of Kansas Biodiversity Institute, Kansas, USA;

LACM—Los Angeles County Museum of Natural History, California, USA;

MACH—Museo del Ámbar de Chiapas, San Cristóbal de las Casas, Chiapas, Mexico;

MALM—Museo del Ámbar Lilia Mijangos, San Cristóbal de las Casas, Chiapas, Mexico;

MCZ—Museum of Comparative Zoology, Harvard University, Massachusetts, USA;

MfN—Museum of Natural History of Berlin, Berlin, Germany;

MfNC—Museum für Naturkunde in Chemnitz, Chemnitz, Germany;

MHN—Natural History Museum of Lille, Lille, France;

ML—Musée de Lodève, Lodève, France;

MM—Manchester Museum, University of Manchester, England, UK;

MMG—Staatliche Museen zu Berlin, Berlin Germany;

MMRB—Muzeum Mineralów, Robert Borzêcki, Klodzko, Poland;

MNCR—Museo Nacional de Costa Rica, San José, Costa Rica;

MNHN—National Museum of Natural History of France, Paris, France;

MPW—Museum of Porcelain in Wałbrzych, Silesia, Poland;

MU—Marshall University, Department of Geology, West Virginia, USA;

NBMG—New Brunswick Museum of Natural History, New Brunswick, Canada;

NGIP—Nanjing Institute of Geology and Palaeontology Chinese Academy of Sciences, Nanjing, China;

NHM—Natural History Museum, London, UK;

NHMW—Naturhistorisches Museum Vienna, Vienna, Austria;

NHV—Natural History Association of Rhineland and Westphalia eV, Bielefeld, Germany;

NMMNH—New Mexico Museum of Natural History, New Mexico, USA;

NMP—National Museum, Prague, Czech Republic;

NMS—National Museums Scotland, UK;

OU—Sam Noble Oklahoma Museum of Natural History, Oklahoma, USA;

PAS—PaleoAmbienti Sulcitani Museum E.A. Martel, Serdegna, Italy;

PC—Private Collection of Patrick Craig, localition unknown;

PIN—Babii Kamen (PIN collection 1062), Kemerovo, Russian Federation;

PPC—Private Collection of Martin Pavela, Opava, Czech Republic;

QMF—Queensland Museum Geosciences Collection, Queensland, Australia;

QM—Queensland Museum, Queensland, Australia;

RGM—Rijks Geologisch Mineralogische, Leiden, Nederland;

RO—Private Collection of Rainer Ohlhoff, Saarland, Germany;

SMNS—Staatliches Museum für Naturkunde Stuttgart, Baden-Wurtemberg, Germany;

SMU—Southern Methodist University collection, Texas, USA;

SSM—Springfield Science Museum, Massachusetts, USA;

SUCCINUM.INAH—Private Collection certified by the Instituto Nacional de Antropología e Historia (INAH), San Cristóbal de las Casas, Chiapas, Mexico;

UCMP—University of California Museum of Paleontology, California, USA;

USMN—Smithsonian Institution, National Museum of Natural History, Washington DC, USA;

WPM—Museum für Naturkunde und Vorgeschichte, Saxony-Anhalt, Germany;

Wu—Private Collection of Jörg Wunderlich, Baden-Wurttemberg, Germany;

YPM—Peabody Museum of Natural History, Yale University, Connecticut, USA;

ZCM—National History Museum of the Pilsen Region, Pilsen, Czech Republic;

ZFMK-MYR—Myriapoda Collection of the Zoological Research Museum Alexander Koenig, North Rhine-Westphalia, Germany;

ZPAL—Palaeozoological Institute of the Polish Academy of Sciences, Warsaw, Poland.

## 3 Catalog

### 3.1 General

This updated catalog comprises three subclasses, six superorders, 25 orders, one superfamily, 54 families, 90 genera, and 156 fossil species in 217 records worldwide (Tables 1-3, Figs 1-2). The fossil record extends over three geological eras: Paleozoic (156 records), Mesozoic (51 records), and Cenozoic (77 records). Fossil preservation consists of compressions, impressions, and ichnites found primarily in Paleozoic and Mesozoic sediments and amber inclusions from late Mesozoic and Cenozoic sites (Table 1, Fig.2).

**Figure 1.**
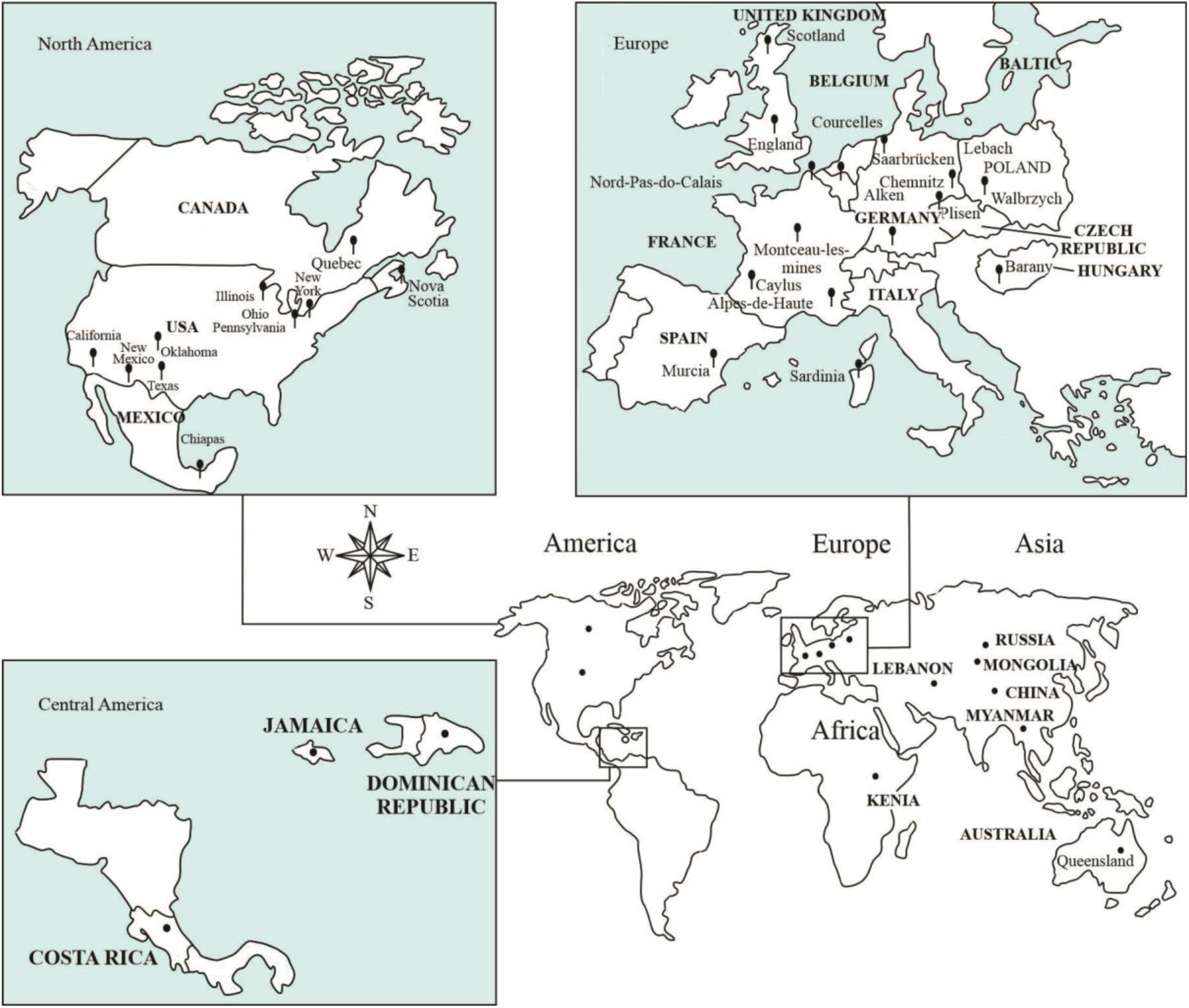
Fossil record of millipedes around the world. Schematic map showing geological sites.

**Figure 2.**
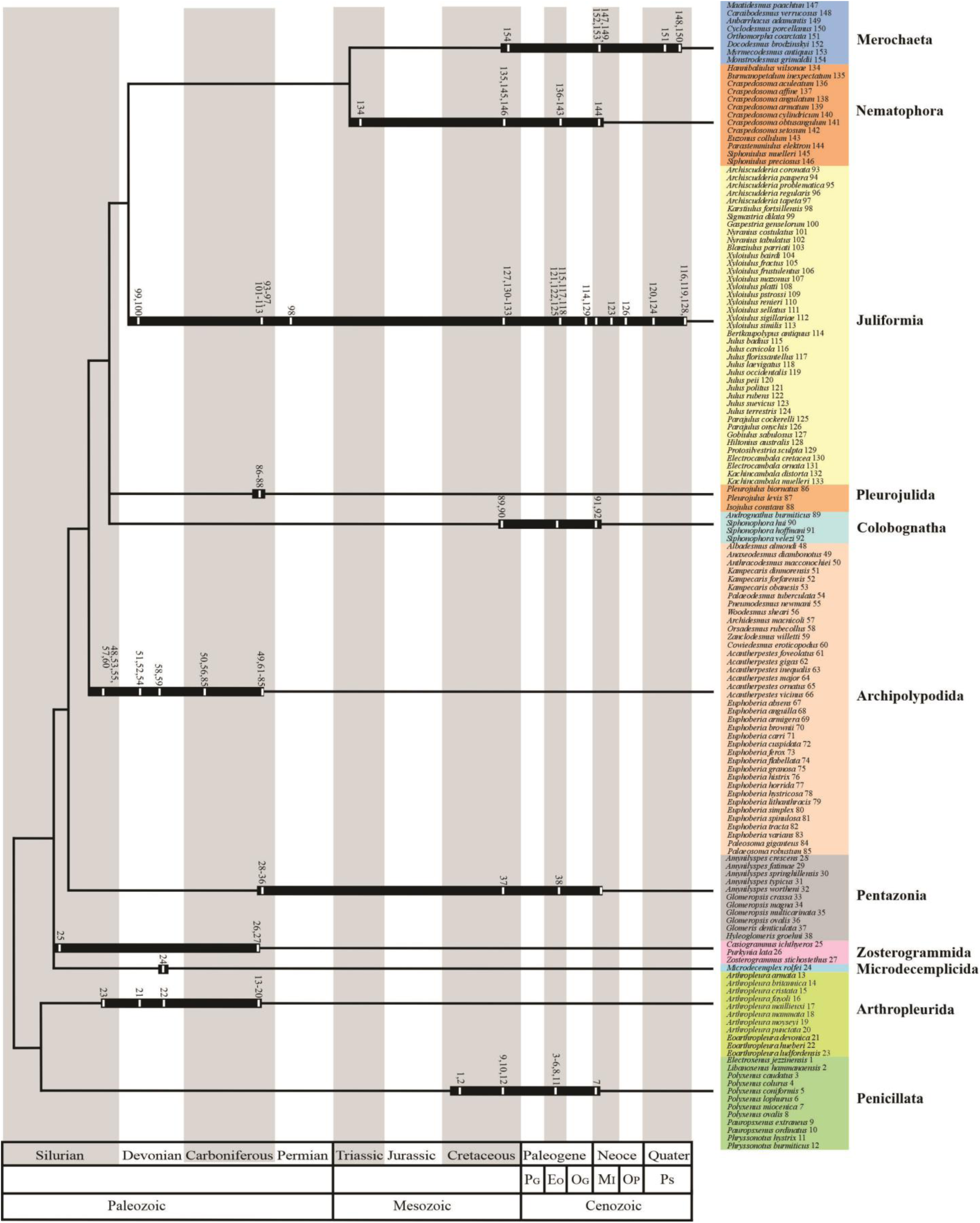
Stratocladogram of the class Diplopoda showing the fossil record from the Middle Silurian to the Upper Pleistocene. Time scale adapted from the ICS International Chronostratigraphic Chart. Phylogenetic position of the higher taxa, including those extinct, follows Wilson (2006), Sierwald & Bond (2007), and Shear & Edgecombe (2010). Range extension is indicated by bold lines and ghost lineages by narrow lines. Taxa are shown on the branches by a number associated with the taxonomic list.

**Table 1.** Current count of millipedes in the worldwide fossil record.

| Era | Fossil record |
| --- | --- |
| Paleozoic | 156 |
| Mesozoic | 51 |
| Cenozoic | 77 |
| Rank |  |
| Subclass | 3 |
| Superorder | 6 |
| Order | 25 |
| Superfamily | 1 |
| Family | 54 |
| Genus | 90 |
| Species | 156 |
| Preservation |  |
| Amber inclusion | 108 |
| Compression | 68 |
| Impression | 87 |
| Ichnites | 19 |

**Table 2.** The fossil record of the class Diplopoda: orders, families, genera, and species, except Siphonocryptida.

|  | Order | Family | Genus | Species |
| --- | --- | --- | --- | --- |
| 1 | Polyxenida Verhoeff, 1934 | 3 | 8 | 12 |
| 2 | †Arthropleurida Waterlot, 1934 | 1 | 1 | 8 |
| 3 | †Eoarthropleurida Shear & Selden, 1995 | 1 | 1 | 3 |
| 4 | †Microdecemplicida Wilson & Shear, 2000 | 1 | 1 | 1 |
| 5 | †Zosterogrammida Wilson, 2005 | 1 | 3 | 3 |
| 6 | †Amynilyspedida Hoffman, 1969 | 2 | 2 | 9 |
| 7 | Glomeridesmida Cook, 1895 | 1 | 1 | - |
| 8 | Glomerida Brandt, 1833 | 2 | 3 | 2 |
| 9 | Sphaerotheriida Brandt, 1833 | 1 | - | 0 |
| 10 | †Archidesmida Wilson & Anderson, 2004 | 2 | 3 | 3 |
| 11 | †Cowiedesmida Wilson & Anderson, 2004 | 1 | 1 | 1 |
| 12 | †Euphoberiida Hoffman, 1969 | 1 | 3 | 23 |
| 13 | †Palaeosomatida Hannibal & Krzeminski, 2005 | 1 | 1 | 2 |
| 14 | †Pleurojulida Schneider & Werneburg, 1998 | 1 | 2 | 3 |
| 15 | Platydesmida Cook, 1895 | 1 | 2 | 1 |
| 16 | Polyzoniida Cook, 1895 | 2 | 2 | - |
| 17 | Siphonophorida Newport, 1844 | 2 | 2 | 3 |
| 18 | Julida Brandt, 1833 | 2 | 3 | 13 |
| 19 | Spirobolida Cook, 1895 | 2 | 2 | 2 |
| 20 | Spirostreptida Brandt, 1833 | 3 | 4 | 5 |
| 21 | Callipodida Pocock, 1894 | 1 | 2 | 2 |
| 22 | Chordeumatida Pocock, 1894 | 2 | 2 | 8 |
| 23 | Stemmiulida Cook, 1895 | 1 | 1 | 1 |
| 24 | Siphoniulida Cook, 1895 | 1 | 1 | 2 |
| 25 | Polydesmida Leach, 1815 | 9 | 16 | 8 |

**Table 3.** Diplopoda fossil species.

| Period | Silurian | Devonian | Carboniferous | Permian | Triassic | Jurassic | Cretaceous | Eocene | Oligocene | Miocene | Pliocene | Pleistocene |
| --- | --- | --- | --- | --- | --- | --- | --- | --- | --- | --- | --- | --- |
| Species | 7 | 10 | 73 | 4 | 3 | 1 | 16 | 22 | 2 | 9 | 1 | 8 |

#### Paleozoic

The earliest fossil forms of Diplopoda are found in the Middle to Upper Silurian rocks. It mainly comprises species within extinct higher taxa (Table 2, Fig.2). One of the oldest fossil records is *Casiogrammus ichthyeros* from the Middle Silurian Fish Bed Formation in Scotland, ca. 430 Ma (Wilson, 2005b). This is a putative millipede of the order Zosterogramida whose record was questioned (Brookfield *et al*., 2020); see below specific comments in the list section. Another oldest uncontroversial fossil record is *Kampecaris obanesis* Peach, 1889 (superorder Archipolypoda) from the Upper Silurian Kerrera Sandstone Formation in Scotland, ca. 425 Ma (Brookfield *et al*., 2020). Other ancient Archipolypoda records are *Albadesmus almondi*, *Cowiedesmus eroticopodus*, and *Pneumodesmus newmani* from the Upper Silurian Cowie Formation in Scotland, ca 414 Ma (Wilson & Anderson, 2004), and *Eoarthropleura ludfordensis* (order Eoarthropleurida) from the Upper Silurian Downton Castle Sandstone Formation in England, ca. 420 Ma (Shear & Selden, 1995).

Devonian diplopods comprise two superorders, three orders, one superfamily, four families, eight genera, and 10 species (Table 3, Fig.2), such as Archipolypoda: *Kampecaris dinmorensis* from the Lower Devonian Herefordshire in England, *Kampecaris forfarensis* from the Lower Devonian Old Red Sandstone in Scotland (Almond, 1985), and *Palaeodesmus tuberculata* from the Lower Devonian Dunure in Scotland (Wilson & Anderson, 2004); the extinct order Microdecempicida: *Microdecemplex rolfei* from the Middle Devonian Panther Mountain Formation in the USA (Wilson & Shear, 2000); the extinct order Eoarthropleurida: *Eoarthropleura devonica* from the Lower Devonian Nellen Koepfchen Beds in Germany (Størmer, 1976) and *Eoarthropleura hueberi*, described from the Upper Devonian Onteora Red Beds Formation in the USA (Kjellsvig-Waering, 1986); Archidesmida: *Orsadesmus rubecollus* from the Upper Devonian Catskill Formation in USA and *Zanclodesmus willetti* from the Upper Devonian Escuminac Formation in Canada (Wilson *et al*., 2005); the extinct superfamily Xyloiuloidea (superorder Juliformia): *Sigmastria dilate* from the Lower Devonian Dundee Formation in Scotland and *Gaspestria genselorum* from the Lower Devonian Battery Point Formation in Canada (Wilson, 2006).

The Carboniferous has the most extensive fossil diversity of millipedes compared to any other period, with 73 species, 20 genera, 12 families, one superfamily, six orders, and two superorders (Table 3, Fig.2). The orders Arthropleurida, Zosterogrammida, Amynilypedida, Euphoberiida, Palaeosomatida, Pleurojulida, and the superfamily Xyloiuloidea were recorded. In the Lower Carboniferous, three helminthomorph species have been described: *Anthracodesmus macconochiei* from the Lower Carboniferous Lennel Braes strata in Scotland (Peach, 1899), *Woodesmus sheari* from Lower Carboniferous Ballagan Formation in Scotland (Ross *et al*., 2018) and *Palaeosoma robustum* from Lower Carboniferous Walbrzych Formation in Poland (Jackson *et al*., 1919). The Upper Carboniferous is characterized by a diverse array of higher taxa, including several extinct orders and families. Among the orders, Arthropleurida, Zosterogrammida, Amynilypedida, Euphoberiida, Palaeosomatida, and Pleurojulida are recorded, and families include Nyraniidae, Proglomeridae, Archiulidae (order *Incertae sedis*), and the superfamily Xyloiulidae. The Upper Carboniferous has significant fossil diversity of genera and species. Arthropleurida counts one family, one genus (*Arthropleura*), and eight species from the Lower Carboniferous to Lower Permian (Davies *et al*., 2021). In contrast, only four species of helminthomorph millipedes are found in the Permian: *Dolesea subtila*, *Oklahomasoma richardsspurense*, *Karstiulus fortsillensis* from the Lower Permian Richards Spur in the USA (Hannibal & May, 2020), and *Archiulus brassi* from the Lower Permian Saarlouis Lebach strata in Germany (Guthörl, 1934).

#### Mesozoic

The fossil record of Diplopoda is limited to four species from the Triassic to Jurassic (Fig.2): *Tomiulus angulatus* from the Lower Triassic Maltsevo Formation in Russia (Martynov, 1936), *Hannibaliulus wilsonae* from the Middle Triassic Grès à Voltzia Formation in France (Shear *et al*., 2009), *Sinosoma luopingense* from the Middle Triassic Guanling Formation in China (Huang & Hannibal, 2018) and the only species found in the Jurassic, *Decorotergum warrenae* from the Lower Jurassic Evergreen Formation in Australia (Jell, 1983).

However, diversity increases significantly with family-level radiation in the Upper Cretaceous to the end of this era. The records of families, genera, and species expanded considerably in the Cretaceous. The order Polyxenida is recorded in the Lower Cretaceous amber of Lebanon, with one family, two genera, and two species: *Electroxenus jezzinensis* and *Libanoxenus hammanaensis* (Nguyen Duy-Jacquemin & Azar, 2004). The genus *Phryssonotus* (family Synxenidae) is recorded from the Upper Cretaceous amber of France (Nguyen Duy-Jacquemin & Azar, 2004). Lowermost Upper Cretaceous (Upper Albian/Lower Cenomanian) amber of Myanmar recorded 14 orders, one suborder, 15 families, 14 genera, and 13 species (Cockerell, 1917; Liu *et al*., 2017; Zhang, 2017; Ross, 2018; Wesener & Moritz, 2018; Jiang *et al*., 2019; Moritz & Wesener, 2019; Stoev *et al*., 2019; Su *et al*., 2020; Moritz & Wesener, 2021; Su *et al*., 2022).

#### Cenozoic

The modern forms are generally found in this era. The variation is observed at the genus and species level. In the Eocene, six orders, eight families, ten genera, and 22 species are recorded (Fig.2). The Upper Eocene from Baltic amber comprises six orders, six families, nine genera, and, 20 species. Other two species were described from the Upper Eocene Florissant Formation in the USA: *Julus florissantellus* (Cockerell, 1907) and *Parajulus cockerelli* (Miner, 1926).

In the Oligocene, two juliform species have been described: *Bertkaupolypus antiquus* (Julida: Julidae) from the Upper Oligocene Rott Formation in Germany (Hoffman, 1969) and *Protosilvestria sculpta* (Spirostreptida: Cambalidae) from the Upper Oligocene in France (Mauries, 1992).

In the Miocene, *Polyxenus miocenica* is described from the Lower Miocene amber of India (Srivastava *et al*., 2006). In this period, the most remarkable fossil diversity is found predominantly in amber sites from southern Mexico and the Dominican Republic. The superorder Juliformia, the orders Siphonophorida, Spirobolida, Stemmiulida, and Polydesmida, the families Siphonophoridae, Stemmiulidae, Xystodesmidae, Platyrhacidae, Chelodesmidae, and Pyrgodesmidae, the genera *Siphonophora*, *Parastemmiulus*, *Anbarrhacus*, *Maatidesmus*, and *Myrmecodesmus*; and the species *Parastemmiulus elektron*, *Anbarrhacus adamantis*, *Maatidesmus paachtun*, and *Myrmecodesmus antiquus*, are described from the Lower Miocene Mexican amber (Riquelme & Hernández-Patricio, 2018; Riquelme *et al*., 2021). This work adds 83 fossil inclusions that were recently recovered in Mexican amber. Polyxenida, Platydesmida, and Julida are reported for the first time, as well as new records of two Polydesmida families: Sphaeriodesmidae and Trichopolydesmidae (Figs 3-4). Thus, the current inventory in Mexican amber includes 117 fossils in one superorder, seven orders, eight families, five genera, and four species (Fig.5). On the other hand, five orders, seven families, eleven genera, and three species*: Siphonophora hoffmani*, *Siphonophora velezi* (Santiago-Blay & Poinar, 1992), and *Docodesmus brodzinskyi* (Shear, 1981), are recorded from the Lower Miocene amber of the Dominican Republic. There is only one record from the Upper Miocene, a juliform species *Julus suevicus* from the Upper Miocene Thermalsinterkalk Formation in Germany (Dietlen, 1902).

**Figure 3.**
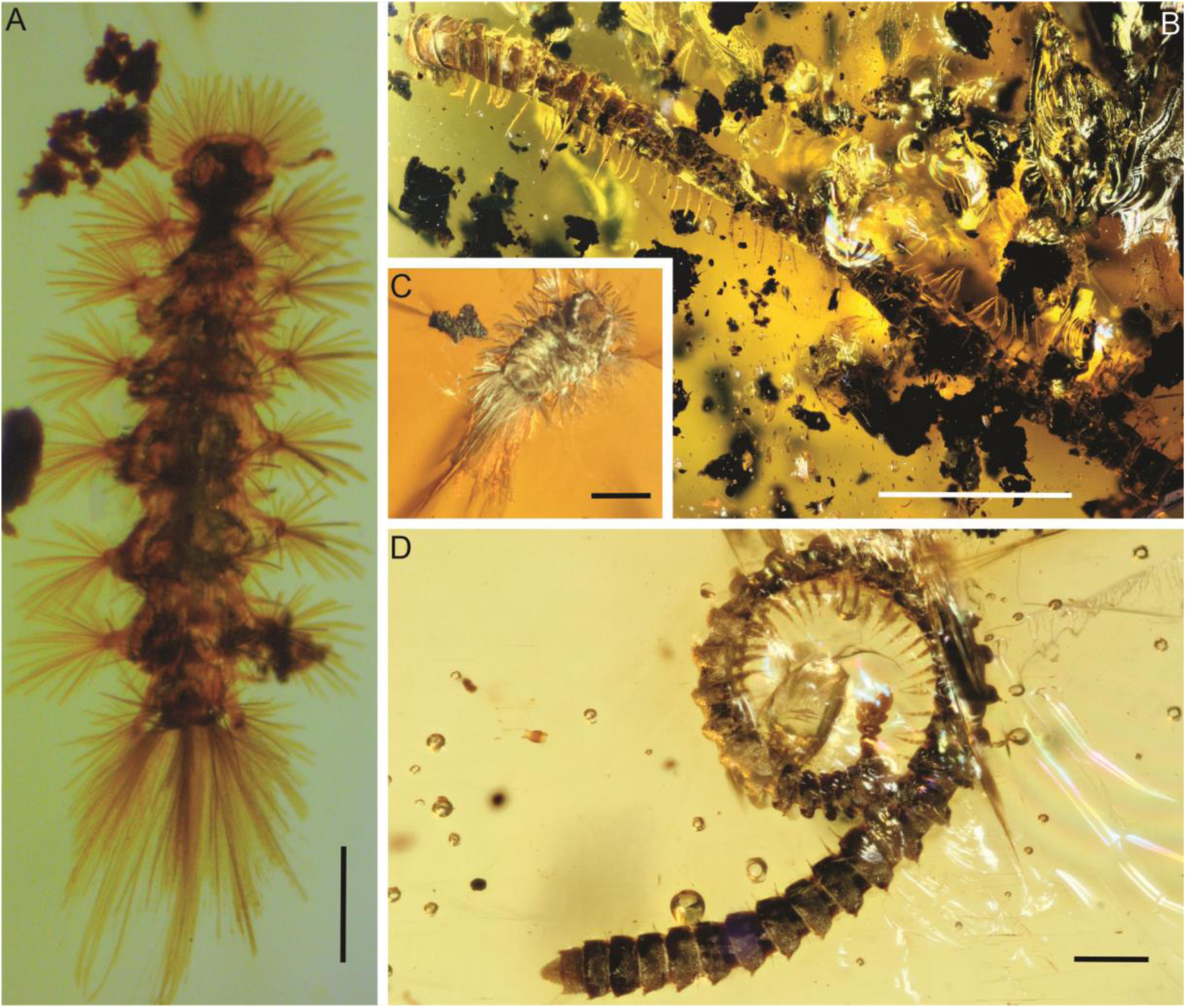
New records of Diplopoda from Mexican amber, Simojovel Formation, Lower Miocene. A. CPAL.143: Polyxenida. B. CPAL.124: Julida. C. CPAL.111: Polyxenidae. D. CPAL.157: Platydesmidae. Scale bars: A = 0.5 mm; B = 0.3 mm; C = 0.2 mm; D= 1.0 mm.

**Figure 4.**
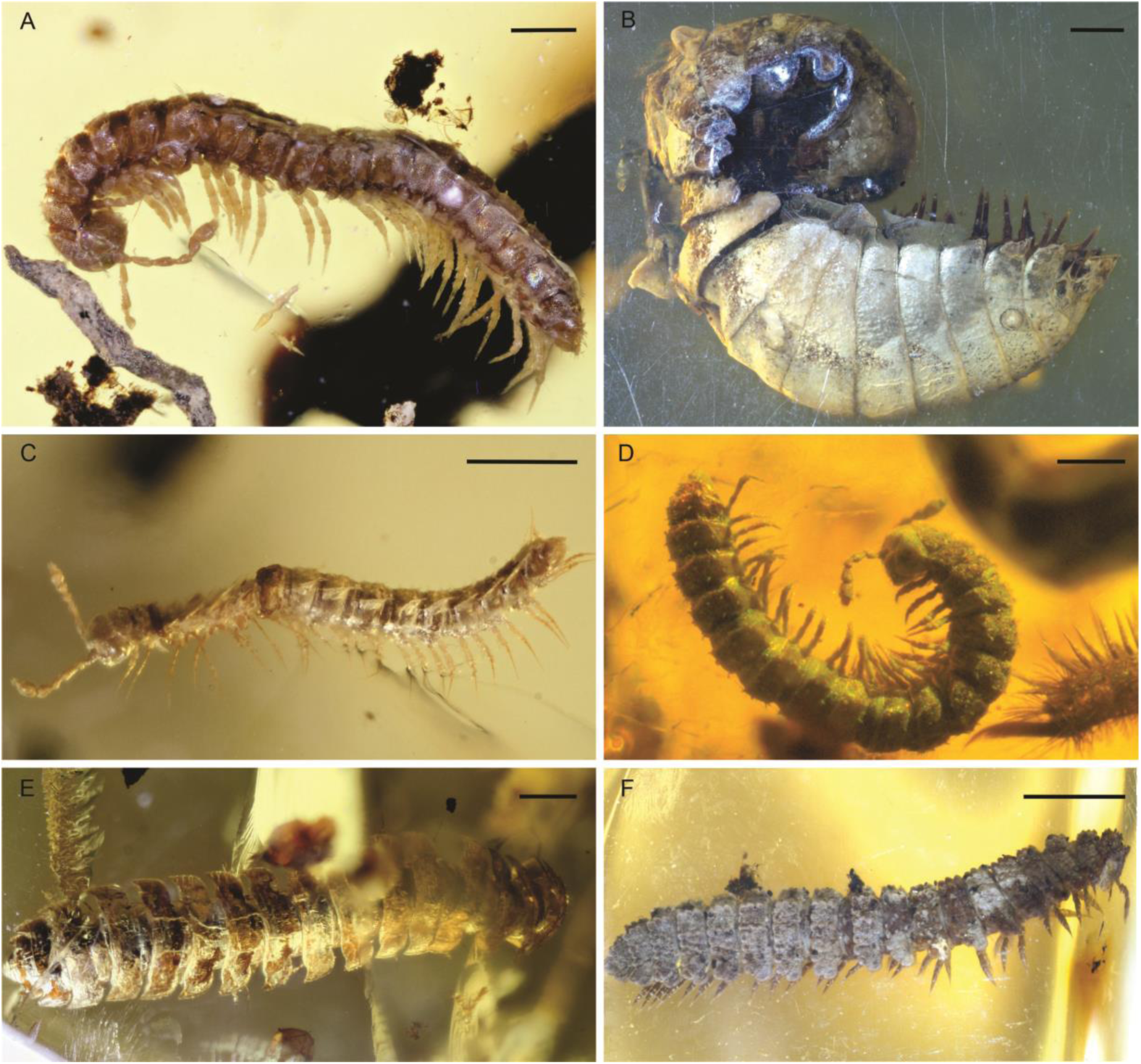
New records of Diplopoda from Mexican amber, Simojovel Formation, Lower Miocene. A. CPAL.121: Trichopolydesmidae (Polydesmida). B. CPAL.130: Sphaeriodesmidae (Polydesmida). C. CPAL.150: Trichopolydesmidae (Polydesmida). D. CPAL.125.1: Polydesmida indet. E. CPAL.137: Polydesmida indet. F. CPAL.138: Pyrgodesmidae (Polydesmida). Scale bars: A, E = 0.5 mm; B, F= 1 mm; C–D = 0.2 mm.

**Figure 5.**
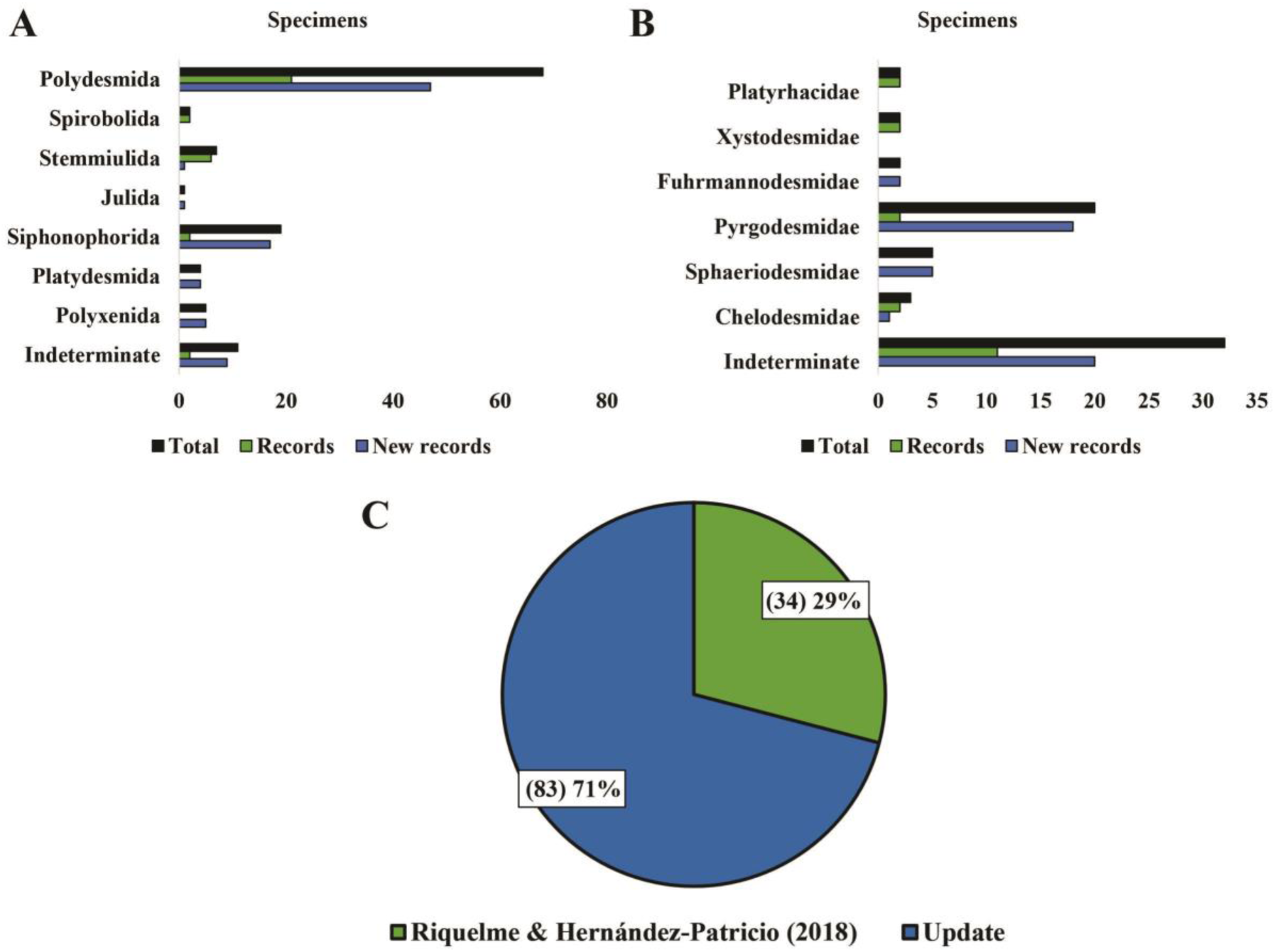
Diplopoda from Mexican amber, Simojovel Formation, Lower Miocene. A. Orders. B. Families of Polydesmida. C. First inventory (2018) and updated (2023).

In the Pliocene, *Parajulus onychis* is the sole record from the Lower Pliocene Onyx Marble Formation in the USA (Pierce, 1951). Finally, the youngest fossil diversity in Pleistocene sites consists of 11 records, with three orders, six families, seven genera, and five species: *Julus cavicola*, *Julus occidental,* and *Hiltonius australis* from the Upper Pleistocene Rancho La Brea in the USA (Grinell, 1908); as well as *Caraibodesmus verrucosus* (Donovan & Veltkamp, 1994) and *Cyclodesmus porcellanus* (Baalbergen & Donovan, 2013) from the Upper Pleistocene Red Hills Road Cave in Jamaica (Table 3).

### 3.2 Systematic Palaeontology

Phylum Arthropoda Gravenhorst, 1843

Clade Mandibulata *sensu* Snodgrass, 1938

Subphylum Myriapoda Latreille, 1802

Class Diplopoda de Blainville *In*: Gervais, 1844

#### 3.2.1 Subclass Penicillata Latrielle, 1831

**Order Polyxenida Verhoeff, 1934**

Polyxenida indet: Wesener & Moritz, 2018: 1132; Álvarez-Rodríguez *et al*., 2023, Fig.3A (this work).

Referred material: BuB2658; BuB2659; BuB2984; BuB3028; RO my295. Preservation: Amber inclusions. Repository: BuB; RO. Locality: Myanmar, Kachin, Hukawng Valley, Noije Bum mine. Horizon: No stated, lowermost Upper Cretaceous, Upper Albian/Lower Cenomanian.

New records: CPAL.112; CPAL.126; CPAL.143. Preservation: Amber inclusions. Repository: CPAL-UAEM. Locality: Mexico, Chiapas, Simojovel, Guadalupe Victoria mine (CPAL.112), Los Pocitos mine (CPAL.126), Monte Cristo mine (CPAL.143). Horizon: Uppermost Simojovel Formation, Lower Miocene.

**Superfamily Polyxenoidea Lucas, 1940**

**Family Lophoproctidae Silvestri, 1897**

**Genus *Lophoproctus* Pocock, 1894**

*Lophoproctus* sp.: Santiago-Blay & Poinar, 1992: 363, Fig.1.

Referred material: GOP DC-3-14; PC: 1 ♀ plus other 3 specimens. Preservation: Amber inclusions. Repository: GOP; PC. Locality: Dominican Republic, Santiago, Cordillera Septentrional. Horizon: El Mamey Formation, Lower Miocene.

**Family Polyxenidae Lucas, 1840**

Polyxenidae indet: Zhang, 2017: 144-145. Wesener & Moritz, 2018: 1137. Álvarez-Rodríguez *et al*., 2023, Fig.3C (this work).

Referred material: No stated. BuB634; BuB2612; BuB2961; BuB2966; Wu F3358/Bu/CJW; Wu F3384/Bu/ CJW; Wu F3389/Bu/CJW; Wu F3394/Bu/CJW. Preservation: Amber inclusions. Repository: NHML. BuB; Wu. Locality: Myanmar, Kachin, Hukawng Valley, Noije Bum mine. Horizon: No stated, lowermost Upper Cretaceous, Upper Albian/Lower Cenomanian.

New records: CPAL.111: juvenile; CPAL.209: juvenile. Preservation: Amber inclusions. Repository: CPAL-UAEM. Locality: Mexico, Chiapas, Totolapa, Río Salado mine (CPAL.111) and Simojovel, Monte Cristo mine (CPAL.209. Horizon: Uppermost Simojovel Formation, Lower Miocene.

Comments: Zhang (2017: 144-145) tentatively identified the genera *Unixenus*, *Propolyxenus*, and *Polyxenus* within Polyxenida. After analyzing Zhang’s (2017: 154-155) photomicrographs, Wesener and Moritz (2018) found a misidentification, but consider the record of the family Polyxenidae to be valid.

†Genus *Electroxenus* Nguyen Duy-Jacquemin & Azar, 2004

**(1) †*Electroxenus jezzinensis* Nguyen Duy-Jacquemin & Azar, 2004**

*Electroxenus jezzinensis* Nguyen Duy-Jacquemin & Azar, 2004: 632, Figs 1-2, 633-634.

Referred material: Holotype: MNHN Acra JS 231/1. Preservation: Amber inclusion. Repository: MNHN. Locality: Lebanon, Jezzine, Mouhafazit Loubnan Al-Janoubi. Horizon: No stated, Lower Cretaceous.

**†Genus *Libanoxenus* Nguyen Duy-Jacquemin & Azar, 2004**

**(2) †*Libanoxenus hammanaensis* Nguyen Duy-Jacquemin & Azar, 2004**

*Libanoxenus hammanaensis* Nguyen Duy-Jacquemin & Azar, 2004: 635, Figs 3-4, 636-637.

Referred material: Holotype: MNHN Azar 633. Preservation: Amber inclusion. Repository: MNHN. Locality: Lebanon, Hammana, Caza of Baabda, Mouhafazit Jabal Loubnan. Horizon: No stated, Lower Cretaceous.

**Genus *Polyxenus* Latrielle, 1802**

*Polyxenus* sp.: Hoffman, 1969: 583, Fig.368; Ross, 2018: 38.

Referred material: No stated. Preservation: Amber inclusion. Repository: No stated. Locality: Baltic. Horizon: Prussian Formation, Upper Eocene.

Referred material: No stated. Preservation: Amber inclusion. Repository: NMS. Locality: Myanmar, Kachin, Hukawng Valley. Horizon: No stated, lowermost Upper Cretaceous, Upper Albian/Lower Cenomanian.

**(3) †Polyxenus caudatus Menge, 1854**

*Polyxenus caudatus* Menge, 1854: 12.

Referred material: No stated. Preservation: Amber inclusion. Repository: WPM. Locality: Baltic. Horizon: Prussian Formation, Upper Eocene.

**(4) †Polyxenus colurus Menge, 1854**

*Polyxenus colurus* Menge, 1854: 12.

Referred material: No stated. Preservation: Amber inclusion. Repository: WPM. Locality: Baltic. Horizon: Baltic, Prussian Formation, Upper Eocene.

**(5) †*Polyxenus coniformis* Koch & Berendt, 1854**

*Polyxenus coniformis* Koch & Berendt, 1854: 11, Fig.133.

Referred material: No stated. Preservation: Amber inclusion. Repository: Berendt collection (currently in MfN).

Locality: Baltic. Horizon: Prussian Formation, Upper Eocene.

**(6) †Polyxenus lophurus Menge, 1854**

*Polyxenus lophurus* Menge, 1854: 12.

Referred material: No stated. Preservation: Amber inclusion. Repository: WPM. Locality: Baltic. Horizon: Prussian Formation, Upper Eocene.

**(7) †Polyxenus miocenica Srivastava et al., 2006**

*Polyxenus miocenica* Srivastava *et al*., 2006: 717, Figs 3-4, 718.

Referred material: Holotype: BSIP 38375-B. Preservation: Amber inclusion. Repository: BSIP. Locality: India, Kerala, Payangadi Clay Mines. Horizon: Warkalli Formation, Lower Miocene, Burdigalian.

**(8) †*Polyxenus ovalis* Koch & Berendt, 1854**

*Polyxenus ovalis* Koch & Berendt, 1854: 12, Fig.3.

Referred material: No stated. Preservation: Amber inclusion. Repository: MfN. Locality: Baltic. Horizon: Prussian Formation, Upper Eocene.

**Genus *Propolyxenus* Silvestri, 1948**

*Propolyxenus* sp.: Ross, 2018: 38.

Referred material: No stated. Preservation: Amber inclusion. Repository: NMS. Locality: Myanmar, Kachin, Hukawng Valley, Noije Bum mine. Horizon: No stated, lowermost Upper Cretaceous, Upper Albian/Lower Cenomanian.

**Genus *Pauropsxenus* Silvestri, 1948**

**(9) †Pauropsxenus extraneus Su et al., 2020**

*Pauropsxenus extraneus* Su *et al*., 2020: 4, Figs 4, 6.

Referred material: Holotype: NGIP168241: ♂; Paratypes: NGIP168242: ♀; NGIP168243: ♀; NGIP168244: sub-♀; additional material: NGIP168245-NGIP168255: 2 ♀; 4 specimens; 2 sub-♂; 1 sub-♀; 2 juveniles; additional with 6 poorly preserved specimens. Preservation: Amber inclusions. Repository: NGIP. Locality: Myanmar, Kachin, Hukawng Valley, Noije Bum mine. Horizon: No stated, lowermost Upper Cretaceous, Upper Albian/Lower Cenomanian.

**(10) †Pauropsxenus ordinatus Su et al., 2020**

*Pauropsxenus ordinatus* Su *et al*., 2020: 2, Figs 1-3, 3-5.

Referred material: Holotype: NGIP168231: ♂; Paratype: NGIP168232: ♀; aditional material: NGIP168233-NGIP168240: ♀, 3 specimens, sub-adult; 2 juveniles (stage V). Preservation: Amber inclusions. Repository: NGIP. Locality: Myanmar, Kachin, Hukawng Valley, Noije Bum mine. Horizon: No stated, lowermost Upper Cretaceous, Upper Albian/Lower Cenomanian.

**Genus *Unixenus* Jones, 1944**

*Unixenus* sp.: Ross, 2018: 38.

Referred material: No stated. Preservation: Amber inclusion. Repository: NMS. Locality: Myanmar, Kachin, Hukawng Valley. Horizon: No stated, lowermost Upper Cretaceous, Upper Albian/Lower Cenomanian.

**Superfamily Synxenoidea Silvestri, 1923**

**Family Synxenidae Silvestri, 1923**

Synxenidae indet: Rasnitsyn & Ross, 2000: 24. Grimaldi *et al*., 2002: 10. Ross & Sheridan, 2013: 50, Fig.15.

Referred material: In.19102-3: 4 specimens; In. 19104-6: 6 specimens; In.19117-22: 2 specimens; In.19123: 24 specimens; In.20149: 5 specimens; In.20150: 18 specimens; In.20169: 1 specimen. Twenty-four uncoded specimens in NHML. NMS material is not stated. Preservation: Amber inclusions. Repository: NHML. NMS. Locality: Myanmar, Kachin, Hukawng Valley. Horizon: No stated, lowermost Upper Cretaceous, Upper Albian/Lower Cenomanian.

**Genus *Phryssonotus* Scudder, 1885**

*Phryssonotus* sp.: Menge, 1854: 12; Ross & Sheridan, 2013: 50, Figs 15, 50; Nguyen Duy-Jacquemin & Azar, 2004: 636, Figs 5-6, 638-639.

Referred material: No stated. Preservation: Amber inclusions. Repository: WPM. No stated. Locality: Baltic. Horizon: Prussian Formation, Upper Eocene.

Referred material: Holotype: SA 5.1: sub-♀. Preservation: Amber inclusion. Repository: MNHN. Locality: France, Salignac, Alpes-de-Haute-Provence. Horizon: No stated, lowermost Upper Cretaceous, Upper Albian/Lower Cenomanian.

**(11) †Phryssonotus hystrix (Menge, 1854)**

*Lophonotus hystrix* Menge, 1854: 12.

*Phryssonotus hystrix*: Scudder, 1885: 731.

Referred material: No stated. Preservation: Amber inclusion. Repository: WPM. Locality: Baltic. Horizon. Prussian Formation, Upper Eocene.

**(12) †Phryssonotus burmiticus (Cockerell, 1917)**

*Polyxenus burmiticus* Cockerell, 1917: 40-41, Fig.1; Conde, 1954: 75; Conde & Jacquemin, 1963: 69; Zherikhin, 1978: 114; Ross & York, 2000: 15, Fig.18; Nguyen Duy-Jacquemin & Geoffroy, 2003: 101; Wesener & Moritz, 2018: 1132; Su *et al*., 2019: 221, Figs 1-6, 217-222.

Referred material: Wu F3388/Bu/CJW; RO my107; RO my191. NIGP167287: ♂; NIGP167288: ♂; NIGP167289: ♂; NIGP167290: ♂; NIGP167291: ♂; NIGP167292: ♀; NIGP167293: sub-?; NIGP167294: sub-♀; NIGP167295; NIGP167296: sub-♀; NIGP167297: sub-♀; NIGP167298: juvenile; NIGP167299: juvenile; NIGP167300: juvenile; NIGP167301. Preservation: Amber inclusions. Repository: Wu; RO; NIGP. Locality: Myanmar, Kachin, Hukawng Valley, Noije Bum mine. Horizon: No stated, lowermost Upper Cretaceous, Upper Albian/Lower Cenomanian.

#### 3.2.2 †Subclass Arthropleuridea Waterlot, 1934

**†Order Arthropleurida Waterlot, 1934**

**†Family Arthropleuridae Scudder, 1885**

**†Genus *Arthropleura* Jorden & Meyer, 1854**

*Arthropleura* sp.: Andrée, 1913: 298. Sterzel, 1918: 211, taf. 14, Figs 5-7, 10; Röβler & Schneider, 1997: 20, Figs 16-18; Pruvost, 1919: 70-84, pl XXV, Figs 10, 20; Pruvost, 1930: 177; Richardson, 1956: 72-76, Figs 39-40; Novozhylov, 1962: 11, Figs 12A-B; De La Comble, 1963: 6; Ferguson, 1966: 130, Fig.2; Calder *et al*., 2005: 152. Briggs *et al*., 1979: 276, pl 28; Briggs *et al*., 1984: 852, table 1; Falcon-Lang *et al*., 2015: 62, table 2; Ryan, 1986: 156; Ryan & Boehner, 1994: 10; Walter & Gaitzsch, 1988: 73-84; Pearson, 1992: 129, Figs 2-3; Briggs & Almond, 1994: 127-135; Perrier & Charbonnier, 2014: 11, Fig.4M; Anderson *et al*., 1997: 203, Figs 3d-f; Castro, 1997: 20, Lám. I, Figs 1, 1a, 2, and 2a; Schneider & Barthel, 1997: 191, Fig.3; Schneider *et al*., 2010: 55, Figs 7F, 8; Langiaux & Sotty, 1977: 74-91; Briggs, 1986: 141-147; Schneider & Werneburg, 1998: Fig.15; Proctor, 1998: 92, table 1, Figs 2-3; Mangano *et al*., 2002: 35, Figs 32A-C; Lucas *et al*., 2005a: 152, Fig.6; Lucas *et al*., 2005b: 279, Fig.2; Falcon-Lang *et al*., 2006: 566, Fig.6, tables 2-3; Falcon-Lang & Miller, 2007: 949, table 2; Štamberg & Zajíc, 2008: 81; Martino & Greb, 2009, 141, Figs 3-4, 6-7; Nelikhov, 2010: 60-69; Pacyna *et al*., 2012: 122, Fig.3A-C; Röβler *et al*., 2012: 819, Fig.12D; Chaney *et al.,* 2013: 64, Fig.2; Getty *et al*., 2017: 188, Fig.3; Pavela, 2018: Figs 1, 5-6; Whyte, 2018: 63, Figs 2-3; Dernov *et al*., 2019: 51, Figs 6-7; Moreau *et al*., 2019: 5, Figs 4, 6-7, table 3; Davies *et al*., 2021: 6, Figs 1-2.

Referred material: No stated. Preservation: Impressions. Repository: No stated. Locality: UK, Yorkshire, Barnsley; Poland, Lower Silesia, Chwałowice; Germany, Saarland, Saarbrücken. Horizon: Pennine Middle Coal Measures Formation, Upper Carboniferous, Bashkirian; Ruda Beds, Upper Carboniferous, Bashkirian; Saarbrücker Subgroup, Upper Carboniferous, Moscovian.

Referred material: MfNC F 1128; MfNC F 1129. MfNC F 1125; MfNC F 439a, b; MfNC F 447; MfNC 445; MMG SaKa 11/3; MfNC F 442; MMG SaKa 2; MMG SaKa 5; MMG SaKa 10. Preservation: Compressions. Repository: MfNC. MMG. Locality: Germany, Saxony, Hainichen Basin. Horizon: Berthelsdorf Formation, Lower Carboniferous, Visean.

Referred material: n° 1874; n° 1876; n° 1877. Preservation: Impressions. Repository: MHN. Locality: France, Nord-Pas-de-Calais, Saint-Charles vein, Lens mines, pit 7 (n° 1874), Douchy mines, Douchy pit, 248 m depth (n° 1876), Lens mines, pit 11, bowette 115 (n° 1877). Horizon: Shale strata, Upper Carboniferous.

Referred material: No stated. Preservation: Compression. Repository: IRSNB. Locality: Belgium, Courcelles, Nord coal mine, roof of 0m60 vein. Horizon: Siliciclastic strata, Upper Carboniferous.

Referred material: FMNH PE 153; FMNH PE 154. Preservation: Impressions. Repository: FMNH. Locality: USA, Illinois, Mazon Creek. Horizon: Francis Creek Shale, Upper Carboniferous, Moscovian.

Referred material: No stated. Preservation: Impression. Repository: No stated. Locality: Kazakhstan, Karaganda. Horizon: Karaganda Basin, Upper Carboniferous, Pennsylvanian.

Referred material: Unknown. Preservation: Impression (fragmentary). Repository: Unknown. Locality: France, Burgundy, Autun. Horizon: Autunian strata, Lower Permian, Asselian.

Referred material: No stated. Preservation: Ichnite. Repository: No stated. Locality: Canada, Nova Scotia. Horizon: Cumberland Group, Joggins Formation, Upper Carboniferous.

Referred material: HM X. 1041; HM X. 1042. Preservation: Ichnites. Repository: HM. Locality: UK, Isle of Arran. Horizon: Limestone Coal Formation, Upper Carboniferous, Serpukhovian.

Referred material: No stated. Preservation: Ichnite. Repository: No stated. Locality: Canada, Gardner Creek, New Brunswick. Horizon: Tynemouth Creek Formation, Upper Carboniferous, Bashkirian.

Referred material: No stated. Preservation: Ichnite. Repository: No stated. Locality: Canada, Nova Scotia, Cape John; Pugwash to Smith Point. Horizon: Cape John Formation, Upper Carboniferous, Gzhelian; Malagash Formation, Upper Carboniferous, Moscovian.

Referred material: No stated. Preservation: Ichnite. Repository: No stated. Locality: Germany, Flechtingen Volcanic Complex. Horizon: Eiche Member, Flechtingen Formation, Upper Carboniferous, Gzhelian.

Referred material: No stated. Preservation: Ichnite. Repository: No stated. Locality: UK, Fife, Crail to St Andrews. Horizon: Anstruther Formation, Lower Carboniferous, Visean.

Referred material: MNHN.F.SOT002122a. Preservation: Compression. Repository: MNHN. Locality: France, Burgundy, Montceau-les-Mines. Horizon: Upper Carboniferous.

Referred material: LL11219; LL11165; LL11166; LL11167; LL11176. Preservation: Compressions. Repository: MM. Locality: UK, Bickershaw, Lancashire. Horizon: Pennine Lower Coal Measures Formation, Upper Carboniferous, Langsettian.

Referred material: No stated. Two specimens. No stated. Preservation: Impressions. Repository: No stated. Locality: Spain, Ciñera-Matallana, El León; Carrocera, El León; Ciudad Real, Puertollano. Horizon: Pastora Formation, Upper Carboniferous, Stephanian; La Magdalena Coalfield, Upper Carboniferous, Gzhelian; Upper Carboniferous, Stephanian C. Referred material: No stated. Preservation: Ichnite. Repository: MU. Locality: USA, Kentucky, Boyd County. Horizon: Conemaugh Formation, Upper Carboniferous, Gzhelian.

Referred material: No stated. Preservation: Ichnite. Repository: No stated. Locality: Germany, Saxony, Döhlen Basin. Horizon: Döhlen Formation, Lower Permian.

Referred material: No stated. Preservation: Ichnite. Repository: No stated. Locality: France, Burgundy, Montceau-les-Mines. Horizon: Montceau Formation, Upper Carboniferous, Gzhelian-Asselian.

Referred material: No stated. Preservation: Impression. Repository: No stated. Locality: Germany, Thuringia, Manebach. Horizon: Manebach Formation, Lower Permian, Asselian.

Referred material: BMB 014847; BRSMG Cd4053; BRSMG Cd4054; BRSMG Cd4055; BRSMG Cd4056; BRSMG Cd4058. Preservation: Impressions. Repository: BMB; BRSMG. Locality: UK, England, Writhlington Geological Nature Reserve. Horizon: Farrington Formation, Upper Carboniferous.

Referred material: No stated. Preservation: Ichnite. Repository: KUMIP. Locality: USA, Kansas, Waverly. Horizon: Stull Shale Member, Kanwaka Formation, Upper Carboniferous, Gzhelian.

Referred material: No stated. Preservation: Ichnite. Repository: No stated. Locality: Canada, Nova Scotia, Joggins. Horizon: Little River Formation and Joggins Formation, Upper Carboniferous.

Referred material: NMMNH P-45287. Preservation: Ichnite. Repository: NMMNH. Locality: USA, New Mexico, El Cobre Canyon. Horizon: Cutler Group, El Cobre Canyon Formation, Upper Carboniferous, Upper Pennsylvanian.

Referred material: No stated. Preservation: Ichnite. Repository: No stated. Locality: Canada, Nova Scotia. Horizon: Cumberland Group, Joggins Formation, Upper Carboniferous.

Referred material: NHM uncat; NBMG 3315. Preservation: Ichnite. Repository: NHM; NBMG. Locality: Canada, New Brunswick, Saint John. Horizon: Lancaster Formation, Upper Carboniferous, Bashkirian.

Referred material: No stated. Preservation: Impression. Repository: NMP. Locality: Czech Republic, Pilsen, Nýřany. Horizon: Kladno Formation, Upper Carboniferous.

Referred material: No stated. Preservation: Ichnite. Repository: No stated. Locality: Kazakhstan, Zhezkazgan. Horizon: Zhezkazgan Group, Upper Carboniferous, Pennsylvanian.

Referred material: MMRB2010.06.16/527/0.00/a-b; MMRB 2010.07.25/564/0.00. Preservation: Impressions.

Repository: MMRB. Locality: Poland, Lower Silesia, Nowa Ruda mine. Horizon: Shale strata, Upper Carboniferous. Referred material: TA0884. Preservation: Impression. Repository: MfNC. Locality: Germany, Saxony, Chemnitz. Horizon: Leukersdorf Formation, Lower Permian, Sakmarian.

Referred material: USNM 43579; USNM 547011. Preservation: Impressions. Repository: USNM. Locality: USA, Utah, Lime Ridge. Horizon: Halgaito Formation, Lower Permian, Asselian.

Referred material: No stated. Preservation: Ichnite. Repository: SSM; YPM-IP. Locality: USA, Massachusetts, Plainville. Horizon: Rhode Island Formation, Upper Carboniferous, Moscovian.

Referred material: No stated. Preservation: Impressions (fragmentary). Repository: PPC. Locality: Poland, KWK Boleslaw, Przygorze; Czech Republic, Důl Staríč ̌, Chlebovice; Czech Republic, Důl Austria, Zbůch. Horizon: Lower Silesian Basin, Upper Carboniferous, Serpukhovian; Upper Silesian Basin, Upper Carboniferous, Serpukhovian; Plzeň Basin, Upper Carboniferous, Moscovian.

Referred material: No stated. Preservation: Ichnite. Repository: No stated. Locality: UK, Scotland, Fife. Horizon: Strathclyde Group, Pittenweem Formation, Lower Carboniferous, Visean.

Referred material: No stated. Preservation: Ichnite. Repository: No stated. Locality: Ukraine, Donets Basin, Makedonovka. Horizon: Mospinka Formation, Middle Carboniferous, Bashkirian.

Referred material: GR1; GR2; GR3; GR4; GR5; GR6; GR7. Preservation: Ichnites. Repository: GR1-GR6 (in situ); ML. Locality: France, Hérault, Graissessac. Horizon: Graissessac Formation, Upper Carboniferous, Gzhelian.

Referred material: CAMSM X.50355. Preservation: Impression. Repository: CAMSM. Locality: UK, England, Northumberland. Horizon: Stainmore Formation, Upper Carboniferous.

Comments: Salter (1863) recorded *Eurypterus pulicaris* from the Upper Carboniferous Lancaster Formation in Canada. But Falcon-Lang and Miller (2007) consider this fossil instead a representative of *Arthropleura*. Davies *et al*. (2021) recorded the largest fossil specimen of *Arthropleura* from the Upper Carboniferous, Stainmore Formation in UK; they mention two other unpublished records: the ichnite NBMG15084 from the Upper Carboniferous Grand Anse Formation in Canada and another from the Upper Carboniferous Limestone Coal Formation in Glasgow, UK.

**(13) †*Arhtropleura armata* (Jordan & Meyer, 1854)**

*Arthropleura armata* Jordan & Meyer, 1854: 1-15; Goldenberg, 1873: 21, pl 1, Fig.2; Kliver, 1883: 262, pl 36, Fig.2; Woodward, 1907: 547; Vernon, 1912: 634, pl 60, Fig.11; Andrée, 1913: 302, Fig.4; Pruvost, 1919: 76, pl 25, Figs 7-9, text-fig 19; Guthörl, 1934: 179, pl 24, Fig.2; Castro, 1997: 20, Lám I, Fig.3; Proctor, 1998: 94, Fig.1; Štamberg & Zajíc, 2008: 81, Fig.72; Pillola & Zoboli, 2021: 51, Fig.3.

*Halonia irregularis* Geinitz, 1855: 38, pl 4, Fig.5.

*Arthropleura affinis* Goldenberg, 1873: 22, pls 1, 3, Fig.12.

*Arthropleura zeilleri* Boule, 1893: 619-638.

Referred material: No stated. Preservation: Impression. Repository: NHV. Locality: Germany, Jaegersfreude mine. Horizon: Upper Saarbrucker layer Formation, Upper Carboniferous.

Referred material: No stated. Preservation: Impression. Repository: BSC. Locality: Germany, Saarland. Horizon: Lower Saarbrucker layer Formation, Upper Carboniferous.

Referred material: No stated. Preservation: Compression. Repository: BRSMG. Locality: UK, England, Writhlington Geological Nature Reserve. Horizon: Farrington Formation, Upper Carboniferous.

Referred material: No stated. Preservation: Impression. Repository: No stated. Locality: UK, Warwickshire, Baxterley. Horizon: Pennine Middle Coal Measures Formation, Upper Carboniferous, Bashkirian.

Referred material: I. 4022. Preservation: Impression (fragmentary). Repository: BMNH. Locality: UK, Fife, Leven. Horizon: Scottish Upper Coal Measures Formation, Upper Carboniferous, Moscovian.

Referred material: n° 1875; n° 1878; n° 1879; n° 1881. Preservation: Impressions. Repository: MHN. Locality: France, Nord-Pas-de-Calais, Anzin, Cuvinot pit, Boulangère vein (n° 1875), Ledoux pit, 9-Paumes vein (n° 1878, n° 1879), Vendin-lez-Béthune, old Annezin pit 2, assise A2 (n° 1881). Horizon: Shale strata, Upper Carboniferous.

Referred material: No stated. Preservation: Compression. Repository: No stated. Locality: Germany, Saarland. Horizon: Sulzbach Formation, Upper Carboniferous, Moscovian.

Referred material: No stated. Preservation: Impression. Repository: HM. Locality: Spain, Ciñera-Matallana, El León, El Oro mine. Horizon: San José Formation, Upper Carboniferous, Kasimovian.

Referred material: Me 76. Preservation: Impression. Repository: NMP. Locality: Czech Republic, Pilsen, Nýřany. Horizon: Kladno Formation, Upper Carboniferous.

Referred material: PAS-GLP 0113. Preservation: Compression. Repository: PAS. Locality: Italy, Sardinia. Horizon: San Giorgio Formation, Upper Carboniferous.

**(14) †Arthropleura britannica Andree, 1913**

*Arthropleura britannica* Andree, 1913: 302, pl 23, Fig.2; Pruvost, 1930: 174, pl 9, Figs 6.

Referred material: I.G. 3.598. Preservation: Impression. Repository: IRSNB. Locality: Belgium, Flénu. Horizon: Flénu Formation, Upper Carboniferous.

**(15) †Arthropleura cristata Richarson, 1959**

*Arthropleura cristata* Richardson, 1959: 79, Figs 42-43; Hannibal, 1997: 3, Figs 3-5.

Referred material: Holotype: FMNH PE 5262. CaMNH 33853; YPM-PU 88076; FMNH PE 26148. Preservation: Impressions. Repository: FMNH. CaMNH; YPM. Locality: USA, Illinois, Mazon Creek; USA, Pennsylvania, Cannelton. Horizon: Francis Creek Shale, Upper Carboniferous, Moscovian; Kittaning Formation, Upper Carboniferous, Moscovian.

**(16) †Arthropleura fayoli Boule, 1893**

*Arthropleura fayoli* Boule, 1893: 636, pl LV.

Referred material: Syntype: MNHN.F.A31045. Preservation: Impression. Repository: MNHN. Locality: France, Auvergne, Allier, Commentry. Horizon: Stephanian strata, Upper Carboniferous, Gzhelian.

**(17) †Arthropleura maillieuxi Pruvost, 1930**

***Arthropleura maillieuxi*** Pruvost, 1930: 174, pl 10, Figs 1-3.

Referred material: I. G., n 8631. Preservation: Impression. Repository: IRSNB. Locality: Belgium, Mariemont coal mine, Placard pit, roof of La Hestre vein. Horizon: Shale strata, Upper Carboniferous.

**(18) †Arthropleura mammata (Salter, 1863)**

*Eurypterus* (*Arthropleura*) *mammatus* Salter, 1863: 85, Figs 1-7.

*Arthropleura mammata*: Pruvost, 1919: 74, pl 25, Fig.6; text-fig 18; Pruvost, 1930: 172, pl 9, Figs 1-5.

Referred material: No stated. Preservation: Impression Repository: No stated. Locality: UK, Manchester, Pendleton Colliery. Horizon: Shale strata, Upper Carboniferous.

Referred material: No.1880. Preservation: Impression. Repository: MHN. Locality: France, Nord-Pas-de-Calais, Dechy pit, Saint-Nicolas vein, Aniche mines. Horizon: Shale strata, Upper Carboniferous.

Referred material: I.G. 8631; I. G. 8701. Preservation: Impressions. Repository: IRSNB. Locality: Belgium, Mariemont coal mine, Sainte-Henriette pit, roof of Veine-aux-Laies (I. G. 8631), Produits colliery, Flénu, pit 25, roof of Toriore vein (I. G. 8701). Horizon: Shale strata, Upper Carboniferous.

**(19) †Arthropleura moyseyi Calman, 1914**

*Arthropleura moyseyi* Calman, 1914: 541, pl XXXVIII.

Referred material: Holotype: No. 34. Preservation: Impression. Repository: No stated. Locality: UK, Derbyshir, Shipley. Horizon: Pennine Middle Coal Measures Formation, Upper Carboniferous, Bashkirian.

Comments: Hahn *et al*. (1986) stated that *A. monyseyi* is a synonymy of *A. armata*.

**(20) †*Arthropleura punctata* Goldenberg, 1873**

*Arthropleura punctata* Goldenberg, 1873: 22, pl 1, Fig.14.

Referred material: No stated. Preservation: Impression. Repository: NHV. Locality: Germany, Saarbrücken, Friedrichstal railway cut. Horizon: No stated, Upper Carboniferous.

**†Order Eoarthropleurida Shear & Selden, 1995**

**†Family Eoarthropleuridae Størmer, 1976**

**†Genus *Eoarthropleura* Størmer, 1976**

**(21) †Eoarthropleura devonica Størmer, 1976**

*Eoarthropleura devonica* Størmer, 1976: 91, pls 1-4, text-figs 1-4, 6-24, 39-41.

Referred material: No stated. Preservation: Impression. Repository: No stated. Locality: Germany, Alken #3 mine. Horizon: Nellen Koepfchen Beds, Lower Devonian.

**(22) †*Eoarthropleura hueberi* Kjellsvig-Waering, 1986**

*Eoarthropleura hueberi* Kjellesvig-Waering, 1986: 126, pls 5-8, text-fig 50.

Referred material: Holotype: USNM 252629; Paratype: USNM 252630; accession USNM 251091. Preservation:

Compressions. Repository: USNM. Locality: USA, New York, Schoharie County County, South Mountain. Horizon: Onteora Red Beds Formation, Upper Devonian.

**(23) †Eoarthropleura ludfordensis Shear & Selden, 1995**

*Eoarthropleura ludfordensis* Shear & Selden, 1995: 352, Figs 1-2, 351-353.

Referred material: Holotype: K25082; Paratype: K25088. Preservation: Impressions. Repository: NMNI. Locality: UK, England. Horizon: Ludlow Bone Bed, Upper Silurian.

**†Order Microdecemplicida Wilson & Shear, 2000**

**†Family Microdecemplicidae Wilson & Shear, 2000**

**†Genus *Microdecemplex* Wilson & Shear, 2000**

**(24) †*Microdecemplex rolfei* Wilson & Shear, 2000**

*Microdecemplex rolfei* Wilson & Shear, 2000: 352, Figs 1-18.

Referred material: Holotype AMNH 411.15.AR34. Preservation: Impression. Repository: AMNH. Locality: USA, York, Gilboa. Horizon: Panther Mountain Formation, Upper Devonian.

#### 3.2.3 Subclass Chilognatha Latrielle, 1802

**†Order Zosterogrammida Wilson, 2005**

**†Family Zosterogrammidae Wilson, 2005**

**†Genus *Casiogrammus* Wilson, 2005**

**(25) †Casiogrammus ichthyeros Wilson, 2005**

*Casiogrammus icthyeros* Wilson, 2005b: 1104, pl 1, Figs 1-2.

Referred material: Holotype: NMS G.1970.2. Preservation: Impression. Repository: NMS. Locality: UK, South Lanarkshire, Smithy Burn, Hagshaw Hills Inlier. Horizon: Fish Bed Formation, Middle Silurian.

Comments: Preliminarily, Rofle (1980: 550) states that the oldest terrestrial arthropod was a probable myriapod named *Archidesmus loganensis* from the Middle Silurian Fish Bed Formation. According to Rofle (1980: 550, Fig.4A), *A. loganensis* resembles a possible juliform with 20 ornate segments. However, as this fossil is fragmented and poorly preserved, its taxonomic identity must be clarified. Almond (1985: 229-230, Fig.1) placed *A. loganensis* in the Middle Silurian (ca. 438 Ma) before *K. obanensis* from the Upper Silurian (ca. 408 Ma). But, he claimed that *A. loganensis* is not a myriapod. It is more like a form of alga. Wilson & Anderson (2004) also concluded that *A. loganensis* is more of a plant or alga. Wilson (2005b) later described another fragmentary fossil specimen from the Fish Bed Formation: *Casiogrammus ichthyeros*. According to Wilson (2006), *C. ichthyeros* belongs to the order Zosterogrammida. In a stratocladogram, Shear & Edgecombe (2010) placed *C. ichthyeros* in the oldest fossil record of Myriapoda, including other Diplopoda genera: *Albadesmus*, *Cowiedesmus*, and *Pneumodesmus* (Wilson & Anderson, 2004). Edgecombe (2015) included *C. ichthyeros* in a summary taxonomic list of Diplopoda. Selden (2019) stated that *C. ichthyeros* is the oldest known millipede. It is recognized as having a regular series of similar short and broad tergites resembling a Polyzonida. However, Brookfield *et al*. (2020) questioned the identity and provenance of *C. ichthyeros*. According to Brookfield *et al*. (2020), the geologic setting of the Middle Silurian Fish Bed Formation (ca. 430 Ma) is associated with a freshwater environment where fossil fish and aquatic arthropods are typically found. These later authors then concluded that if *C. ichthyeros* is considered a millipede, it could have come from other strata that must have been about the same age as the Upper Silurian Kerrera Sandstone Formation. Thus, *C. ichthyeros* must be as old as *K. obanensis* (Brookfield *et al*., 2020).

**†Genus *Purkynia* Fritsch, 1899**

**(26) †*Purkynia lata* Fritsch, 1899**

*Purkynia lata* Fritsch, 1899: 41, pl 144, Figs 1-3; text-fig 346; Wilson, 2005b: 1106; Štamberg & Zajíc, 2008: 86, Fig.87.

Referred material: Holotype: Me250. Preservation: Compression. Repository: NMP. Locality: Czech Republic, Pilsen, Nýřany. Horizon: Kladno Formation, Upper Carboniferous.

**†Genus *Zosterogrammus* Wilson, 2005**

**(27) †Zosterogrammus stichostethus Wilson, 2005**

*Zosterogrammus stichostethus* Wilson, 2005b: 1103-1104, Figs 1-2.

Referred material: Holotype: YPM 204068; Paratypes: YPM 204067 and YPM 19819; additional material: YPM 204066. Preservation: Impressions. Repository: YPM. Locality: USA, Illinois, Grundy County, Morris, Mazon Creek. Horizon: Carbondale Formation, Upper Carboniferous.

**Infraclass Pentazonia Brandt, 1833**

**†Order Amynilypedida Hoffman, 1969**

**†Family Amynilyspedidae Hoffman, 1969**

**†Genus *Amynilyspes* Scudder, 1882**

*Amynilyspes* sp.: Scudder, 1895: 59, pl IV., Figs 1-2; Racheboeuf *et al*., 2004: 222.

Referred material: No stated. Preservation: Compression. Repository: BMNH. Locality: Canada, Nova Scotia. Horizon: Joggins Formation, Upper Carboniferous.

Referred material: BIRUG 4267. Preservation: Impression. Repository: BIRUG. Locality: UK, England, Deepfields.

Horizon: Brooch Formation, Upper Carboniferous.

**(28) †Amynilyspes crescens Fritsch, 1899**

*Amynilyspes crescens* Fritsch, 1899: 35, pl 146, Fig.1; Štamberg & Zajíc, 2008: 84.

Referred material: Holotype: A 76. Preservation: Compression. Repository: NMP. Locality: Czech Republic, Pilsen, Nýřany. Horizon: Kladno Formation, Upper Carboniferous.

**(29) †Amynilyspes fatimae Racheboeuf et al., 2004**

*Amynilyspes fatimae* Racheboeuf *et al*., 2004: 224, Figs 2-6.

Referred material: Holotype: MNHN SOT 2134; other material: MNHN SOT 2129 and MNHN SOT 14983. Preservation: Impressions. Repository: MNHN. Locality: France, Montceau-les-Mines. Horizon: No stated, Upper Carboniferous.

**(30) †Amynilyspes springhillensis Copeland, 1957**

*Amynilyspes springhillensis* Copeland, 1957: 52, pl 15, Fig.2.

Referred material: Holotype: GSC 10385. Preservation: Impression. Repository: GSC. Locality: Canada, Nova Scotia, Springhill, GSC 1041 site. Horizon: Cumberland Group, Joggins Formation, Upper Carboniferous.

**(31) †Amynilyspes typicus Fritsch, 1899**

*Amynilispes typicus* Fritsch, 1899: 34-35, Fig.340, pl 145, Figs 1-2, pl 147, Figs 1-3; Hoffman, 1969: 586, Fig.370/1; Racheboeuf,

Hannibal & Vannier, 2004: 222; Štamberg & Zajíc, 2008: 84, Fig.82.

Referred material: M 1072; M 1037. Compressions. Repository: NMP. Locality: Czech Republic, Pilsen, Nýřany.

Horizon: Kladno Formation, Upper Carboniferous.

**(32) †Amynilyspes wortheni Scudder, 1882**

*Amynilyspes wortheni* Scudder, 1882: 178, pl 13, Figs 1-4, 9; Hannibal & Feldmann, 1981: 735, pl 1, Figs 1-6, pl 2, Figs 1-6, text-figs 3-6, 9.

Referred material: No stated. Preservation: Compression. Repository: FMNH. Locality: USA, Illinois, Grundy County, Morris, Mazon Creek. Horizon: Carbondale Formation, Upper Carboniferous.

**†Family Sphaerherpestidae Fritsch, 1899**

**†Genus *Glomeropsis* Fritsch, 1895**

**(33) †*Glomeropsis crassa* Fristch, 1899**

*Glomeropsis crassa* Fritsch, 1899: 40, pl 150, Figs 2-3; Štamberg & Zajíc, 2008: 85, Fig.84.

Referred material: Holotype: Me 3. Preservation: Compression. Repository: NMP. Locality: Czech Republic, Pilsen, Nýřany. Horizon: Kladno Formation, Upper Carboniferous.

**(34) †*Glomeropsis magna* Fritsch, 1899**

*Glomeropsis magna* Fritsch, 1899: 40, pl 152, Figs 1-5, text-fig 344; Štamberg & Zajíc, 2008: 85, Fig.85.

Referred material: Holotype: Me 33. Preservation: Compression. Repository: NMP. Locality: Czech Republic, Pilsen, Nýřany. Horizon: Kladno Formation, Upper Carboniferous.

**(35) †Glomeropsis multicarinata Fritsch, 1899**

*Glomeropsis multicarinata* Fritsch, 1899, Fig.358; Fritsch, 1901: 97, pl 165, Figs 1-2; Štamberg & Zajíc, 2008: 85.

Referred material: Holotype: IFG 668. Preservation: Compression. Repository: NMP. Locality: Czech Republic, Pilsen, Nýřany. Horizon: Kladno Formation, Upper Carboniferous.

**(36) †*Glomeropsis ovalis* Fritsch, 1895**

*Glomeropsis ovalis* Fritsch, 1895: 2; Fritsch, 1899: 38, pl 149, Figs 1, 2, 4-7, pl 150, Fig.1, text-figs 343-345; Štamberg & Zajíc, 2008: 85, Fig.83.

Referred material: M 1000; M 1072; and other uncoded material deposited at NMP. Preservation: Compressions.

Repository: NMP. Locality: Czech Republic, Pilsen, Nýřany. Horizon: Kladno Formation, Upper Carboniferous.

**Superorder Limacomorpha Pocock, 1894**

**Order Glomeridesmida Cook, 1895**

**Family Glomeridesmidae Latzel, 1884**

Glomeridesmidae indet: Wesener & Moritz, 2018: 1133, Fig.1A.

Referred material: ZFMK MYR06117: ♂; BuB2423: ♂; BuB3285. Preservation: Amber inclusions. Repository: ZFMK-MYR; BuB. Locality: Myanmar, Kachin, Hukawng Valley, Noije Bum mine. Horizon: No stated, lowermost Upper Cretaceous, Upper Albian/Lower Cenomanian.

**Genus *Glomeridesmus* Gervais, 1844**

*Glomeridesmus* sp.: Santiago-Blay & Poinar, 1992: 366, Figs 2-3.

Referred material: Holotype: GOP DC-3-4: ♀. Preservation: Amber inclusion. Repository: GOP. Locality: Dominican Republic, Santiago, Cordillera Septentriona. Horizon: El Mamey Formation, Lower Miocene.

**Superorder Oniscomorpha Pocock, 1887**

**Order Glomerida Brandt, 1833**

Glomerida indet: Zhang, 2017: 154-155; Wesener & Moritz, 2018: 1133, Fig.1B.

Referred material: No stated. CG-My7276: ♂; CG-BURMA11119; CG-BURMA11127; BuB992: 3 specimens; BuB1821: 3 specimens; BuB2438; BuB2603; BuB2604; BuB2703; BuB2704: 3 specimens; BuB2705; BuB2706; BuB2707; BuB2718; BuB2957; BuB2990; BuB2995; BuB2996; BuB3013; BuB 3014; BuB3015; BuB3016; BuB3053; BuB3058; BuB3257: ♀; BuB3259; ZFMK MYR06116; ZFMK MYR07365; ZFMK MYR07371; ZFMK MYR07372; ZFMK

MYR07376. Preservation: Amber inclusions. Repository: NHML. GC; BuB; ZFMK. Locality: Myanmar, Kachin, Hukawng Valley, Noije Bum mine. Horizon: No stated, lowermost Upper Cretaceous, Upper Albian/Lower Cenomanian.

Comments: After analyzing Zhang’s (2017: 154-155) photomicrographs, Wesener and Moritz (2018) found a misidentification, but consider the record of Glomerida to be valid.

**Family Glomeridellidae Cook, 1896**

**Genus *Glomeridella* Brölemann, 1895**

*Glomeridella* sp.: Ross, 2018: 38.

Referred material: No stated. Preservation: Amber inclusion. Repository: NMS. Locality: Myanmar, Kachin, Hukawng Valley. Horizon: No stated, lowermost Upper Cretaceous, Upper Albian/Lower Cenomanian.

**Family Glomeridae Leach, 1816**

**Genus *Glomeris* Latreille, 1802**

**(37) †Glomeris denticulata Menge, 1854**

*Glomeris denticulata* Menge, 1854: 12.

Referred material: No stated. Preservation: Amber inclusion. Repository: WPM. Locality: Baltic. Horizon: Prussian Formation, Upper Eocene.

**Genus *Hyleoglomeris* Verhoeff, 1910**

**(38) †Hyleoglomeris groehni Wesener, 2019**

*Hyleoglomeris groehni* Wesener, 2019: 41, Figs 1-3.

Referred material: Holotype: GPIH 4931: ♂; #582: ♀. Preservation: Amber inclusions. Repository: GPIH; BNiel.

Locality: Baltic. Horizon: Prussian Formation, Upper Eocene.

**Order Sphaerotheriida Brandt, 1833**

**Family Zephroniidae Gray *In*: Jones, 1843**

Zephroniidae indet: Ross, 2018: 39.

Referred material: No stated. Preservation: Amber inclusion. Repository: NMS. Locality: Myanmar, Kachin, Hukawng Valley, Noije Bum mine. Horizon: No stated, lowermost Upper Cretaceous, Upper Albian/Lower Cenomanian.

**Infraclass Helminthomorpha Pocock, 1887**

**Superorder *Incertae sedis***

**Order Incertae sedis**

**Family Incertae sedis**

**Genus *Archicambala* Cook, 1895**

**(39) Archicambala dawsoni (Scudder, 1868)**

*Xylobius dawsoni* Scudder, 1868; Scudder, 1895: 61, pl 5, Fig.3.

*Archicambala dawsoni*: Cook, 1895: 6.

Referred material: No stated. Preservation: Compression. Repository: BMNH. Locality: Canada, Nova Scotia, Coal Mine Point; Divison 4. Horizon: Joggins Formation, Upper Carboniferous.

**†Genus *Dolesea* Hannibal & May, 2020**

**(40) †*Dolesea subtila* Hannibal & May, 2020**

*Dolesea subtila* Hannibal & May, 2020: 591, Figs 1-2.

Referred material: Holotype: OU 12152; Paratype: OU 12153. Preservation: Impressions. Repository: OU. Locality: USA, Oklahoma, Comanche County, Richards Spur site. Horizon: Fort Sill fissures, Cisuralian Series, Lower Permian.

**†Genus *Sinosoma* Huang & Hannibal, 2018**

**(41) †*Sinosoma luopingense* Huang & Hannibal, 2018**

*Sinosoma luopingense* Huang & Hannibal, 2018: 3, Figs 2-3.

Referred material: Holotype: LPI-61593. Preservation: Compression. Repository: CDCGS. Locality: China, Yunnan, Luoping County, Luoping biota. Horizon: Member II of the Guanling Formation, Anisian, Middle Triassic.

**†Family Archiulidae Scudder, 1873**

**†Genus *Archiulus* Scudder, 1868**

**(42) †*Archiulus brassi* (Dohrn, 1868)**

*Julus brassi* Dohrn, 1868: 335: Taf. VI. Fig.2a u. b; Goldenberg, 1877: 33.

*Palaeojulus brassi*: Werveke, 1906 (in McClennen *et al*., 2017).

*Archiulus brassi*: Guthörl, 1934 (in McClennen *et al*., 2017).

Referred material: Holotype: Mus. F. Naturk. Berlín, 7. Preservation: Impression. Repository: MfN. Locality: Germany, Rhineland-Palatinate. Horizon: Saarlouis Lebach ironstone nodules, Lower Permian.

**(43) †Archiulus euphoberioides Scudder, 1895**

*Archiulus euphoberioides* Scudder, 1895: 59, pl. IV, Figs 5-6.

Referred material: No stated. Preservation: Impression. Repository: BMNH. Locality: Canada, Nova Scotia, Coal Mine Point; Divison 4. Horizon: Cumberland Group, Joggins Formation, Upper Carboniferous.

**(44) †Archiulus glomeratus Sccuder, 1890**

*Archiulus glomeratus* Scudder, 1890: 436, pl 37, Figs 2-3.

Referred material: Holotype: USNM PAL37993. Preservation: Compression. Repository: USNM. Locality: USA, Illinois, Grundy County, Morris, Mazon Creek. Horizon: Carbondale Formation, Upper Carboniferous.

**(45) †*Archiulus lyelli* Scudder, 1895**

*Archiulus lyelli* Scudder, 1895: 60, pl 4, Figs 3, 7.

Referred material: No stated. Preservation: Impression. Repository: BMNH. Locality: Canada, Nova Scotia, GSC 1041, Springhill. Horizon: Cumberland Group, Joggins Formation, Upper Carboniferous.

**(46) †Archiulus xylobioides Sccuder, 1868**

*Archiulus xylobioides* Scudder, 1868: 496, Fig.151b; Scudder, 1895: 59; Copeland, 1957: 53.

Referred material: No stated. Preservation: Impression. Repository: BMNH. Locality: Canada, Nova Scotia. Horizon: Cumberland Group, Joggins Formation, Upper Carboniferous.

**†Family Oklahomasomatidae Hannibal & May, 2020**

**†Genus *Oklahomasoma* Hannibal & May, 2020**

**(47) †Oklahomasoma richardsspurense Hannibal & May, 2020**

*Oklahomasoma richardsspurense* Hannibal & May, 2020: 590, Figs 2-3.

Referred material: Holotype: OU 44526. Preservation: Impression. Repository: OU. Locality: USA, Oklahoma, Comanche County, Richards Spur. Horizon: Fort Sill fissures, Cisuralian Series, Lower Permian.

**†Superorder Archipolypoda Scudder, 1882**

**Order *Incertae sedis***

**Family Incertae sedis**

**†Genus *Albadesmus* Wilson & Anderson, 2004**

**(48) †*Albadesmus almondi* Wilson & Anderson, 2004**

*Albadesmus almond* Wilson & Anderson, 2004: 174, Figs 8.1-8.3.

Referred material: Holotype: AMS F.64847; other material: BGS GSE 14780. Preservation: Impressions. Repository: AMS; BGS. Locality: UK, Scotland, Cowie Harbour, Stonehaven. Horizon: Cowie Formation, Upper Silurian.

**†Genus *Anaxeodesmus* Wilson, 2005**

**(49) †Anaxeodesmus diambonotus Wilson, 2005**

*Anaxeodesmus diambonotus* Wilson, 2005a: 1099, Figs 1-2.

Referred material: Holotype: NHM In.18370. Preservation: Impression. Repository: NHM. Locality: UK, Staffordshire, Coseley. Horizon: Foot Ironstone, Upper Carboniferous.

**†Genus *Anthracodesmus* Peach, 1899**

**(50) †Anthracodesmus macconochiei Peach, 1899**

*Anthracodesmus macconochiei* Peach, 1899: 121, pl 3, Figs 3; Wilson & Anderson, 2004: 177, Figs 10.4-10.5.

Referred material: Holotype: BGS 2176. Preservation: Impression. Repository: BGS. Locality: UK, Scotland, Coldstream, Lennel Braes. Horizon: No stated, Lower Carboniferous.

**†Genus *Kampecaris* Page, 1856**

**(51) †Kampecaris dinmorensis Clarke, 1951**

*Kampecaris dinmorensis* Clarke, 1951 (in McClennen *et al*., 2017); Almond, 1985: 16, Fig.1.

Referred material: No stated. Preservation: Compression. Repository: No stated. Locality: UK, England, Herefordshire, Dinmore Hill. Horizon: No stated, Lower Devonian.

**(52) †Kampecaris forfarensis Peach, 1882**

*Kampecaris forfarensis* Peach, 1882: 179, pl 2, Figs 1a-g; Almond, 1985: 231, pl 1, Figs 5, 7.

Referred material: No stated. Preservation: Compression, Repository: BMNH. Locality: UK, Scotland, Angus, Mirestone Quarry. Horizon: Old Red Sandstone, Carmyllie Group, Lower Devonian.

**(53) †Kampecaris obanesis Peach, 1889**

*Kampecaris obanesis* Peach, 1899: 122, pl IV, Fig.2.

Referred material: Holotype: Preservation: Compression. No stated. Repository: IGCSE. Locality: UK, Scotland.

Horizon: Kerrera Sandstone Formation, Upper Silurian.

Comments: Brookfield *et al*. (2020) claimed that it is the oldest terrestrial animal fossil.

**†Genus *Palaeodesmus* Brade-Birks, 1923**

**(54) †*Palaeodesmus tuberculata* (Brade-Birks, 1923)**

*Kampecaris tuberculata* Brade-Birks, 1923: 277, pl 33; Almond, 1985: 233, pl 1, Fig.8.

*Palaeodesmus tuberculata*: Wilson & Anderson, 2004: 177, Figs 10.1-10.3.

Referred material: Holotype: NHM In. 23803. Preservation: Compression. Repository: NHM. Locality: UK, Scotland, Dunure. Horizon: No stated, Lower Devonian, Emsian.

**†Genus *Pneumodesmus* Wilson & Anderson, 2004**

**(55) †*Pneumodesmus newmani* Wilson & Anderson, 2004**

*Pneumodesmus newmani* Wilson & Anderson, 2004: 174, Figs 9.1-9.3.

Referred material: Holotype: NMS G.2001.109.1; other material: NHM In. 43574. Repository: Impressions. Repository: NMS. Locality: UK, Scotland, Cowie Harbour, Stonehaven. Horizon: Cowie Formation, Upper Silurian.

**†Family Woodesmidae Ross *et al*., 2008**

**†Genus *Woodesmus* Ross *et al*., 2018**

**(56) †Woodesmus sheari Ross et al., 2018**

*Woodesmus sheari* Ross *et al*., 2018: 4, Figs 6-7.

Referred material: Holotype: NMS G.2012.39.10; Paratype NMS G. 2016.23.1. Preservation: Compressions.

Repository: NMS. Locality: UK, Scotland, Burnmouth. Horizon: Ballagan Formation, Lower Carboniferous.

**†Order Archidesmida Wilson & Anderson, 2004**

**†Family Archidesmidae Scudder, 1885**

**†Genus *Archidesmus* Peach, 1882**

**(57) †Archidesmus macnicoli Peach, 1882**

*Archidesmus macnicoli* Peach, 1882: 182, pl 2, Fig.2; Almond, 1985: 231, pl 1, Fig.4; Wilson & Anderson, 2004: 172, Figs 2-6.

Referred material: AUGD 12303a, b; AUGD 12302a, b; NMS G. 1964.31.15. Syntypes: NMS G. 1953.7.1, and NMS G. 1891.92.72. Preservation: Impressions. Repository: NMS. Locality: UK, Scotland, Tillywhandland Quarry. Horizon: Dundee Formation, Upper Silurian.

**†Family Zanclodesmidae Wilson *et al*., 2005**

**†Genus *Orsadesmus* Wilson *et al*., 2005**

**(58) †Orsadesmus rubecollus Wilson et al., 2005**

*Orsadesmus rubecollus* Wilson *et al*., 2005a: 743, Figs 2-3.

Referred material: Holotype: ANSP 80134; Paratypes: ANSP 80133; ANSP 80135; ANSP 80136. Preservation: Impressions. Repository: ANSP. Locality: USA, Pennsylvania, Clinton County, Red Hill. Horizon: Catskill Formation, Upper Devonian.

**†Genus *Zanclodesmus* Wilson *et al*., 2005**

**(59) †Zanclodesmus willetti Wilson et al., 2005**

*Zanclodesmus willetti* Wilson *et al*., 2005a: 743, Fig.4.

Referred material: Holotype: MHNM 27-40a. Preservation: Impression. Repository: MHNM. Locality: Canada, Quebec, Miguasha, bed 220. Horizon: Escuminac Formation, Upper Devonian.

**†Order Cowiedesmida Wilson & Anderson, 2004**

**†Family Cowiedesmidae Wilson & Anderson, 2004**

**†Genus *Cowiedesmus* Wilson & Anderson, 2004**

**(60) †*Cowiedesmus eroticopodus* Wilson & Anderson, 2004**

*Cowiedesmus eroticopodus* Wilson & Anderson, 2004: 174, Fig.7.

Referred material: Holotype: AMS F.64845. Preservation: Impression. Repository: AMS. Locality: UK, Scotland, Cowie Harbour, Stonehaven. Horizon: Cowie Formation, Upper Silurian.

**†Order Euphoberiida Hoffman, 1969**

**†Family Euphoberiidae Scudder, 1882**

**†Genus *Acantherpestes* Meek & Worthen, 1868**

**(61) †Acantherpestes foveolatus Fritsch, 1899**

*Acantherpestes foveolatus* Fritsch, 1899: 21, pl 137, Figs 1-6, text-figs 329-330; Štamberg & Zajíc, 2008: 83, Fig.77.

Referred material: Holotype: Me 4. Preservation: Compression. Repository: CGH. Locality: Czech Republic, Pilsen, Nýřany. Horizon: Kladno Formation, Upper Carboniferous.

**(62) †Acantherpestes gigas Fritsch, 1895**

*Acantherpestes gigas* Fritsch, 1895: 2; Fritsch, 1899: 16, pl 134, Figs 1-7, text-figs 323, 324; Štamberg & Zajíc, 2008: 82.

Referred material: Holotype: Me 5. Preservation: Compression. Repository: CGH. Locality: Czech Republic, Pilsen, Nýřany. Horizon: Kladno Formation, Upper Carboniferous.

**(63) †Acantherpestes inequalis Scudder, 1890**

*Acantherpestes inequalis* Scudder, 1890: 424, pl 33, Figs 2, 4.

Referred material: Holotype: USNM PAL38042. Preservation: Compression. Repository: USNM. Locality: United States, Illinois, Grundy County, Morris, Mazon Creek. Horizon: Carbondale Formation, Upper Carboniferous.

**(64) †*Acantherpestes major* (Meek & Worthen, 1868)**

*Euphoberia major* Meek & Worthen, 1868: 26.

*Acantherpestes major*: Scudder, 1882: 150, pl 10, pl 11, Figs 1-4, 6-8, 10-11; Scudder, 1890: 424.

Referred material: Uncoded holotype. Preservation: Impression. Repository: NHMW. Locality: USA, Illinois, Grundy County, Morris, Mazon Creek. Horizon: Carbondale Formation, Upper Carboniferous.

**(65) †Acantherpestes ornatus Fritsch, 1899**

*Acantherpestes ornatus* Fritsch, 1899: 19, pl 136, Figs 1-9, text-figs 327-328; Štamberg & Zajíc, 2008: 83.

Referred material: Holotype: No. 58. Preservation: Compression. Repository: ZCM. Locality: Czech Republic, Pilsen, Nýřany. Kladno Formation, Upper Carboniferous.

**(66) †Acantherpestes vicinus Fritsch, 1899**

*Acantherpestes vicinus* Fritsch, 1899: 18, pl 135, Figs 1-6, text-figs 325-326.

Referred material: Holotype: Me 1; Paratype: Me 2. Preservation: Compressions. Repository: CGH. Locality: Czech Republic, Pilsen, Nýřany. Horizon: Kladno Formation, Upper Carboniferous.

**†Genus *Euphoberia* Meek & Worthen, 1868**

**(67) †*Euphoberia absens* Fritsch, 1899**

*Euphoberia absens* Fritsch, 1899: 25, pl 135, Fig. 9, text-fig 334.

Referred material: Holotype: Me 24. Preservation: Compression. Repository: CGH. Locality: Czech Republic, Pilsen, Nýřany. Horizon: Kladno Formation, Upper Carboniferous.

**(68) †Euphoberia anguilla Scudder, 1882**

*Euphoberia anguilla* Scudder, 1882: 177, pl 12, Fig. 20; Scudder, 1890: 435, pl 36, Fig.3.

Referred material: Holotype: USNM PAL263917. Preservation: Compression. Repository: USNM. Locality: USA, Illinois, Grundy County, Morris, Mazon Creek. Horizon: Carbondale Formation, Upper Carboniferous.

**(69) †*Euphoberia armigera* Meek & Worthen, 1868**

*Euphoberia armigera* Meek & Worthen, 1868: 26; Scudder, 1882: 160, pl 12, Figs 1-3, 5-6, 13, pl 13, Figs 7-8, 10; Scudder, 1890: 427, pl 34, Figs 2, 4, 8, pl 35, Fig.3; Baldwin, 1911: 77, pl 5, Fig.4; Jackson *et al*., 1919: 408, pl 9, Fig.2, text-fig 3.

Referred material: Holotype: PE 45046. Preservation: Impression. Repository: FMNH. Locality: USA, Illinois, Grundy County, Morris, Mazon Creek. Horizon: Carbondale Formation, Upper Carboniferous.

**(70) †*Euphoberia brownii* Woodward, 1871**

*Euphoberia brownii* Woodward, 1871: 102, pl 3, Figs 6a-c; Scudder, 1882: 167, pl 12, Figs 7-8, 21.

Referred material: Holotype: NMS G.1884.46.180. Preservation: Compression. Repository: NMS. Locality: UK, Woodhill Quarry, Kilmaurs. Horizon: No stated, Upper Carboniferous.

**(71) †*Euphoberia carri* Scudder, 1882**

*Euphoberia carri* Scudder, 1882: 171, pl 12, Figs 4, 9-12, 14-19, pl 13, Figs 16, 18; Scudder, 1890: 429.

Referred material: Uncoded holotype. Preservation: Compression. Repository: USNM. Locality: USA, Illinois, Grundy County, Morris, Mazon Creek. Horizon: Carbondale Formation, Upper Carboniferous.

**(72) †Euphoberia cuspidata Scudder, 1890**

*Euphoberia cuspidata* Scudder, 1890: 429, pl 34, Figs 3, 7.

Referred material: Holotype: USNM PAL38017. Preservation: Compression. Repository: USNM. Locality: USA, Illinois, Grundy County, Morris, Mazon Creek. Horizon: Carbondale Formation, Upper Carboniferous.

**(73) †*Euphoberia ferox* (Salter, 1863)**

*Eurypterus* (*Arthropleura*) *ferox* Salter, 1863: 86, Fig.8.

*Acantherpestes brodiei* Scudder, 1882: 156, pl 11, Fig.5.

*Euphoberia ferox*: Scudder, 1882: 157, pl 12, Fig.23; Woodward, 1887: 6, pl 1, Figs 1-13, text-figs 1-2; Pruvost, 1930: 168, pl 7, Fig.3.

Referred material: Uncoded holotype. Preservation: Impression. Repository: BGS. Locality: Belgium, Mariemont coal mine. Horizon: Shale strata, Upper Carboniferous.

**(74) †Euphoberia flabellata Scudder, 1882**

*Euphoberia flabellata* Scudder, 1882: 174, pl 13, Fig.15.

Referred material: Holotype. Repository: Compression. Repository: YPM. Locality: USA, Illinois, Grundy County, Morris, Mazon Creek. Horizon: Carbondale Formation, Upper Carboniferous.

**(75) †*Euphoberia granosa* Scudder, 1882**

*Euphoberia granosa* Scudder, 1882: 168, pl 12, Figs 22, 24-26, pl 13, Fig.13; Scudder, 1890: 427, pl 34, Figs 5-6, pl 36, Fig.2.

Referred material: Holotype. Preservation: Impression. Repository: USNM. Locality: USA, Illinois, Grundy County, Morris, Mazon Creek. Horizon: Carbondale Formation, Upper Carboniferous.

**(76) †*Euphoberia histrix* Fritsch, 1899**

*Euphoberia histrix* Fritsch, 1899: 23, pl 138, Fig.8, text-figs 332-333; Štamberg & Zajíc, 2008: 81, Figs 73-74.

Referred material: Holotype: M 1013. Preservation: Impression. Repository: CGH. Locality: Czech Republic, Pilsen, Nýřany. Horizon: Kladno Formation, Upper Carboniferous.

**(77) †*Euphoberia horrida* Scudder, 1882**

*Euphoberia horrida* Scudder, 1882: 158, pl 13, Figs 11-12, 14.

Referred material: Holotype. Preservation: Compression. Repository: USNM. Locality: USA, Illinois, Grundy County, Morris, Mazon Creek. Horizon: Carbondale Formation, Upper Carboniferous.

**(78) †Euphoberia hystricosa Scudder, 1890**

*Euphoberia hystricosa* Scudder, 1890: 426, pl 33, Figs 1, 3.

Referred material: Holotype: USNM PAL38025. Preservation: Compression. Repository: USNM. Locality: USA, Illinois, Grundy County, Morris, Mazon Creek. Horizon: Carbondale Formation, Upper Carboniferous.

**(79) †*Euphoberia lithanthracis* (Jordan & Meyer, 1854)**

*Chonionotus lithanthracis* Jordan & Meyer, 1854: Taf. I. Fig.19; Goldenberg, 1873: 21, pl 1, Fig.19.

*Euphoberia lithanthracis*: Guthörl, 1934: 21.

Referred material: Holotype: Mus. F. Naturk. Berlín, i 1. Preservation: Impression. Repository: MfN. Locality: Germany, Jaegersfreude mine. Horizon: Upper Saarbrucker layer Formation, Upper Carboniferous.

**(80) †*Euphoberia simplex* Scudder, 1890**

*Euphoberia simplex* Scudder, 1890: 432, pl 35, Figs 2, 6-7.

Referred material: Holotype: USNM PAL38024. Preservation: Compression. Repository: USNM. Locality: USA, Illinois, Grundy County, Morris, Mazon Creek. Horizon: Carbondale Formation, Upper Carboniferous.

**(81) †Euphoberia spinulosa Scudder, 1890**

*Euphoberia spinulosa* Scudder, 1890: 430, pl 34, Fig.1, pl 35, Figs 1, 4-5, pl 36, Figs 7-8.

Referred material: Holotype: USNM PAL37990. Preservation: Compression. Repository: USNM. Locality: USA, Illinois, Grundy County, Morris, Mazon Creek. Horizon: Carbondale Formation, Upper Carboniferous.

**(82) †*Euphoberia tracta* Scudder, 1890**

*Euphoberia tracta* Scudder, 1890: 433, pl 36, Figs 1, 4-6.

Referred material: Holotype: USNM PAL37992. Preservation: Compression. Repository: USNM. Locality: USA, Illinois, Grundy County, Morris, Mazon Creek. Horizon: Carbondale Formation, Upper Carboniferous.

**(83) †*Euphoberia varians* Fritsch, 1899**

*Euphoberia varians* Fritsch, 1899: 23, pl 138, Figs 1-7, text-fig 331; Štamberg & Zajíc, 2008: 82, Fig.75.

Referred material: Holotype: Me 20. Preservation: Compression. Repository: CGH. Locality: Czech Republic, Pilsen, Nýřany. Horizon: Kladno Formation, Upper Carboniferous.

**†Genus Myriacantherpestes Burke, 1979**

*Myriacantherpestes* sp.: Wilson & Anderson, 2004: 171.

Referred material: Holotype: FMNH PE 2464. Preservation: Impression. Repository: FMNH. Locality: UK, England, Coseley. Horizon: No stated, Upper Carboniferous.

**†Order Palaeosomatida Hannibal & Krzeminski, 2005**

**†Family Palaeosomatidae Hannibal & Krzeminski, 2005**

**†Genus *Paleosoma* Jackson *et al*., 1919**

**(84) †*Paleosoma giganteus* (Baldwin, 1911)**

*Acantherpestes giganteus* Baldwin, 1911: 76, pl 4, Fig.1.

*Palaeosoma giganteum*: Jackson *et al*., 1919: 406, pl 9, Fig.1, text-figs 1-2.

Referred material: Holotype: MM L9941. Preservation: Impression. Repository: MM. Locality: UK, England, Sparth Bottoms, Rochdale. Horizon: No stated, Upper Carboniferous.

**(85) †Palaeosoma robustum Jackson et al., 1919**

*Euphoberia robusta* Baldwin, 1911: 77, pl 5, Fig.3.

*Euphoberia woodwardi* Baldwin, 1911: 78, pl 4, Fig.2.

*Palaeosoma robustum* Jackson *et al*., 1919: 409, pl 9, Fig.3; Hannibal & Krzemiński, 2005: 208, Figs 2-10.

Referred material: Holotype: MM L9943. Preservation: Impression. Repository: MM. Locality: UK, England, Sparth Bottoms, Rochdale. Horizon: No stated, Upper Carboniferous.

Referred material: No. 1089. Preservation: Impression. Repository: MPW. Locality: Poland, Jan shaft, Bialy Kamen, Walbrzych. Horizon: Walbrzych Formation, Lower Carboniferous.

**Subterclass Colobognatha Brandt, 1834**

**†Order Pleurojulida Schneider & Werneburg, 1998**

**†Family Pleurojulidae Schneider & Werneburg, 1998**

**†Genus *Pleurojulus* Fritsch, 1899**

**(86) †Pleurojulus biornatus Fritsch, 1899**

*Pleurojulus biornatus* Fritsch, 1899, pl 139, Figs 1-9, pl 143, Fig.9; Wilson & Hannibal, 2005: 1107, Figs 2.5-2.6; Štamberg & Zajíc,

2008: 83, Figs 79-80.

Referred material: Holotype: Me 7. Preservation: Impression. Repository: CGH. Locality: Czech Republic, Pilsen, Nýřany. Horizon: Kladno Formation, Upper Carboniferous.

**(87) †*Pleurojulus levis* Fritsch, 1899**

*Pleurojulus levis* Fritsch, 1899: 28, pl 141, Figs 1-11; Wilson & Hannibal, 2005: 1107, Figs 2.1-2.4; Štamberg & Zajíc, 2008: 84, Fig.81.

*Pleurojulus aculeatus* Fritsch, 1899: 28, pl 141, Figs 12-14.

*Pleurojulus pinguis* Fritsch, 1899: 29, pl 140, Figs 8-9.

Referred material: Holotype: Me 13. Preservation: Compression. Repository: CGH. Locality: Czech Republic, Pilsen, Nýřany. Horizon: Kladno Formation, Upper Carboniferous.

**†Genus *Isojulus* Fritsch, 1899**

**(88) †*Isojulus constans* (Frič, 1875)**

*Julus constans* Frič, 1875: 74.

*Archijulus constans*: Fritsch, 1895: 2.

*Isojulus constans*: Fritsch, 1899: 25, pl 142, Figs 1-3, text-fig 335; Wilson & Hannibal, 2005: 1109, Figs 7-8; Štamberg & Zajíc, 2008:

83, Fig.78.

*Isojulus marginatus* Fritsch, 1899: 26, pl 140, Figs 1-2, pl 142, Figs 9, 16.

*Isojulus setipes* Fritsch, 1899: 26, pl 142, Figs 4-8.

*Pleurojulus longipes* Fritsch, 1899: 28, pl 140, Figs 3-9.

*Pleurojulus falcifer* Fritsch, 1899: 29, pl 140, Fig.10.

Referred material: Holotype: M1040. Preservation: Compression. Repository: CGH. Locality: Czech Republic, Pilsen, Nýřany. Horizon: Kladno Formation, Upper Carboniferous.

**Order Platydesmida Cook, 1895**

Platydesmida indet: Álvarez-Rodríguez *et al*., 2023, Fig.3D (this work).

Referred material: CPAL.123: ♀; CPAL.157: ♀; CPAL.161: ♀; CPAL.207: juvenile. Preservation: Amber inclusions. Repository: CPAL-UAEM. Locality: Mexico, Chiapas, Simojovel, Guadalupe Victoria mine (CPAL.123), Monte Cristo mine (CPAL.157 and CPAL.161), San Antonio mine (CPAL.207). Horizon: Uppermost Simojovel Formation, Lower Miocene.

**Family Andrognathidae Cope, 1869**

Andrognathidae indet: Wesener & Moritz, 2018: 1134, Fig.1E.

Referred material: Wu F3391/Bu/CJW: ♂; BuB1413: ♀; BuB2670: ♂; BuB2991: ♂; BuB3237: ♂; BuB3291: ♀; BuB3307: ♂; BuB3308: ♀. Preservation: Amber inclusions. Repository: Wu; BuB. Locality: Myanmar, Kachin, Hukawng Valley, Noije Bum mine. Horizon: No stated, lowermost Upper Cretaceous, Upper Albian/Lower Cenomanian.

**Genus *Brachycybe* Wood, 1864**

*Brachycybe* sp.: Ross, 2018: 38.

Referred material: No stated. Preservation: Amber inclusion. Repository: NMS. Locality: Myanmar, Kachin, Hukawng Valley. Horizon: No stated, lowermost Upper Cretaceous, Upper Albian/Lower Cenomanian.

**Genus *Andrognathus* Cope, 1869**

**(89) †*Andrognathus burmiticus* Moritz & Wesener, 2019**

*Andrognathus burmiticus* Moritz & Wesener, 2019: 425, Figs 1-4.

Referred material: Holotype: ZFMK-MYR8241: ♂; Paratypes: ZFMK-MYR08689: ♂; ZFMK-MYR08617: ♀; ZFMK-MYR08679: ♀; ZFMK-MYR08709; ♀; ZFMK-MYR-08710: juvenile; additional material: BuB997: ♂; BuB1413:

♂; BuB2991: ♂; BuB3005: ♂; BuB3237: ♂; BuB3308: ♂. Preservation: Amber inclusions. Repository: ZFMK-MYR; BuB. Locality: Myanmar, Kachin, Hukawng Valley, Noije Bum mine. Horizon: No stated, lowermost Upper Cretaceous, Upper Albian/Lower Cenomanian.

**Order Polyzoniida Cook, 1895**

Polyzoniida indet: Wesener & Moritz, 2018: 1134.

Referred material: BuB112: ♀; BuB219: ♀; BuB913: ♀; BuB914: 8 ♀; BuB919: ♀: BuB1031-B; BuB1994; BuB1996; BuB2964: ♀; BuB2965; BuB2980: ♀; BuB3017; BuB3033; BuB3044; Wu F3167/BU/CJW: 3 specimens; Wu F3172/BU/CJW; Wu F3201/Bu/CJW; Wu F3202/Bu/CJW; Wu F3107/Bu/CJW; Wu F3390/Bu/CJW: ♀; Wu

F3395/Bu/CJW; Wu F3400/Bu/CJW; Wu F3401/Bu/CJW: ♀; RO my153: ♀; RO my199: ♀. Preservation: Amber inclusions. Repository: BuB; Wu; RO. Locality: Myanmar, Kachin, Hukawng Valley, Noije Bum mine. Horizon: No stated, lowermost Upper Cretaceous, Upper Albian/Lower Cenomanian.

**Family Polyzoniidae Newport, 1844**

**Genus *Polyzonium* Brandt, 1837**

*Polyzonium* sp.: Hoffman, 1969: 589, Fig.375, 1.

Referred material: No stated. Preservation: Amber inclusion. Repository: No stated. Locality: Baltic region. Horizon: Prussian Formation, Upper Eocene.

Comments: Hoffman (1969: 589) stated that *Polyzonium* sp. is probably *P. germanicum*.

**Family Siphonotidae Cook, 1895**

Siphonotidae indet: Zhang, 2017: 182. Wesener & Moritz, 2018: 1134, Fig.1D.

Referred material: No stated. BuB612; BuB825: ♀; BuB837: ♀; BuB817: ♀; BuB824: ♀; BuB831: ♀; BuB826: ♀; BuB836: ♀; BuB840: ♀; BuB925: ♀; BuB979: ♀; BuB1130: ♀; BuB1034: ♀; BuB1087: ♀; BuB1161: ♀; BuB1162: 3 specimnes; BuB1163: ♀; BuB1164: ♀; BuB1166: ♀; BuB1167: ♀; BuB1834: ♀; BuB1837: two ♀; BuB1853:♀; BuB1854:

♀; BuB1855: ♀; BuB1856: two ♀; BuB1956: ♀; BuB1966: ♀; BuB1972: ♀; BuB1976: ♀; BuB1983: ♀; BuB1984: 2 specimens; BuB1993: ♀; BuB2607: ♀; BuB2608: ♀; BuB2609: ♀; BuB2610: ♀; BuB2611: ♀; BuB2615; BuB2656; BuB2657; BuB3266: ♀; BuB3268: ♀; BUB3270: ♀; BuB3272: two ♀; BuB3273: ♀; BuB3280: juvenile; BuB3281: two ♀; BuB3284: five ♀; ZFMK MYR06122: ♀; ZFMK MYR06124: ♀; ZFMK MYR07374: ♀; ZFMK MYR07381: ♀.

Preservation: Amber inclusions. Repository: NMS. BuB; ZFMK-MYR. Locality: Myanmar, Kachin, Hukawng Valley, Noije Bum mine. Horizon: No stated, lowermost Upper Cretaceous, Upper Albian/Lower Cenomanian.

**Genus *Siphonotus* Brandt, 1837**

*Siphonotus* sp.: Santiago-Blay & Poinar, 1992: 366, Fig.7.

Referred material: Holotype: GOP DC-3-15: juvenile. Preservation: Amber inclusion. Repository: GOP. Locality: Dominican Republic, Santiago, Cordillera Septentrional. Horizon: El Mamey Formation, Lower Miocene.

**Order Siphonophorida Newport, 1844**

Siphonophorida indet: Wesener & Moritz, 2018: 1134, Fig.1C. Álvarez-Rodríguez *et al*., 2023 (this work).

Referred material: BuB823: ♀; BuB977: ♂; BuB982; BuB905: ♂; BuB1835: ♀; BuB1951: ♀; BuB1959: ♂; BuB1966b; BuB1970; BuB1971: ♀; BuB1977: ♀; BuB1978: ♀; BuB1980: ♀; BuB1981: ♂; BuB1991: ♂; BuB2605; BuB2973; BuB3019: ♀; BuB3035: ♂; BuB3036: ♂; BuB3037; BuB343; BuB3045: ♀; BuB3054: ♀; BuB3057.

Preservation: Amber inclusions. Repository: BuB. Locality: Myanmar, Kachin, Hukawng Valley, Noije Bum mine. Horizon: No stated, lowermost Upper Cretaceous, Upper Albian/Lower Cenomanian.

New records: CPAL.118.1; CPAL.118.2: juvenile; CPAL.133: (two parts); CPAL.142; CPAL.162: juvenile; CPAL.163; CPAL.164; CPAL.165: juvenile; CPAL.166; CPAL.175; CPAL.180; CPAL.203: ♀; CPAL.204: juvenile; CPAL.212; CPAL.213; CPAL.218: juvenile; CPAL.182: incomplete. Preservation: Amber inclusions. Repository: CPAL-UAEM. Locality: Mexico, Chiapas, Simojovel, El Pabuchil mine (CPAL.180), Los Pocitos mine (CPAL.118.1, CPAL.118.2, CPAL.133, CPAL.142, CPAL.162, CPAL.163, CPAL.164, CPAL.165, CPAL.166, CPAL.175), Monte Cristo mine (CPAL.203, CPAL.212, CPAL.219), San Antonio mine (CPAL.213); Palenque, Estella de Belén mine (CPAL.204). Horizon: Uppermost Simojovel Formation, Lower Miocene.

**Family Siphonophoridae Newport, 1844**

Siphonophoridae indet: Grimaldi *et al*., 2002: 10. Zhang, 2017: 167-169. Wesener & Moritz, 2018: 1134.

Referred material: 4 specimens. No stated. RO my130: ♀; RO my330; BuB1030: ♀; BuB2243: ♀; BuB644: ♀; BuB73; BuB828; BuB978: ♀; BuB981: ♀; BuB984; BuB986; BuB1143: ♀; BuB1159: ♀; BuB2963: ♀; BuB2973: ♀; BuB2986; BuB2989: ♂; BuB2997; BuB3006: ♂; BuB3007: ♂; BuB3010; BuB3034: ♂; BuB3047; BuB3052: ♀; BuB3239: ♀;

BuB3245; BuB3261: two ♀; BuB3262: ♀; Wu F3149/Bu/CJW; Wu F3393/Bu/CJW. Preservation: Amber inclusions. Repository: AMNH. NMS. RO; BuB; Wu. Locality: Myanmar, Kachin, Hukawng Valley, Noije Bum mine. Horizon: No stated, lowermost Upper Cretaceous, Upper Albian/Lower Cenomanian.

Comments: Zhang (2017: 167-169) preliminarily identified the family Siphonorhinidae. Subsequently, based on the microphotographs published by Zhang (2017: 167-169), the Siphonorhinidae material was assigned to Siphonophoridae (Wesener & Moritz, 2018: 1138).

**Genus *Siphonophora* Brandt, 1837**

*Siphonophora* sp.: Santiago-Blay & Poinar, 1992: 366. Riquelme & Hernández-Patricio, 2018: 638, Figs 1A-C. Referred material: Holotype: GOP DC-3-9A: ♀; two juveniles. Preservation: Amber inclusions. Repository: GOP; PC.

Locality: Dominican Republic, Santiago, Cordillera Septentrional. Horizon: El Mamey Formation, Lower Miocene.

Referred material: CPAL.102: ♂; MALM.21. Preservation: Amber inclusions. Repository: CPAL; MALM. Locality: Mexico, Chiapas, Simojovel, Guadalupe Victoria mine (CPAL.102), La Pimienta (MALM.21). Horizon: Uppermost Simojovel Formation, Lower Miocene.

**(90) †Siphonophora hui Jiang et al., 2019**

*Siphonophora hui* Jiang *et al*., 2019: 35, Figs 1-4; Su *et al*., 2021: 280, Figs 1-7.

Referred material: Holotype: IBGAS PAL 001-1: ♂; Paratype: IBGAS PAL 001-2. NIGP175067: ♂; NIGP175068: ♂; NIGP175069: ♀. Preservation: Amber inclusions. Repository: IBGAS. NIGP: Locality: Myanmar, Kachin, Hukawng Valley, Noije Bum mine. Horizon: No stated, lowermost Upper Cretaceous, Upper Albian/Lower Cenomanian.

**(91) †*Siphonophora hoffmani* Santiago-Blay & Poinar, 1992**

*Siphonophora hoffmani* Santiago-Blay & Poinar, 1992: 368, Figs 18-20.

Referred material: Holotype: GOP DC-3-9: ♂; Paratype: GOP DC-3-7: ♂; Allotype: GOP DC-3-8: ♀; ♂. Preservation: Amber inclusions. Repository: GOP; PC. Locality: Dominican Republic, Santiago, Cordillera Septentrional. Horizon: El Mamey Formation, Lower Miocene.

**(92) †*Siphonophora velezi* Santiago-Blay & Poinar, 1992**

*Siphonophora velezi* Santiago-Blay & Poinar, 1992: 369, Fig. 21.

Referred material: Holotype: ♂ (incomplete); Allotype: ♀. Preservation: Amber inclusions. Repository: PC. Locality: Dominican Republic, Santiago, Cordillera Septentrional. Horizon: El Mamey Formation, Lower Miocene.

**Genus *Siphonocybe* Pocock, 1903**

*Siphonocybe* sp.: Shear, 1981: 54.

Referred material: No stated. Preservation: Amber inclusion. Repository: JB. Locality: Dominican Republic, No stated.

Horizon: No stated, Lower Miocene.

**Family Siphonorhinidae Cook, 1895**

Siphonorhinidae indet: Wesener & Moritz, 2018: 1134.

Referred material: BuB1086: ♂; BuB997: ♂; BuB1123: ♀; BuB1150: ♀; BuB1822: ♀; BuB1838: ♀; BuB1842: ♀; BuB1845: ♀; BuB1851; BuB2979; BuB3243: ♀; BuB3283. Preservation: Amber inclusions. Repository: BuB. Locality: Myanmar, Kachin, Hukawng Valley, Noije Bum mine. Horizon: No stated, lowermost Upper Cretaceous, Upper Albian/Lower Cenomanian.

**Subterclass Eugnatha Attems, 1898 Superorder Juliformia Attems, 1926**

Juliformia indet: Riquelme & Hernández-Patricio, 2018: 639, Figs 3A-B.

Referred material: MALM.306; AM.CH.Id34; AM.CH.Id35. Preservation: Amber inclusions. Repository: MALM; AM.CH. Locality: Mexico, Chiapas, Simojovel, La Pimienta mine (MALM.306), Los Pocitos mine (AM.CH.Id34; AM.CH.Id35). Horizon: Uppermost Simojovel Formation, Lower Miocene.

**Order Incertae sedis**

**†Family Proglomeridae Fritsch, 1899**

**†Genus *Archiscudderia* Fritsch, 1899**

**(93) †Archiscudderia coronata Fritsch, 1899**

*Archiscudderia coronata* Fritsch, 1899: 36, pl 148, Figs 4-5, pl 149, Fig.3; Štamberg & Zajíc, 2008: 88, Fig.92.

Referred material: Holotype: Me 28. Preservation: Compression. Repository: CGH. Locality: Czech Republic, Pilsen, Nýřany. Horizon: Kladno Formation, Upper Carboniferous.

**(94) †Archiscudderia paupera Fritsch, 1899**

*Archiscudderia paupera* Fritsch, 1899: 35, pl 147, Fig.4, text-fig 341; Štamberg & Zajíc, 2008: 88, Fig.91.

Referred material: Me 26; M 870. Preservation: Compressions. Repository: CGH. Locality: Czech Republic, Pilsen, Nýřany. Horizon: Kladno Formation, Upper Carboniferous.

**(95) †Archiscudderia problematica Fritsch, 1899**

*Archiscudderia problematica* Fritsch, 1899: 37, pl 151, Figs 1-6; Štamberg & Zajíc, 2008: 89, Fig.94.

Referred material: Holotype: Me 33. Repository: Compression. Repository: CGH. Locality: Czech Republic, Pilsen, Nýřany. Horizon: Kladno Formation, Upper Carboniferous.

**(96) †Archiscudderia regularis Fritsch, 1899**

*Archiscudderia regularis* Fritsch, 1899: 37, pl 148, Figs 1-3; Štamberg & Zajíc, 2008: 88, Fig.93.

Referred material: Holotype: Me 27. Repository: Compression. Repository: CGH. Locality: Czech Republic, Pilsen, Nýřany. Horizon: Kladno Formation, Upper Carboniferous.

**(97) †Archiscudderia tapeta Fritsch, 1899**

*Archiscudderia tapeta* Fritsch, 1899: 36, pl 146, Figs 2-3; Štamberg & Zajíc, 2008: 88.

Referred material: Holotype: NHM2008z0097/001. Repository: Compression. Repository: NHM. Locality: Czech Republic, Pilsen, Nýřany. Horizon: Kladno Formation, Upper Carboniferous.

**†Superfamily Xyloiuloidea Attems, 1926 Family *Incertae sedis***

**†Genus *Karstiulus* Hannibal & May, 2020**

**(98) †*Karstiulus fortsillensis* Hannibal & May, 2020**

*Karstiulus fortsillensis* Hannibal & May, 2020: 589, Fig.1.

Referred material: Holotype: OU 12154. Preservation: Impression. Repository: OU. Locality: USA, Comanche County, Oklahoma, Richards Spur. Horizon: Fort Sill fissures, Cisuralian Series, Lower Permian.

**†Family Gaspestriidae Wilson, 2006**

**†Genus S*igmastria* Wilson, 2006**

**(99) †*Sigmastria dilata* Wilson, 2006**

*Sigmastria dilata* Wilson, 2006: 640, Fig.3.

Referred material: Holotype: NMS G.1957.1.5149. Preservation: Impression. Repository: NMS. Locality: UK, Scotland, Carmyllie. Horizon: Dundee Flagstone Formation, Lower Devonian.

**†Genus *Gaspestria* Wilson, 2006**

**(100) †Gaspestria genselorum Wilson, 2006**

*Gaspestria genselorum* Wilson, 2006: 640, Figs 1-2.

Referred material: Holotype: MHNM 01.120; other material examined: NBMG 10089; NBMG 10090; NBMG10091. Preservation: Impressions. Repository: MHNM, NBMG. Locality: Canada, Quebec, Gensel and Andrews, locality W. Horizon: Battery Point Formation, Lower Devonian, Lochkovian-Emsian.

**†Family Nyraniidae Hoffman, 1969**

**†Genus *Nyranius* Frič, 1875**

**(101) †Nyranius costulatus (Frič, 1875)**

*Julus costulatus* Frič, 1875: 74.

*Xylobius costulatus*: Fritsch, 1895: 2; Fritsch, 1899: 31, pl 144, Figs 4-5.

*Nyranius costulatus*: Hoffman, 1963: 172; Štamberg & Zajíc, 2008: 87, Fig.89.

Referred material: Holotype: Me 18. Preservation: Compression. Repository: CGH. Locality: Czech Republic, Pilsen, Nýřany. Horizon: Kladno Formation, Upper Carboniferous.

**(102) †*Nyranius tabulatus* (Fritsch, 1899)**

*Xylobius tabulatus* Fritsch, 1899: 32, pl 144, Figs 9-12.

*Nyranius tabulatus*: Hoffman, 1963: 172; Štamberg & Zajíc, 2008: 87, Fig.90.

Referred material: Holotype: M 1001. Preservation: Compression. Repository: CGH. Locality: Czech Republic, Pilsen, Nýřany. Horizon: Kladno Formation, Upper Carboniferous.

**†Family Xyloiulidae Cook, 1895**

**†Genus *Blanziulus* Langiaux & Sotty, 1976**

**(103) †*Blanziulus parriati* Langiaux & Sotty, 1976**

*Blanziulus parriati* Langiaux & Sotty, 1976: 43, pl 4, text-figs 1-2; Lheritier *et al*., 2023:11, Figs 2D-H, 4B, D, F, 5, 8I, K-M, 9D, 10C-D, 11D, 13C.

Referred material: Holotype: MNHN.F.SOT.76-H-1-A; MNHN.F.SOT.2114a; MNHN.F.SOT.5148. Preservation: Impressions. Repository: MNHN. Locality: France, Montceau-les-Mines. Horizon: Ferruginous mudstone strata, Upper Carboniferous.

**†Genus *Xyloiulus* Cook, 1895**

*Xyloiulus* sp.: Baird, 1958: 239-240.

Referred material: MCZ 5448; MCZ 5449; MCZ 5450. Preservation: Impressions. Repository: MCZ. Locality: Canada, Nova Scotia. Horizon: Cumberland Group, Joggins Formation, Upper Carboniferous.

Referred material: MCZ 5268; MCZ 5329; MCZ 5321. Preservation: Impressions. Repository: MCZ. Locality: USA, Ohio, Jefferson County, Linton. Horizon: Upper Freeport Coal Member, Upper Carboniferous.

Referred material: MCZ 5451. Preservation: Compression. Repository: MCZ. Locality: USA, Dallas, Texas, Coffee Creek. Horizon: Arroyo Formation, Lower Permian.

**(104) †*Xyloiulus bairdi* Hoffman, 1963**

*Xyloiulus bairdi* Hoffman, 1963:171, pl 24, Figs 2-3, text-figs 1-4.

Referred material: MCZ 5268. Preservation: Impression. Repository: MCZ. Locality: USA, Jefferson County, Ohio, Linton. Horizon: Upper Freeport Coal Member, Upper Carboniferous.

**(105) † *Xyloiulus fractus* Scudder, 1868**

*Xyloiulus fractus* Scudder, 1868 (in McClennen *et al*., 2017) Scudder, 1895: 61.

Referred material: No stated. Preservation: Compression. Repository: BMNH. Locality: Canada, Nova Scotia, Coal Mine Point, Divison 4. Horizon: Cumberland Group, Joggins Formation, Upper Carboniferous.

**(106) †Xyloiulus frustulentus (Scudder, 1890)**

*Xylobius frustulentus* Scudder, 1890: 438, pl 37, Figs 4-6.

*Xyloiulus frustulentus*: Hoffman, 1963: 172.

Referred material: Holotype: USNM PAL37995. Preservation: Impression. Repository: USNM. Locality: USA, Illinois, Grundy County, Morris, Mazon Creek. Horizon: Carbondale Formation, Upper Carboniferous.

**(107) †*Xyloiulus mazonus* (Scudder, 1890)**

*Xylobius mazonus* Scudder, 1890: 439, pl 37, Figs 7-11.

*Xyloiulus mazonus*: Hoffman, 1963: 172.

Referred material: Holotype: USNM PAL38036. Preservation: Impression. Repository: USNM. Locality: USA, Illinois, Grundy County, Morris, Mazon Creek. Horizon: Carbondale Formation, Upper Carboniferous.

**(108) †*Xyloiulus platti* (Baldwin, 1911)**

*Xylobius platti* Baldwin, 1911: 79, Fig.1.

Referred material: No stated. Preservation: Impression. Repository: BMNH. Locality: UK, England, Sparth Bottoms, Rochdale. Horizon: Shale strata, Upper Carboniferous.

**(109) †*Xyloiulus pstrossi* (Fritsch, 1899)**

*Xylobius pstrossi* Fritsch, 1899: 31, pl 144, Figs 6-8; Hoffman, 1963: 172; Štamberg & Zajíc, 2008: 86, Fig.88.

Referred material: Me 19; M 1039. Preservation: Compressions. Repository: CGH. Locality: Czech Republic, Pilsen, Nýřany. Horizon: Kladno Formation, Upper Carboniferous.

**(110) †*Xyloiulus renieri* Pruvost, 1930**

*Xyloiulus renieri* Pruvost, 1930: 169, pl 7, Figs 5, text-fig 4.

Referred material: No stated. Preservation: Impression. Repository: IRSN. Locality: Belgium, Courcelles. Horizon: Siliciclastic strata, Upper Carboniferous.

**(111) †*Xyloiulus sellatus* (Fritsch, 1899)**

*Xylobius sellatus* Fritsch, 1899: 32, pl 140, Fig.11.

*Xyloiulus sellatus*: Hoffman, 1963: 172; Štamberg & Zajíc, 2008: 87.

Referred material: Holotype: No. 141. Preservation: Compression. Repository: CGH. Locality: Czech Republic, Pilsen, Nýřany. Horizon: Kladno Formation, Upper Carboniferous.

**(112) †Xyloiulus sigillariae (Dawson, 1860)**

*Xylobius sigillariae* Dawson, 1860: 271; Scudder, 1868: pl 16, Fig.4; Scudder, 1895: 61; Copeland, 1957: 53, pl 11, Fig.6.

*Xyloiulus sigillariae*: Hoffman, 1963: 169.

Referred material: Holotype. Preservation: Impression. Repository. BMNH. Locality: UK, England, Sparth Bottoms, Rochdale. Horizon: Shale strata, Upper Carboniferous.

**(113) †*Xyloiulus similis* (Scudder, 1868)**

*Xylobius similis* Scudder, 1868: Scudder, 1895: 61, pl 5, Figs 1-2.

Referred material: No stated. Preservation: Compression. Repository: BMNH. Locality: Canada, Nova Scotia, Coal Mine Point, Divison 4. Horizon: Cumberland Group, Joggins Formation, Upper Carboniferous.

**Order Julida Brandt, 1833**

Julida indet: Duncan *et al*., 1988: 848. Álvarez-Rodríguez *et al*., 2023, Fig.3B (this work).

Referred material: QM F34596. Preservation: Compression. Repository: QM. Locality: Australia, Queensland. Horizon: System B Formation, Lower Miocene.

Referred material: CPAL.124: ♀. Preservation: Amber inclusion. Repository: CPAL-UAEM. Locality: Mexico, Chiapas, Simojovel, Guadalupe Victoria mine. Horizon: Uppermost Simojovel Formation, Lower Miocene.

**Superfamily Juloidea Leach, 1814 Family Julidae Leach, 1814**

**†Genus *Bertkaupolypus* Verhoeff, 1926**

**(114) †Bertkaupolypus antiquus (Bertkau, 1878)**

*Julus antiquus* Bertkau, 1878: 360, pl 5, Fig.8.

*Pseudoiulus antiquus*: Verhoeff, 1897: 280.

*Bertkaupolypus antiquus*: Hoffman, 1969: 591.

Referred material: No stated. Preservation: Compression. Repository: BMNH. Locality: Germany, Nordrhein-Westfalen. Horizon: Lignite strata, Upper Oligocene.

**Genus *Julus* Linnaeus, 1758**

*Julus* sp.: Janossy, 1986: 57.

Referred material: No stated. Preservation: Compression. Repository: HNHM. Locality: Hungary, Barany, Villany locality 6. Horizonte: No stated, Lower Pleistocene.

**(115) †*Julus badius* Menge, 1854**

*Julus badius* Menge, 1854: 13.

Referred material: No stated. Preservation: Amber inclusion. Repository: WPM. Locality: Baltic. Horizon: Prussian Formation, Upper Eocene.

**(116) †*Julus cavicola* Grinnell, 1908**

*Julus cavicola* Grinnell, 1908: 210, pl 15, Figs 1, 5, 10, 12.

Referred material: Holotype: UCMP 10007. Preservation: Impression. Repository: UCMP. Locality: USA, Los Angeles County, California, Rancho La Brea. Horizon: Poorly lithified tar, Upper Pleistocene.

**(117) †*Julus florissantellus* Cockerell, 1907**

*Julus florissantellus* Cockerell, 1907: 605, Fig.1.

Referred material: Holotype. Preservation: Compression. Repository: AMNH. Locality: USA, Teller County, Colorado.

Horizon: Florissant Formation, Upper Eocene.

**(118) †*Julus laevigatus* Koch & Berendt, 1854**

*Julus laevigatus* Koch & Berendt, 1854: 12, pl 1, Fig.4.

Referred material: No stated. Preservation: Amber inclusion. Repository: MfN. Locality: Baltic. Horizon: Prussian Formation, Upper Eocene.

**(119) †*Julus occidentalis* Grinnell, 1908**

*Julus occidentalis* Grinnell, 1908: 209, pl 15, Figs 9, 11.

Referred material: UCMP 10005; UCMP 10006. Preservation: Impressions. Repository: UCMP. Locality: United States, Shasta County, California, Samwel Cave No. I. Horizon: Claystone strata, Upper Pleistocene.

**(120) †*Julus peii* Chia & Liu, 1950**

*Julus peii* Chia & Liu, 1950: 25, pl 1, Figs 9-10.

Referred material: No stated. Preservation: Impression (fragmentary). Repository: No stated. Locality: China, Choukoutien, Loc. l, 15, and 4; Upper Cave. Horizon: Lower-Upper Pleistocene.

**(121) †*Julus politus* Menge, 1854**

*Julus politus* Menge, 1854: 13.

Referred material: No stated. Preservation: Amber inclusion. Repository: WPM. Locality: Baltic. Horizon: Prussian Formation, Upper Eocene.

**(122) †*Julus rubens* Menge, 1854**

*Julus rubens* Menge, 1854: 13.

Referred material: No stated. Preservation: Amber inclusion. Repository: WPM. Locality: Baltic. Horizon: Prussian Formation, Upper Eocene

**(123) †*Julus suevicus* Dietlen, 1902**

*Julus suevicus* Dietlen, 1902: 84.

Referred material: No stated. Preservation: Impression. Repository: SMNS. Locality: Germany, Wurttemberg, Böttingen. Horizon: Thermalsinterkalk Formation, Upper Miocene.

**(124) †*Julus terrestris* Linnaeus, 1758**

*Julus* cf. *terrestris* Chia & Liu, 1950: 24, pl I, Figs 1-8.

Referred material: No stated. Preservation: Impression (fragmentary). Repository: No stated. Locality: China, Choukoutien, Loc. l, 15, 3; and Upper Cave. Horizon: Lower-Upper Pleistocene.

**Superfamily Parajuloidea Bollman, 1893**

**Family Parajulidae Bollman, 1893**

**Genus *Parajulus* Humbert & Saussure, 1869**

**(125) †Parajulus cockerelli Miner, 1926**

*Parajulus cockerelli* Miner, 1926: 3, Figs 1-5.

Referred material: Holotype: AMNH 22564. Preservation: Compression. Repository: AMNH. Locality: USA, Teller County, Colorado. Horizon: Florissant Formation, Upper Eocene.

**(126) †*Parajulus onychis* Pierce, 1951**

*Parajulus onychis* Pierce, 1951: 41, pl 16, text-figs 8-9.

Referred material: Holotype: LACM BQ19. Preservation: Impression. Repository: LACM. Locality: USA, Arizona, Bonner quarry. Horizon: Onyx Marble Formation, Lower Pliocene.

**Order Spirobolida Cook, 1895**

Spirobolida indet: Riquelme & Hernández-Patricio, 2018: 638, Figs 1D-E, Figs 2A-C; Wesener & Moritz, 2018: 1136, Fig.2E.

Referred material: MALM.313; MALM.18: juvenile. Preservation: Amber inclusions. Repository: MALM. Locality: Mexico, Chiapas, Simojovel, La Pimienta mine. Horizon: Uppermost Simojovel Formation, Lower Miocene.

Referred material: BuB830: ♀; BuB916: ♂ (head missing); BuB1795: ♂; BuB1840: ♀; BuB2616; BuB3020; BuB3000: ♂; BuB3260: ♀; ZFMK MYR7373: ♂. Preservation: Amber inclusions. Repository: BuB; ZFMK-MYR. Locality: Myanmar, Kachin, Hukawng Valley, Noije Bum mine. Horizon: No stated, lowermost Upper Cretaceous, Upper Albian/Lower Cenomanian.

**Suborder Spirobolidea Cook, 1895**

**Family Atopetholidae Chamberlin, 1918**

**†Genus *Gobiulus* Dzik, 1975**

**(127) †Gobiulus sabulosus Dzik, 1975**

*Gobiulus sabulosus* Dzik, 1975: 18, Figs 1-5.

Referred material: Holotype: ZPAL Mg Myr/6. Preservation: Impression. Repository: ZPAL. Locality: Mongolia, Omnogov. Horizon: Baruungoyot Formation, Upper Cretaceous, Middle Campanian.

**Family Spirobolidae Bollman, 1893**

**Genus *Hiltonius* Chamberlin, 1918**

**(128) †*Hiltonius australis* (Grinnell, 1908)**

*Spirobolus australis* Grinnell, 1908: 210, pl 15, Figs 13-14.

*Hiltonius australis*: Hoffman, 1969: 587.

Referred material: Holotype: UCMP 10008/9. Preservation: Impression. Repository: UCMP. Locality: USA, Los Angeles County, California, Rancho La Brea. Horizon: Poorly lithified tar, Upper Pleistocene.

**Family Rhinocricidae Brölemann, 1913**

**Genus *Rhinocricus* Karsch, 1881**

*Rhinocricus* sp.: Baalbergen & Donovan, 2013: 6.

Referred material: BMNH IU 1-IU 13; BMNH IU 17-IU 19; RGM 789 601; RGM 789 602: 4 specimens; RGM 789603. Preservation: Impressions. Repository: BMNH; RGM. Locality: Jamaica, Red Hills Road Cave. Horizon: Limestone strata, Upper Pleistocene.

**Order Spirostreptida Brandt, 1833**

**Suborder Cambalidea Cook, 1895**

Cambalidea indet: Zhang, 2017: 172. Wesener & Moritz, 2018: 1136, Fig.2D.

Referred material: No stated. BuB1115; BuB1144; BuB1165: ♀; BuB1824; BuB1825: ♀; BuB1826: ♀; BuB1990: ♀; BuB1955: ♀; BuB1962: ♂; BuB2981; BuB3005; BuB3012; BuB3250; BuB3255: ♂; ZFMK MYR06121: ♂; ZFMK MYR07368: ♂; ZFMK MYR07369: ♂; ZFMK MYR07370:2 ♀. Preservation: Amber inclusions. Repository: NMS: BuB; ZFMK-MYR. Locality: Myanmar, Kachin, Hukawng Valley, Noije Bum mine. Horizon: No stated, lowermost Upper Cretaceous, Upper Albian/Lower Cenomanian.

**Family Cambalidae Cook, 1895**

Cambalidae indet: Wesener & Moritz, 2018: 1136.

Referred material: ZFMK MYR06696: ♂. Preservation: Amber inclusion. Repository: ZFMK. Locality: Myanmar, Kachin, Hukawng Valley, Noije Bum mine. Horizon: No stated, lowermost Upper Cretaceous, Upper Albian/Lower Cenomanian.

**†Genus *Protosilvestria* Handschin, 1944**

**(129) †Protosilvestria sculpta Handschin, 1944**

*Protosilvestria sculpta* Handschin, 1944: 4, pl 1, Figs 1-6, text-fig 1; Mauries, 1992: 24, Figs 1, 6-8, 11-15.

Referred material: No stated. Preservation: Compression. Repository: NHMB. Locality: France, Caylux. Horizon: No stated, Upper Oligocene.

**†Family Electrocambalidae Moritz & Wesener, 2021**

Electrocambalidae indet: Moritz & Wesener, 2021: 39.

Referred material: BuB1144. Preservation: Amber inclusion. Repository: BuB. Locality: Myanmar, Kachin, Hukawng Valley, Noije Bum mine. Horizon: No stated, lowermost Upper Cretaceous, Upper Albian/Lower Cenomanian.

**†Genus *Electrocambala* Moritz & Wesener, 2021**

*Electrocambala* sp.: Moritz & Wesener, 2021: 32.

Referred material: BuB1990: ♀. Preservation: Amber inclusion. Repository: BuB. Locality: Myanmar, Kachin, Hukawng Valley, Noije Bum mine. Horizon: No stated, lowermost Upper Cretaceous, Upper Albian/Lower Cenomanian.

**(130) †*Electrocambala cretacea* Moritz & Wesener, 2021**

*Electrocambala cretacea* Moritz & Wesener, 2021: 30, Fig.3.

Referred material: Holotype: ZFMK-MYR7370: ♀. Preservation: Amber inclusion. Repository: ZFMK-MYR. Locality: Myanmar, Kachin, Hukawng Valley, Noije Bum mine. Horizon: No stated, lowermost Upper Cretaceous, Upper Albian/Lower Cenomanian.

**(131) †*Electrocambala ornata* Moritz & Wesener, 2021**

*Electrocambala ornata* Moritz & Wesener, 2021: 26, Figs 1-2.

Referred material: Holotype: ♂; Paratype: ZFMK-MYR7370: ♀; additional material: BuB1825: ♂; BuB1962: ♂. Preservation: Amber inclusions. Repository: ZFMK-MYR; BuB. Locality: Myanmar, Kachin, Hukawng Valley, Noije Bum mine. Horizon: No stated, lowermost Upper Cretaceous, Upper Albian/Lower Cenomanian.

**†Genus *Kachincambala* Moritz & Wesener, 2021**

*Kachincambala* sp.: Moritz & Wesener, 2021: 38.

Referred material: BuB4098: ♀, BuB1115; ZFMK-MYR6121. Preservation: Amber inclusions. Repository: BuB; ZFMK-MYR. Locality: Myanmar, Kachin, Hukawng Valley, Noije Bum mine. Horizon: No stated, lowermost Upper Cretaceous, Upper Albian/Lower Cenomanian.

**(132) †*Kachincambala distorta* Moritz & Wesener, 2021**

*Kachincambala distorta* Moritz & Wesener, 2021: 35, Fig.5.

Referred material: Holotype: ZFMK-MYR7368: ♂. Preservation: Amber inclusion. Repository: ZFMK-MYR. Locality: Myanmar, Kachin, Hukawng Valley, Noije Bum mine. Horizon: No stated, lowermost Upper Cretaceous, Upper Albian/Lower Cenomanian.

**(133) †*Kachincambala muelleri* Moritz & Wesener, 2021**

*Kachincambala muelleri* Moritz & Wesener, 2021: 33, Fig.4.

Referred material: Holotype: ZFMK-MYR10225: ♂. Preservation: Amber inclusion. Repository: ZFMK-MYR. Locality: Myanmar, Kachin, Hukawng Valley, Noije Bum mine. Horizon: No stated, lowermost Upper Cretaceous, Upper Albian/Lower Cenomanian.

**Family Pseudonannolenidae Silvestri, 1895**

**Genus *Epinannolene* Brölemann, 1903**

*Epinannolene* sp.: Santiago-Blay & Poinar, 1992: 366, Fig.6.

Referred material: GOP DC-3-12: juvenile. Preservation: Amber inclusion. Repository: GOP. Locality: Dominican Republic, Santiago, Cordillera Septentrional. Horizon: El Mamey Formation, Lower Miocene.

**Superorder Nematophora Verhoeff, 1913**

**Order Callipodida Pocock, 1894**

Callipodida indet: Wesener & Moritz, 2018: 1136, Fig.2C.

Referred material: ZFMK MYR07366: ♀. Preservation: Amber inclusion. Repository: ZFMK MYR. Locality: Myanmar, Kachin, Hukawng Valley, Noije Bum mine. Horizon: No stated, lowermost Upper Cretaceous, Upper Albian/Lower Cenomanian.

**Family Incertae sedis**

**†Genus Hannibaliulus Shear et al., 2009**

**(134) †Hannibaliulus wilsonae Shear et al., 2009**

*Hannibaliulus wilsonae* Shear *et al*., 2009: 6, Figs 2-8.

Referred material: Holotype: MYR1 a, b; Paratype: MYR10 a, b; other material: MYR2a; MYR3a, b; MYR4; MYR5; MYR6; MYR7a, b; MYR8a, b; MYR9a, b. Preservation: Compressions. Repository: IUT. Locality: France, Moselle, Bust-Hangviller, locality 238. Horizon: Grès à Voltzia Formation, Middle Triassic.

**†Suborder Burmanopetalidea Stoev, Moritz & Wesener, 2019**

**†Family Burmanopetalidae Stoev, Moritz & Wesener, 2019**

**†Genus *Burmanopetalum* Stoev, Moritz & Wesener, 2019**

**(135) †*Burmanopetalum inexpectatum* Stoev, Moritz & Wesener, 2019**

*Burmanopetalum inexpectatum* Stoev *et al*., 2019: 83, Figs 1-3.

Referred material: Holotype: ZFMK-MYR07366: ♀. Preservation: Amber inclusion. Repository: ZFMK-MYR. Locality: Myanmar, Kachin, Hukawng Valley, Noije Bum mine. Horizon: No stated, lowermost Upper Cretaceous, Upper Albian/Lower Cenomanian.

**Order Chordeumatida Pocock, 1894**

Chordeumatida indet: Wesener & Moritz, 2018: 1135.

Referred material: BuB0974; BuB1978; BuB2978: ♂. Preservation: Amber inclusions. Repository: BuB. Locality: Myanmar, Kachin, Hukawng Valley, Noije Bum mine. Horizon: No stated, lowermost Upper Cretaceous, Upper Albian/Lower Cenomanian.

**Suborder Craspedosomatidea Cook, 1895 Superfamily Craspedosomatoidea Gray in Jones, 1843 Family Craspedosomatidae Gray in Jones, 1843**

**Genus *Craspedosoma* Leach, 1814**

**(136) †Craspedosoma aculeatum Menge, 1854**

*Craspedosoma aculeatum* Menge, 1854: 14.

Referred material: No stated. Preservation: Amber inclusion. Repository: WPM. Locality: Baltic. Horizon: Prussian Formation, Upper Eocene.

**(137) †*Craspedosoma affine* Koch & Berendt, 1854**

*Craspedosoma affine* Koch & Berendt, 1854: 13, pl 1, Fig.5a.

Referred material: No stated. Preservation: Amber inclusion. Repository: MfN. Locality: Baltic. Horizon: Prussian Formation, Upper Eocene.

**(138) †*Craspedosoma angulatum* Koch & Berendt, 1854**

*Craspedosoma angulatum* Koch & Berendt, 1854: 13, pl 1, Fig.5.

Referred material: No stated. Preservation: Amber inclusion. Repository: MfN. Locality: Baltic. Horizon: Prussian Formation, Upper Eocene.

**(139) †Craspedosoma armatum Menge, 1854**

*Craspedosoma armatum* Menge, 1854: 14.

Referred material: No stated. Preservation: Amber inclusion. Repository: WPM. Locality: Baltic. Horizon: Prussian Formation, Upper Eocene.

**(140) †Craspedosoma cylindricum Menge, 1854**

*Craspedosoma cylindricum* Menge, 1854: 14.

Referred material: No stated. Preservation: Amber inclusion. Repository: WPM. Locality: Baltic. Horizon: Prussian Formation, Upper Eocene.

**(141) †Craspedosoma obtusangulum Menge, 1854**

*Craspedosoma obtusangulum* Menge, 1854: 14.

Referred material: No stated. Preservation: Amber inclusion. Repository: WPM. Locality: Baltic. Horizon: Prussian Formation, Upper Eocene.

**(142) †Craspedosoma setosum Menge, 1854**

*Craspedosoma setosum* Menge, 1854: 14.

Referred material: No stated. Preservation: Amber inclusion. Repository: WPM. Locality: Baltic. Horizon: Prussian Formation, Upper Eocene.

**†Genus *Euzonus* Menge, 1854**

**(143) †*Euzonus collulum* Menge, 1854**

*Euzonus collulum* Menge, 1854: 14.

Referred material: No stated. Preservation: Amber inclusion. Repository: WPM. Locality: Baltic. Horizon: Prussian Formation, Upper Eocene.

**Suborder Heterochordeumatidea Shear, 2000**

**Superfamily Heterochordeumatoidea Pocock, 1894**

**Family Heterochordeumatidae Pocock, 1894**

Heterochordeumatidae indet: Wesener & Moritz, 2018: 1135.

Referred material: Wu F2806/Bu/CJW: ♀; BuB0642: ♀; BuB0833: two ♂; BuB0899: ♀; BuB1141: ♂; BuB1410: ♀; BuB1411: ♀; BuB1412: ♀; BuB1823: ♂; BuB1827; BuB2685: ♂; BuB3022: ♀; BuB3030; BuB3051: ♀; BuB3056; ZFMK MYR05545: ♂; ZFMK MYR06123: ♂; ZFMK MYR06624: ♂; ZFMK MYR07367: ♂. Preservation: Amber inclusions. Repository: Wu; BuB; ZFMK-MYR. Locality: Myanmar, Kachin, Hukawng Valley, Noije Bum mine. Horizon: No stated, lowermost Upper Cretaceous, Upper Albian/Lower Cenomanian.

**Order Stemmiulida Pocock, 1895**

Stemmiulida indet: Wesener & Moritz, 2018: 1135, Fig.2B.

Referred material: BuB994: ♂; BuB1961: ♀; BuB1968; BuB2998; BuB3009; BuB3038: ♀; BuB3241; ZFMK MYR07378: ♂. Preservation: Amber inclusions. Repository: BuB; ZFMK-MYR. Locality: Myanmar, Kachin, Hukawng Valley, Noije Bum mine. Horizon: No stated, lowermost Upper Cretaceous, Upper Albian/Lower Cenomanian.

**Family Stemmiulidae Pocock, 1894**

Stemmiulidae indet: Santiago-Blay & Poinar, 1992: 366, Figs 4-5; Riquelme & Hernández-Patricio, 2018: 643, Figs 3A-B; Álvarez-Rodríguez *et al*., 2023 (this work).

Referred material: GOP DC-3-13: ♀. Preservation: Amber inclusion. Repository: GOP. Locality: Dominican Republic, Santiago, Cordillera Septentrional. Horizon: El Mamey Formation, Lower Miocene.

Referred material: CPAL.104; CPAL.105: juvenile; MALM.304; MALM.308; MALM.309. Preservation: Amber inclusions. Repository: CPAL; MALM. Locality: Mexico, Chiapas, Simojovel, Guadalupe Victoria mine (CPAL.104, CPAL.105), La Pimienta mine (MALM.304, MALM.308, MALM.309). Horizon: Uppermost Simojovel Formation, Lower Miocene.

New records: CPAL.141: ♀. Preservation: Amber inclusion. Repository: CPAL. Locality: Mexico, Chiapas, Simojovel, Los Pocitos mine. Horizon: Uppermost Simojovel Formation, Lower Miocene.

**†Genus *Parastemmiulus* Riquelme *et al*., 2013**

**(144) †Parastemmiulus elektron Riquelme et al., 2013**

*Parastemmiulus elektron* Riquelme *et al*., 2013: 6, Figs 3-9.

Referred material: Holotype: SUCCINUM14.INAH.2661: ♀. Preservation: Amber inclusion. Repository: SUCCINUM/INAH. Locality: Mexico, Chiapas, Simojovel, Guadalupe Victoria mine. Horizon: Uppermost Simojovel Formation, Lower Miocene.

**Order Siphoniulida Cook, 1895 Family Siphoniulidae Pocock, 1894**

**Genus *Siphoniulus* Pocock, 1894**

**(145) †Siphoniulus muelleri Liu et al., 2017**

*Siphoniulus muelleri* Liu *et al*., 2017: 102, Figs 1-2.

Referred material: Holotype: ZFMK MYR 6098: ♀. Preservation: Amber inclusion. Repository: ZFMK-MYR. Locality: Myanmar, Kachin, Hukawng Valley, Noije Bum mine. Horizon: No stated, lowermost Upper Cretaceous, Upper Albian/Lower Cenomanian.

**(146) †Siphoniulus preciosus Liu et al., 2017**

*Siphoniulus preciosus* Liu *et al*., 2017: 104, Figs 3-5.

Referred material: Holotype: ZFMK MYR 5543: ♀. Preservation: Amber inclusion. Repository: ZFMK-MYR. Locality: Myanmar, Kachin, Hukawng Valley, Noije Bum mine. Horizon: No stated, lowermost Upper Cretaceous, Upper Albian/Lower Cenomanian.

**Superorder Merochaeta Cook, 1895 Order Polydesmida Pocock, 1887**

Polydesmida indet: Riquelme & Hernández-Patricio, 2018: 640, Figs 3A-B, F; Wesener & Moritz, 2018: 1136, Fig.2; Álvarez-Rodríguez *et al*., 2023, Figs 4D-E (this work).

Referred material: CPAL.103: juvenile; CPAL.106: ♀; CPAL.107: juvenile; CPAL.108: juvenile; CPAL.113: juvenile; MACH.21; MALM.301: juvenile; MALM.302: juvenile; MALM.310; MALM.311: juvenile; MALM.312: juvenile. Preservation: Amber inclusions. Repository: CPAL-UAEM; MACH; MALM. Locality: Mexico, Chiapas, Simojovel, Guadalupe Victoria mine (CPAL.106, CPAL.107), La Pimienta mine (CPAL.103, CPAL.113, MALM.301, MALM.302, MALM.310, MALM.311, MALM.312), Los Pocitos mine (MACH.21); Totolapa, Rìo Salado mine (CPAL.108). Horizon: Uppermost Simojovel Formation, Lower Miocene.

Referred material: BuB600; BuB672; BuB818; BuB902: 6 specimens; BuB909; BuB911: ♂; BuB912: 4 specimens; BuB915; BuB966; BuB975: ♂; BuB976: ♂; BuB980: ♂; BuB983: ♀; BuB993: ♂; BuB995; BuB1029: ♂; BuB1031-A: ♂; BuB1035: ♂; BuB1084: ♀; BuB1085: ♀; BuB1146: ♂; BuB1148: ♀; BuB1149: ♂; BuB1154: 2 ♂, 5 ♀; BuB1155; ♀; BuB1156: ♀; BuB1414; BuB1548: ♀; BuB1794; ♀; BuB1830: 2 ♀; BuB1832: ♀; BuB1836: ♀; BuB1844: ♂, BuB1847: ♂; BuB1848: ♂; BuB1849: ♂; BuB1850; BuB1852: ♂; BuB1954A; BuB1957: 3 ♀; BuB1958; BuB1964: 2 ♀; BuB1967; BuB1975: ♂; BuB1985: ♀; BuB1986; BuB1987: ♀; BuB1989B; BuB1992; BuB1993; BuB2436; Bub2437; BuB2613; BuB2622; BuB2624; BuB2631; BuB2632: 2 ♀; BuB2639; BuB2640; BuB2645; BuB2646; BuB2647: ♀; BuB2648: ♀; BuB2653; BuB2672; BuB2683: ♀; BuB2684; BuB2686; BuB2687: ♀; BuB2688: ♀; BuB2960; BuB2967; BuB2968; BuB2969; BuB2970; BuB2972: 2 specimens; BuB2976: ♂; BuB2982: ♀; BuB2983: 2 ♀; BuB2987; BuB2988: ♀; BuB2992: ♂; BuB2994; BuB2999; BuB3001: ♀; BuB3002: ♀; BuB3003: ♂; BuB3004: ♀; BuB3008: ♂; BuB3011; BuB3021; BuB3023; BuB3025: ♀; BuB3029; BuB3032; BuB3034: ♂; BuB3039: ♂; BuB3040: ♀; BuB3028; BuB3049: ♂; BuB3055: ♂; BuB3238: ♂; BuB3246: ♀; BuB3251: ♀; BuB3252: ♀; BuB3253: ♀; BuB3254: ♀; BuB3256: ♂; BuB3265; BuB3267: 2 ♀; BuB3269; BuB3270; BuB3274; BuB3275: 2 specimens; BuB3276; BuB3277: ♀; BuB3278; BuB3279; BuB3285; BuB3286: ♀; BuB3293: ♂; ZFMK MYR06118: ♂; ZFMK MYR06120: ♀; ZFMK MYR07374: ♀; ZFMK MYR07377: ♀, 1 specimen; ZFMK MYR07375; ZFMK MYR07379: ♂; Wu F2817/Bu/CJW: ♀; Wu F3385/Bu/ CJW; Wu F3396/Bu/CJW; Wu F3397/Bu/CJW; RO my249: ♀; RO my301: ♀; RO my304: ♀. Preservation: Amber inclusions. Repository: BuB; ZFMK-MYR; Wu; RO. Locality: Myanmar, Kachin, Hukawng Valley, Noije Bum mine. Horizon: No stated, lowermost Upper Cretaceous, Upper Albian/Lower Cenomanian.

New records: CPAL.125.1; CPAL.125.2; CPAL.135; CPAL.137; CPAL.139; CPAL.140: juvenile; CPAL.144: ♀; CPAL.146; CPAL.149; CPAL.153; CPAL.156; CPAL.172: juvenile; CPAL.177; CPAL.178; CPAL.181: ♂; CPAL.183: ♀; CPAL.199; CPAL.215; CPAL.219; MACH.22: juvenile. Preservation: Amber inclusions. Repository: CPAL-UAEM; MACH. Locality: Mexico, Chiapas, Simojovel, Campo La Granja mine (CPAL.140), El Pabuchil mine (CPAL.178), Guadalupe Victoria mine (CPAL.137, CPAL.139), Los Pocitos mine (CPAL.125.1, CPAL.125.2, CPAL.135, CPAL.144, CPAL.153, CPAL.177, CPAL.181, CPAL.183), Monte Cristo mine (CPAL.146, CPAL.149, CPAL.156, CPAL.172, CPAL.199, CPAL.215, CPAL.219, MACH.22). Horizon: Uppermost Simojovel Formation, Lower Miocene.

Suborder Leptodesmidea Brölemann, 1916

**Superfamily Chelodesmoidea Cook, 1895**

**Family Chelodesmidae Cook, 1895**

Chelodesmidae indet: Santiago-Blay & Poinar, 1992: 366, Fig.8; Riquelme & Hernández-Patricio, 2018: 644, Fig.3E; Álvarez-Rodríguez *et al*., 2023 (this work).

Referred material: GOP DC-3-11: ♀. Preservation: Amber inclusion. Repository: GOP. Locality: Dominican Republic, Santiago, Cordillera Septentrional. Horizon: El Mamey Formation, Lower Miocene.

Referred material: AM.CH.ID33. Preservation: Amber inclusion. Repository: AM.CH. Locality: Mexico, Chiapas, Simojovel, Los Pocitos mine. Horizon: Uppermost Simojovel Formation, Lower Miocene.

New records: CPAL.211: ♀. Preservation: Amber inclusion. Repository: CPAL-UAEM. Locality: Mexico, Chiapas, Simojovel, Monte Cristo mine. Horizon: Uppermost Simojovel Formation, Lower Miocene.

**†Genus *Maatidesmus* Riquelme & Hernández-Patricio, 2014**

**(147) †*Maatidesmus paachtun* Riquelme & Hernández-Patricio, 2014**

*Maatidesmus paachtun* Riquelme *et al*., 2014: 5, Figs 2-3, 5A.

Referred material: Holotype: MALM.28: ♀. Preservation: Amber inclusion. Repository: MALM. Locality: Mexico, Chiapas, Simojovel, Guadalipe Victoria mine. Horizon: Uppermost Simojovel Formation, Lower Miocene.

**Genus *Caraibodesmus* Chamberlin, 1918**

**(148) †Caraibodesmus verrucosus Pocock, 1894**

*Caraibodesmus verrucosus* Donovan & Veltkamp, 1994: 359, Fig.4.

Referred material: Holotype: BMNH IU 15. Preservation: Impression. Repository: BMNH. Locality: Jamaica, Red Hills Road Cave. Horizon: Limestone strata, Upper Pleistocene.

**Genus *Chondrotropis* Loomis, 1936**

*Chondrotropis* sp.: Donovan & Veltkamp, 1994: 360.

Referred material: Holotype: BMNH IU 16. Preservation: Impression. Repository: BMNH. Locality: Jamaica, Red Hills Road Cave. Horizon: Limestone strata, Upper Pleistocene.

**Superfamily Platyrhacoidea Pocock, 1895**

**Family Platyrhacidae Pocock, 1895**

Platyrhacidae indet: Riquelme & Hernández-Patricio, 2018: 642.

Referred material: CPAL.101: ♀; CPAL.110: juvenile. Preservation: Amber inclusions. Repository: CPAL-UAEM. Locality: Mexico, Chiapas, Simojovel, Guadalupe Victoria mine. Horizon: Uppermost Simojovel Formation, Lower Miocene.

**†Genus *Anbarrhacus* Riquelme & Hernández-Patricio, 2014**

**(149) †*Anbarrhacus adamantis* Riquelme & Hernández-Patricio, 2014**

*Anbarrhacus adamantis* Riquelme *et al*., 2014: 8, Fig.4.

Referred material: Holotype: IGM 4544: juvenile. Preservation: Amber inclusion. Repository: IGL-UNAM. Locality: Mexico, Chiapas, Simojovel, Guadalupe Victoria mine. Horizon: Uppermost Simojovel Formation, Lower Miocene.

**Genus *Nyssodesmus* Cook, 1896**

*Nyssodesmus* sp.: Laurito & Valerio, 2018: 182, Figs 4C-D.

Referred material: Holotype: CFM-5199. Preservation: Compression. Repository: MNCR. Locality: Costa Rica, Alajuela, San Carlos, La Palmera. Horizon: Travertine outcrop, Upper Pleistocene.

**Superfamily Sphaeriodesmoidea Humbert & DeSaussure, 1869**

**Family Sphaeriodesmidae Humbert & DeSaussure, 1869**

Sphaeriodesmidae indet: Álvarez-Rodríguez *et al*., 2023, Fig.4B (this work).

New records: CPAL.130; CPAL.131; CPAL.152; CPAL.184: juvenile; CPAL.206: juvenile. Preservation: Amber inclusions. Repository: CPAL-UAEM. Locality: Mexico, Chiapas, Simojovel, Campo La Granja mine (CPAL.152), Guadalupe Victoria mine (CPAL.130, CPAL.131), Monte Cristo mine (CPAL.184), Palenque, Estrella de Belén mine (CPAL.206). Horizon: Uppermost Simojovel Formation, Lower Miocene.

**Genus *Cyclodesmus* Humbert & De Saussure, 1869**

**(150) Cyclodesmus porcellanus Pocock, 1894**

*Cyclodesmus porcellanus* Baalbergen & Donovan, 2012: 7, Fig.4.

Referred material: RGM 789 610; RGM 789 611; RGM 789 612. Preservation: Impressions. Repository: RGM. Locality: Jamaica, Red Hills Road Cave. Horizon: Limestone strata, Upper Pleistocene.

**Superfamily Xystodesmoidea Cook, 1895**

**Family Xystodesmidae Cook, 1895**

Xystodesmidae indet: Riquelme & Hernández-Patricio, 2018: 642, Figs 2D-F.

Referred material: CPAL.109: ♀; MALM.307: juvenile. Preservation: Amber inclusions. Repository: CPAL-UAEM; MALM. Locality: Mexico, Chiapas, Simojovel, Guadalupe Victoria mine (CPAL.109), La Pimienta mine (MALM.307). Horizon: Uppermost Simojovel Formation, Lower Miocene.

**Suborder Strongylosomatidea Brölemann, 1916**

**Family Paradoxosomatidae Daday, 1889**

**Genus *Orthomorpha* Bollman, 1893**

**(151) *Orthomorpha coarctata* (De Saussure, 1860)**

*Polydesmus coarctata* De Saussure, 1860: 297.

*Orthomorpha coarctata*: Li *et al*., 2021: 304, Figs 1b-g.

Referred material: LIV1: No stated; LIV2: No stated. Preservation: Impression (fragmentary). Repository: No stated.

Locality: China, Shaanxi, Luonan. Horizon: Middle Pleistocene.

***Suborder Polydesmidea Pocock, 1887***

***Infraorder Incertae sedis***

***Superfamily Incertae sedis***

***Family Incertae sedis***

**Genus *Dasyodontus* Loomis, 1936**

*Dasyodontus* sp.: Santiago-Blay & Poinar, 1992: 367, Fig.16.

Referred material: GOP DC-3-10: ♀. Preservation: Amber inclusion. Repository: GOP. Locality: Dominican Republic, Santiago, Cordillera Septentrional. Horizon: El Mamey Formation, Lower Miocene.

**Infraorder Oniscodesmoides Simonsen, 1990**

**Superfamily Pyrgodesmoidea Silvestri, 1896**

**Family Pyrgodesmidae Silvestri, 1896**

Pyrgodesmidae indet: Álvarez-Rodríguez *et al*., 2023, Fig.4F (this work).

New records: CPAL.120; CPAL.128; CPAL.136; CPAL.138; CPAL.155: ♀; CPAL.158; CPAL.167: ♀; CPAL.168; CPAL.173: ♀; CPAL.176: ♀; CPAL.185: ♀; CPAL.186: juvenile; CPAL.187: ♀; CPAL.200; CPAL.205: ♀; CPAL.210: ♀; CPAL.214: ♀; CPAL.217: ♀. Preservation: Amber inclusions. Repository: CPAL-UAEM. Locality: Mexico, Chiapas, Simojovel, El Chapayal mine (CPAL.167, CPAL.168, CPAL.187), El Porvenir mine (CPAL.128), Guadalupe Victoria mine (CPAL.120, CPAL.13), Los Pocitos mine (CPAL.176, CPAL.185, CPAL.186, CPAL.200, CPAL.205), Monte Cristo mine (CPAL.136, CPAL.155, CPAL.158, CPAL.173, CPAL.200, CPAL.214, CPAL.217). Totolapa, Río Salado mine (CPAL.210).

Horizon: Uppermost Simojovel Formation, Lower Miocene.

**Genus *Docodesmus*** Cook, 1896

*Docodesmus* sp.: Santiago-Blay & Poinar, 1992: 367, Fig.11.

Referred material: GOP DC-3-3: ♂; two ♀. Preservation: Amber inclusions. Repository: GOP; PC. Locality: Dominican Republic, Santiago, Cordillera Septentrional. Horizon: El Mamey Formation, Lower Miocene.

**(152) †Docodesmus brodzinskyi Shear, 1981**

*Docodesmus brodzinskyi* Shear, 1981: 53, Figs 1-2.

Referred material: Holotype: ♀. Preservation: Amber inclusion. Repository: JB. Locality: Dominican Republic, Santiago, No stated. Horizon: No stated, Lower Miocene.

**Genus *Iomus* Cook, 1911**

*Iomus* sp.: Santiago-Blay & Poinar, 1992: 367, Figs 12-13.

Referred material: GOP DC-3-6: ♀; GOP DC-3-5: ♂. Preservation: Amber inclusions. Repository: GOP. Locality: Dominican Republic, Santiago, Cordillera Septentrional. Horizon: El Mamey Formation, Lower Miocene.

**Genus *Lophodesmus* Pocock, 1894**

*Lophodesmus* sp.: Santiago-Blay & Poinar, 1992: 367, Fig.14.

Referred material: ♂. Preservation: Amber inclusion. Repository: PC. Locality: Dominican Republic, Santiago, Cordillera Septentrional. Horizon: El Mamey Formation, Lower Miocene.

**Genus *Myrmecodesmus* Silvestri, 1910**

**(153) †*Myrmecodesmus antiquus* Riquelme & Hernández-Patricio, 2021**

*Myrmecodesmus antiquus* Riquelme *et al*., 2021: 3, Figs 1-5.

Referred material: Holotype CPAL.132: ♂; Paratype CPAL.117: ♀. Preservation: Amber inclusions. Repository: CPAL-UAEM. Locality: México, Chiapas, Simojovel, La Pimienta mine (CPAL.132), Los Pocitos mine (CPAL.117). Horizon: Uppermost Simojovel Formation, Lower Miocene.

Genus *Psochodesmus* Cook, 1896

*Psochodesmus* sp.: Santiago-Blay & Poinar, 1992: 367, Fig.15.

Referred material: GOP DC-3-20: ♂. Preservation: Amber inclusion. Repository: GOP. Locality: Dominican Republic, Santiago, Cordillera Septentrional. Horizon: El Mamey Formation, Lower Miocene.

**Infraorder Polydesmoides Pocock, 1887**

**Superfamily Haplodesmoidea Cook, 1895**

**Family Haplodesmidae Cook, 1895**

**Genus *Inodesmus* Cook, 1896**

*Inodesmus* sp.: Santiago-Blay & Poinar, 1992: 367, Fig.17.

Referred material: GOP DC-3-2: 3 ♀; GOP DC-3-1: 3 juveniles, ♂; 1 specimen, ♂. Preservation: Amber inclusions. Repository: GOP; PC. Locality: Dominican Republic, Santiago, Cordillera Septentrional. Horizon: El Mamey Formation, Lower Miocene.

**Superfamily Polydesmoidea Leach, 1815**

**Family Polydesmidae Leach, 1815**

**Genus *Polydesmus* Latreille, 1802**

*Polydesmus* sp.: Hoffman, 1969: 595, Fig.378.

Referred material: Holotype. Preservation: Amber inclusion. Repository: No stated. Locality: Baltic. Horizon: Prussian Formation, Upper Eocene.

**Superfamily Trichopolydesmoidea Verhoeff, 1910**

**Family Trichopolydesmidae Verhoeff, 1910**

Trichopolydesmidae indet: Álvarez-Rodríguez *et al*., 2023, Figs 4A-C (this work).

New records: CPAL.119: ♀; CPAL.121: ♂; CPAL.150: ♂. Preservation: Amber inclusions. Repository: CPAL-UAEM. Locality: Mexico, Chiapas, Simojovel, Guadalupe Victoria mine (CPAL.121), Los Pocitos mine (CPAL.119, CPAL.150). Horizon: Uppermost Simojovel Formation, Lower Miocene.

Genus *Monstrodesmus* Golovatch, Geoffroy &VandenSpiegel, 2014

**(154) †Monstrodesmus grimaldii Su et al., 2022**

*Monstrodesmus grimaldii* Su *et al*., 2022: 607, Figs 1-14.

Referred material: Holotype: NIGP175093: ♂, Paratypes: NIGP175094: ♀ NIGP175096: ♂. Additional specimens: 5♀ and 2 indeterminate. Preservation: Amber inclusions. Repository: NGIP. Locality: Myanmar, Kachin, Hukawng Valley, Noije Bum mine. Horizon: No stated, lowermost Upper Cretaceous, Upper Albian/Lower Cenomanian.

**3.2.4 Subclass *Incertae sedis***

**Order Incertae sedis**

**†Family Proiulidae Fritsch, 1899**

**†Genus *Tomiulus* Martynov, 1936**

**(155) †*Tomiulus angulatus* Martynov, 1936**

*Tomiulus angulatus* Martynov, 1936: 1258.

Referred material: Holotype: PIN 1062/1bc. Preservation: Compression. Repository: PIN. Locality: Russian Federation, Kemerovo. Horizon: Maltsevo Formation, Lower Triassic.

**Family Incertae sedis**

**†Genus *Decorotergum* Jell, 1983**

**(156) †Decorotergum warrenae Jell, 1983**

*Decorotergum warrenae* Jell, 1983: 197, Figs 1-3, 196-198.

Referred material: Holotype: QMF12294; Paratype: QMF12295; QMF12296: one small fragment of a third specimen. Preservation: Compressions. Repository: QMF. Locality: Australia, Queensland, Kolane. Horizon: Evergreen Formation, Lower Jurassic.

#### 3.2.5 Diplopoda Indeterminate

Diplopoda indet: Pickford & Andrews, 1981: 32; Slaughter, 1966: 79; Shear *et al*., 1992: 136; Janossy, 1986: 20; Montoya *et al*., 2001: 388; Rasnitsyn & Ross, 2000: 24; Röβler *et al*., 2012: 819, Fig.12A; Ross *et al*., 2016: 3; Huang *et al*., 2018: 5, Fig.4; Riquelme & Hernández-Patricio, 2018: 643, Figs 3A-B, 642; Álvarez-Rodríguez *et al*., 2023 (this work).

Referred material: Locality 11: 16 specimens, Locality 29: 17 specimens. Preservation: Compressions. Repository: NHML. Locality: Kenya, Kimusu, sites 11 and 29. Horizon: Legetet Formation, Lower Miocene.

Referred material: No stated. Preservation: Compression. Repository: SMU. Locality: USA, Dallas County, Texas, Moore Pit Local Fauna. Horizon: Shuler Formation, Upper Pleistocene.

Referred material: No stated. Preservation: Impression. Repository: NMMNH. Locality: USA, New Mexico, Bernalillo County, Kinney Quarry. Horizon: Wild Cow Formation, Upper Carboniferous.

Referred material: No stated. Preservation: Compression. Repository: HNHM. Locality: Hungary, Barany. Horizon: Limestone strata, Pliocene.

Referred material: No stated. Preservation: Compression. Repository: GCPE. Locality: Spain, Murcia, Sierra de Quibas.

Horizon: No stated, Lower Pleistocene.

Referred material: In.1912. Preservation: Amber inclusion. Repository: NHML. Locality: Myanmar, Kachin, Hukawng Valley, Noije Bum mine. Horizon: No stated, lowermost Upper Cretaceous, Upper Albian/Lower Cenomanian.

Referred material: TA0851. Preservation: Impression. Repository: MfNC. Locality: Germany, Saxony, Chemnitz.

Horizon: Leukersdorf Formation, Lower Permian, Sakmarian.

Referred material: LPI-63009. Preservation: Compression. Repository: CDCGS. Locality: China, Yunnan, Luoping County, Luoping biota. Horizon: Member II of the Guanling Formation, Anisian, Middle Triassic.

Referred material: G.2005.147.2; G.2006.42.2. Preservation: Amber inclusions. Repository: NMS. Locality: Mexico, Chiapas. Horizon: Uppermost Simojovel Formation, Lower Miocene.

Referred material: MALM.303; MALM.305. Preservation: Amber inclusions. Repository: MALM. Locality: Mexico, Chiapas, Simojovel, La Pimienta mine. Horizon: Uppermost Simojovel Formation, Lower Miocene.

New records: CPAL.151; CPAL.154; CPAL.170: juvenile; CPAL.171: juvenile; CPAL.216; CPAL.159; CPAL.179; CPAL.208. Preservation: Amber inclusions. Repository: CPAL-UAEM. Locality: Mexico, Chiapas, Simojovel, Campo La Granja mine (CPAL.151), El Chapayal mine (CPAL.159), El Pabuchil mine (CPAL.179), Los Pocitos mine (CPAL.154), Monte Cristo mine (CPAL.208, CPAL.216). Huitiupán (CPAL.170, CPAL.171). Horizon: Uppermost Simojovel Formation, Lower Miocene.

#### 3.2.6 Summary taxonomic list of Diplopoda fossil record

Phylum **Arthropoda** Gravenhorst, 1843

Clade **Mandibulata** *sensu* Snodgrass, 1938

Subphylum **Myriapod**a Latreille, 1802

Class **Diplopoda** de Blainville in Gervais, 1844

Subclass **Penicillata** Latrielle, 1831 (1 order)

Order **Polyxenida** Verhoeff, 1934 (2 superfamilies)

Superfamily **Polyxenoidea** Lucas, 1940 (2 families)

Family **Lophoproctidae** Silvestri, 1897 (1 genus) Genus ***Lophoproctus*** Pocock, 1894

Family **Polyxenidae** Lucas, 1840 (6 genera)

† Genus ***Electroxenus*** Nguyen Duy–Jacquemin & Azar, 2004 (1 species)

† ***Electroxenus jezzinensis*** Nguyen Duy–Jacquemin & Azar 2004

† Genus ***Libanoxenus*** Nguyen Duy–Jacquemin & Azar, 2004 (1 species)

† ***Libanoxenus hammanaensis*** Nguyen Duy–Jacquemin & Azar, 2004

Genus ***Polyxenus*** Latrielle, 1802 (6 species)

† ***Polyxenus caudatus*** Menge, 1854

† ***Polyxenus colurus*** Menge, 1854

† ***Polyxenus coniformis*** Koch & Berendt, 1854

† ***Polyxenus lophurus*** Menge, 1854

† ***Polyxenus miocenica*** Srivastava *et al*., 2006

† ***Polyxenus ovalis*** Koch & Berendt, 1854

Genus ***Propolyxenus*** Silvestri, 1948

Genus ***Pauropsxenus*** Silvestri, 1948 (2 species)

† *Pauropsxenus extraneus* Su *et al*., 2020

† *Pauropsxenus ordinatus* Su *et al*., 2020

Genus ***Unixenus*** Jones, 1944

Superfamily **Synxenoidea** Silvestri, 1923 (1 family)

Family **Synxenidae** Silvestri, 1923 (1 genus)

Genus ***Phryssonotus*** Scudder, 1885 (2 species)

† ***Phryssonotus hystrix*** Menge, 1854

† ***Phryssonotus burmiticus*** (Cockerell, 1917)

† Subclass **Arthropleuridea** Waterlot, 1934 (3 orders)

† Order **Arthropleurida** Waterlot, 1934 (1 family)

† Family **Arthropleuridae** Scudder, 1885 (1 genus)

† Genus ***Arthropleura*** Jorden & Meyer, 1854 (8 species)

† ***Arhtropleura armata*** (Jordan & Meyer, 1854)

† ***Arthropleura britannica*** Andree, 1913

† ***Arthropleura cristata*** Richarson, 1959

† ***Arthropleura fayoli*** Boule, 1893

† ***Arthropleura maillieuxi*** Pruvost, 1930

† ***Arthropleura mammata* (**Salter, 1863)

**† *Arthropleura moyseyi*** Calman, 1915

† ***Arthropleura punctata*** Goldenberg, 1873

† Order **Eoarthropleurida** Shear & Selden, 1995 (1 family)

† Family **Eoarthropleuridae** Størmer, 1976 (1 genus)

† Genus ***Eoarthropleura*** Størmer, 1976 (3 species)

† ***Eoarthropleura devonica*** Størmer, 1976

† ***Eoarthropleura hueberi*** Kjellsvig–Waering, 1986

† ***Eoarthropleura ludfordensis*** Shear & Selden, 1995

† Order **Microdecemplicida** Wilson & Shear, 2000 (1 family)

† Family **Microdecemplicidae** Wilson & Shear, 2000 (1 genus)

† Genus ***Microdecemplex*** Wilson & Shear, 2000 (1 species)

† ***Microdecemplex rolfei*** Wilson & Shear, 2000

Subclass **Chilognatha** Latrielle, 1802 (2 infraclass)

† Order **Zosterogrammida** Wilson, 2005 (1 family)

† Family **Zosterogrammidae** Wilson, 2005 (1 genus)

† Genus ***Casiogrammus*** Wilson, 2005 (1 species)

† ***Casiogrammus ichthyeros*** Wilson, 2005

† Genus ***Purkynia*** Fritsch, 1899 (1 species)

† ***Purkynia lata*** Fritsch, 1899

† Genus ***Zosterogrammus*** Wilson, 2005 (1 species)

† ***Zosterogrammus stichostethu****s* Wilson, 2005

Infraclass **Pentazonia** Brandt, 1833 (2 superorders)

† Order **Amynilypedida** Hoffman, 1969 (2 families)

† Family **Amynilyspedidae** Hoffman, 1969 (1 genus)

† Genus ***Amynilyspes*** Scudder, 1882 (5 species)

† ***Amynilyspes crescens*** Fritsch, 1899

† ***Amynilyspes fatimae*** Racheboeuf *et al*., 2004

† ***Amynilyspes springhillensis*** Copeland, 1957

† ***Amynilyspes typicus*** Fritsch, 1899

† ***Amynilyspes wortheni*** Scudder, 1882

† Family **Sphaerherpestidae** Fritsch, 1899 (1 genus)

† Genus ***Glomeropsis*** Fritsch, 1895 (4 species)

† ***Glomeropsis crassa*** Fristch, 1899

† ***Glomeropsis magna*** Fritsch, 1899

† ***Glomeropsis multicarinata*** Fritsch, 1899

† ***Glomeropsis ovalis*** Fritsch, 1895

Superorder **Limacomorpha** Pocock, 1894 (1 order)

Order **Glomeridesmida** Cook, 1895 (1 family)

Family **Glomeridesmidae** Latzel, 1884 (1 genus)

Genus ***Glomeridesmus*** Gervais, 1844

Superorder **Oniscomorpha** Pocock, 1887 (2 orders)

Order **Glomerida** Brandt, 1833 (2 families)

Family **Glomeridellidae** Cook, 1896 (1 genus)

Genus ***Glomeridella*** Brölemann, 1895

Family **Glomeridae** Leach, 1816 (2 genera)

Genus ***Glomeris*** Latreille, 1802 (1 species)

† ***Glomeris denticulata*** Menge, 1854

Genus ***Hyleoglomeris*** Verhoeff, 1910 (1 species)

† ***Hyleoglomeris groehni*** Wesener, 2019

Order **Sphaerotheriida** Brandt, 1833 (1 family)

Family **Zephroniidae** Gray in Jones, 1843

Infraclass **Helminthomorpha** Pocock, 1887 (2 subterclasses)

Superorder *Incertae sedis*

Order *Incertae sedis*

Family *Incertae sedis*

† Genus ***Dolesea*** Hannibal & May, 2020 (1 species)

† ***Dolesea subtila*** Hannibal & May, 2020

† Genus ***Sinosoma*** Huang & Hannibal, 2018 (1 species)

† ***Sinosoma luopingense*** Huang & Hannibal, 2018

Superorder *Incertae sedis*

Order *Incertae sedis*

Family *Incertae sedis*

Genus ***Archicambala*** Cook, 1895 (1 species)

***Archicambala dawsoni*** (Scudder, 1868)

Superorder *Incertae sedis*

Order *Incertae sedis*

† Family **Archiulidae** Scudder, 1873 (1 genus)

† Genus ***Archiulus*** Scudder, 1868 (5 species)

† ***Archiulus brassi* (**Dohrn, 1868)

† ***Archiulus euphoberioides*** Scudder, 1895

† ***Archiulus glomeratus*** Sccuder, 1890

† ***Archiulus lyelli*** Scudder, 1895

† ***Archiulus xylobioides*** Sccuder, 1868

Superorder *Incertae sedis*

Order *Incertae sedis*

† Family **Oklahomasomatidae** Hannibal & May, 2020 (1 genus)

† Genus ***Oklahomasoma*** Hannibal & May, 2020 (1 species)

† ***Oklahomasoma richardsspurense*** Hannibal & May, 2020

† Superorder **Archipolypoda** Scudder, 1882 (5 orders)

Order *Incertae sedis*

Family *Incertae sedis*

† Genus ***Albadesmus*** Wilson & Anderson, 2004 (1 species)

† ***Albadesmus almondi*** Wilson & Anderson, 2004

Order *Incertae sedis*

Family *Incertae sedis*

† Genus ***Anaxeodesmus*** Wilson, 2005 (1 species)

† ***Anaxeodesmus diambonotus*** Wilson, 2005

Order *Incertae sedis*

Family *Incertae sedis*

† Genus ***Anthracodesmus*** Peach, 1899 (1 species)

† ***Anthracodesmus macconochiei*** Peach, 1899

Order *Incertae sedis*

Family *Incertae sedis*

† Genus ***Kampecaris*** Page, 1856 (3 species)

† ***Kampecaris dinmorensis*** Clarke, 1951

† ***Kampecaris forfarensis*** Peach, 1882

† ***Kampecaris obanesis*** Peach, 1889

Order *Incertae sedis*

Family *Incertae sedis*

† Genus ***Palaeodesmus*** Brade–Birks, 1923 (1 species)

† ***Palaeodesmus tuberculata*** (Brade–Birks, 1923)

Order *Incertae sedis*

Family *Incertae sedis*

† Genus ***Pneumodesmus*** Wilson & Anderson, 2004 (1 species)

† ***Pneumodesmus newmani*** Wilson & Anderson, 2004

Order *Incertae sedis*

† Family **Woodesmidae** Ross *et al*., 2008 (1 genus)

† Genus ***Woodesmus*** Ross *et al*., 2018 (1 species)

† ***Woodesmus sheari*** Ross *et al*., 2018

† Order **Archidesmida** Wilson & Anderson, 2004 (2 families)

† Family **Archidesmidae** Scudder, 1885 (1 genus)

† Genus ***Archidesmus*** Peach, 1882 (1 species)

† ***Archidesmus macnicoli*** Peach, 1882

† Family **Zanclodesmidae** Wilson *et al*., 2005 (2 genera)

† Genus ***Orsadesmus*** Wilson *et al*., 2005 (1 species)

† *Orsadesmus rubecollus* Wilson *et al*., 2005

† Genus ***Zanclodesmus*** Wilson *et al*., 2005 (1 species)

† *Zanclodesmus willetti* Wilson *et al*., 2005

† Order **Cowiedesmida** Wilson & Anderson, 2004 (1 family)

† Family **Cowiedesmidae** Wilson & Anderson, 2004 (1 genus)

† Genus ***Cowiedesmus*** Wilson & Anderson, 2004 (1 species)

† ***Cowiedesmus eroticopodus*** Wilson & Anderson, 2004

† Order **Euphoberiida** Hoffman, 1969 (1 family)

† Family **Euphoberiidae** Scudder, 1882 (3 genera)

† Genus ***Acantherpestes*** Meek & Worthen, 1868 (6 species)

† ***Acantherpestes foveolatus*** Fritsch, 1899

† ***Acantherpestes gigas*** Fritsch, 1895

† ***Acantherpestes inequalis*** Scudder, 1890

† ***Acantherpestes major*** (Meek & Worthen, 1868)

† ***Acantherpestes ornatus*** Fritsch, 1899

† ***Acantherpestes vicinus*** Fritsch, 1899

† Genus ***Euphoberia*** Meek & Worthen, 1868 (17 species)

† ***Euphoberia absens*** Fritsch, 1899

† ***Euphoberia anguilla*** Scudder, 1882

† ***Euphoberia armigera*** Meek & Worthen, 1868

† ***Euphoberia brownii*** Woodward, 1871

† ***Euphoberia carri*** Scudder, 1882

† ***Euphoberia cuspidata*** Scudder, 1890

† ***Euphoberia ferox*** (Salter, 1863)

† ***Euphoberia flabellata*** Scudder, 1882

† ***Euphoberia granosa*** Scudder, 1882

† ***Euphoberia histrix*** Fritsch, 1899

† ***Euphoberia horrida*** Scudder, 1882

† ***Euphoberia hystricosa*** Scudder, 1890

† ***Euphoberia lithanthracis* (**Jordan & Meyer, 1854)

† ***Euphoberia simplex*** Scudder, 1890

† ***Euphoberia spinulosa*** Scudder, 1890

† ***Euphoberia tracta*** Scudder, 1890

† ***Euphoberia varians*** Fritsch, 1899

† Genus ***Myriacantherpestes*** Burke, 1979

† Order **Palaeosomatida** Hannibal & Krzeminski, 2005 (1 family)

† Family **Palaeosomatidae** Hannibal & Krzeminski, 2005 (1 genus)

† Genus ***Paleosoma*** Jackson *et al*., 1919 (2 species)

† ***Paleosoma giganteus* (**Baldwin, 1911)

† **Palaeosoma robustum (**Jackson *et al*., 1919)

Subterclass **Colobognatha** Brandt, 1834 (3 orders)

† Order **Pleurojulida** Schneider & Werneburg, 1998 (1 family)

† Family **Pleurojulidae** Schneider & Werneburg, 1998 (2 genera)

† Genus ***Pleurojulus*** Fritsch, 1899 (2 species)

† ***Pleurojulus biornatus*** Fritsch, 1899

†***Pleurojulus levis* (**Fritsch, 1899)

† Genus ***Isojulus*** Fritsch, 1899 (1 species)

† ***Isojulus constans* (**Frič, 1875)

Order **Platydesmida** Cook, 1895 (1 family)

Family **Andrognathidae** Cope, 1869 (2 genera)

Genus ***Brachycybe*** Wood, 1864

Genus ***Andrognathus*** Cope, 1869 (1 species)

† ***Andrognathus burmiticus*** Moritz & Wesener, 2019

Order **Polyzoniida** Cook, 1895 (2 families)

Family **Polyzoniidae** Newport, 1844 (1 genus)

Genus ***Polyzonium*** Brandt, 1837

Family **Siphonotidae** Cook, 1895 (1 genus)

Genus ***Siphonotus*** Brandt, 1837

Order **Siphonophorida** Newport, 1844 (2 families)

Family **Siphonophoridae** Newport, 1844 (2 genera)

Genus ***Siphonophora*** Brandt, 1837 (3 species)

† ***Siphonophora hui*** Jiang *et al*., 2019

† ***Siphonophora hoffmani*** Santiago–Blay & Poinar, 1992

† ***Siphonophora velezi*** Santiago–Blay & Poinar, 1992

Genus ***Siphonocybe*** Pocock, 1903

Family **Siphonorhinidae** Cook, 1895

Subterclass **Eugnatha** Attems, 1898 (3 superorders)

Superorder **Juliformia** Attems, 1926 (3 orders, 1 superfamily)

Order *Incertae sedis*

† Family **Proglomeridae** Fritsch, 1899 (1 genus)

† Genus ***Archiscudderia*** Fritsch, 1899 (5 species)

† ***Archiscudderia coronata*** Fritsch, 1899

† ***Archiscudderia paupera*** Fritsch, 1899

† ***Archiscudderia problematica*** Fritsch, 1899

† ***Archiscudderia regularis*** Fritsch, 1899

† ***Archiscudderia tapeta*** Fritsch, 1899

† Superfamily **Xyloiuloidea** Attems, 1926 (4 families)

Family *Incertae sedis*

† Genus ***Karstiulus*** Hannibal & May, 2020 (1 species)

† ***Karstiulus fortsillensis*** Hannibal & May, 2020

† Family **Gaspestriidae** Wilson, 2006 (2 genera)

† Genus ***Sigmastria*** Wilson, 2006 (1 species)

† ***Sigmastria dilata*** Wilson, 2006

† Genus ***Gaspestria*** Wilson, 2006 (1 species)

† ***Gaspestria genselorum*** Wilson, 2006

† Family **Nyraniidae** Hoffman, 1969 (1 genus)

† Genus ***Nyranius*** Frič, 1875 (2 species)

† ***Nyranius costulatus*** (Frič, 1875)

† ***Nyranius tabulatus*** (Fritsch, 1899)

† Family **Proglomeridae** Fritsch, 1899

† Family **Xyloiulidae** Cook, 1895 (1 genus)

† Genus ***Blanziulus*** Langiaux & Sotty, 1976 (1 species)

† ***Blanziulus parriati*** Langiaux & Sotty, 197

† Genus ***Xyloiulus*** Cook, 1895 (10 species)

† ***Xyloiulus bairdi*** Hoffman, 1963

† ***Xyloiulus fractus*** Scudder, 1868

† ***Xyloiulus frustulentus* (**Scudder, 1890)

† ***Xyloiulus mazonus* (**Scudder, 1890)

†***Xyloiulus platti*** (Baldwin, 1911)

† ***Xyloiulus pstrossi*** (Fritsch, 1899)

† ***Xyloiulus renieri*** Pruvost, 1930

† ***Xyloiulus sellatus*** (Fritsch, 1899)

† ***Xyloiulus sigillariae*** (Dawson, 1860)

† ***Xyloiulus similis*** (Scudder, 1868)

Order **Julida** Brandt, 1833 (2 superfamilies)

Superfamily **Juloidea** Leach, 1814 (1 family)

Family **Julidae** Leach, 1814 (2 genera)

† Genus ***Bertkaupolypus*** Verhoeff, 1926 (1 species)

† ***Bertkaupolypus antiquus*** (Bertkau, 1878)

Genus ***Julus*** Linnaeus, 1758 (10 species)

† ***Julus badius*** Menge, 1854

† ***Julus cavicola*** Grinnell, 1908

† ***Julus florissantellus*** Cockerell, 1907

† ***Julus laevigatus*** Koch & Berendt, 1854

† ***Julus occidentalis*** Grinnell, 1908

**† *Julus peii*** Chia & Liu, 1950

† ***Julus politus*** Menge, 1854

† ***Julus rubens*** Menge, 1854

† ***Julus suevicus*** Dietlen, 1902

† ***Julus terrestris*** Chia & Liu, 1950

Superfamily **Parajuloidea** Bollman, 1893 (1 family)

Family **Parajulidae** Bollman, 1893 (1 genus)

Genus ***Parajulus*** Humbert & Saussure, 1869 (2 species)

† ***Parajulus cockerelli*** Miner, 1926

† ***Parajulus onychis*** Pierce, 1951

Order **Spirobolida** Cook, 1895 (1 suborder)

Suborder **Spirobolidea** Cook, 1895 (3 families)

Family **Atopetholidae** Chamberlin, 1918 (1 genus)

† Genus ***Gobiulus*** Dzik, 1975 (1 species)

† ***Gobiulus sabulosus*** Dzik, 1975

Family **Spirobolidae** Bollman, 1893 (1 genus)

Genus ***Hiltonius*** Chamberlin, 1918 (1 species)

† ***Hiltonius australis*** (Grinnell, 1908)

Family **Rhinocricidae** Brölemann, 1913 (1 genus)

Genus ***Rhinocricus*** Karsch, 1881

Order **Spirostreptida** Brandt, 1833 (1 suborder)

Suborder **Cambalidea** Cook, 1895 (2 families)

Family **Cambalidae** Cook, 1895 (1 genus)

† Genus ***Protosilvestria*** Handschin, 1944 (1 species)

† ***Protosilvestria sculpta*** Handschin, 1944

† Family **Electrocambalidae** Moritz & Wesener, 2021 (2 genera)

† Genus ***Electrocambala*** Moritz & Wesener, 2021 (2 species)

† ***Electrocambala cretacea*** Moritz & Wesener, 2021

† ***Electrocambala ornata*** Moritz & Wesener, 2021

† Genus ***Kachincambala*** Moritz & Wesener, 2021 (2 species)

† ***Kachincambala distorta*** Moritz & Wesener, 2021

† ***Kachincambala muelleri*** Moritz & Wesener, 2021

Family **Pseudonannolenidae** Silvestri, 1895 (1 genus)

Genus ***Epinannolene*** Brölemann, 1903

Superorder **Nematophora** Verhoeff, 1913 (4 orders)

Order **Callipodida** Pocock, 1894 (1 suborder)

Family *Incertae sedis*

† Genus ***Hannibaliulus*** Shear *et al*., 2009 (1 species)

† *Hannibaliulus wilsonae* Shear *et al*., 2009

† Suborder **Burmanopetalidea** Stoev, Moritz & Wesener, 2019 (1 family)

† Family **Burmanopetalidae** Stoev, Moritz & Wesener, 2019 (1 genus)

† Genus ***Burmanopetalum*** Stoev, Moritz & Wesener, 2019 (1 species)

† ***Burmanopetalum inexpectatum*** Stoev, Moritz & Wesener, 2019

Order **Chordeumatida** Pocock, 1894 (2 suborders)

Suborder **Craspedosomatidea** Cook, 1895 (1 superfamily)

Superfamily **Craspedosomatoidea** Gray in Jones, 1843 (1 family)

Family **Craspedosomatidae** Gray in Jones, 1843 (2 genera)

Genus ***Craspedosoma*** Leach, 1814 (7 species)

† ***Craspedosoma aculeatum*** Menge, 1854

† ***Craspedosoma affine*** Koch & Berendt, 1854

† ***Craspedosoma angulatum*** Koch & Berendt, 1854

† ***Craspedosoma armatum*** Menge, 1854

† ***Craspedosoma cylindricum*** Menge, 1854

† ***Craspedosoma obtusangulum*** Menge, 1854

† ***Craspedosoma setosum*** Menge, 1854

† Genus ***Euzonus*** Menge, 1854 (1 species)

† ***Euzonus collulum*** Menge, 1854

Suborder **Heterochordeumatidea** Shear, 2000 (1 superfamily)

Superfamily **Heterochordeumatoidea** Pocock, 1894 (1 family)

Family **Heterochordeumatidae** Pocock, 1894

Order **Stemmiulida** Pocock, 1895 (1 family)

Family **Stemmiulidae** Pocock, 1894 (1 genus)

† Genus ***Parastemmiulus*** Riquelme *et al*., 2013 (1 species)

† *Parastemmiulus elektron* Riquelme *et al*., 2013

Order **Siphoniulida** Cook, 1895 (1 family)

Family **Siphoniulidae** Pocock, 1894 (1 genus)

Genus ***Siphoniulus*** Pocock, 1894 (2 species)

† *Siphoniulus muelleri* Liu *et al*., 2017

† ***Siphoniulus preciosus*** Liu *et al*., 2017 Superorder **Merochaeta** Cook, 1895 (1 order)

Order **Polydesmida** Pocock, 1887 (2 suborders)

Suborder **Leptodesmidea** Brölemann, 1916 (4 superfamilies)

Superfamily **Chelodesmoidea** Cook, 1895 (1 family)

Family **Chelodesmidae** Cook, 1895 (3 genera)

† Genus ***Maatidesmus*** Riquelme & Hernández–Patricio, 2014 (1 species)

† ***Maatidesmus paachtun*** Riquelme & Hernández–Patricio, 2014

Genus ***Caraibodesmus*** Chamberlin, 1918 (1 species)

† ***Caraibodesmus verrucosus*** Pocock, 1894

Genus ***Chondrotropis*** Loomis, 1936

Superfamily **Platyrhacoidea** Pocock, 1895 (1 family)

Family **Platyrhacidae** Pocock, 1895 (2 genera)

† Genus ***Anbarrhacus*** Riquelme & Hernández–Patricio, 2014 (1 species)

† ***Anbarrhacus adamantis*** Riquelme & Hernández–Patricio, 2014

Genus ***Nyssodesmus*** Cook, 1896

Superfamily **Sphaeriodesmoidea** Humbert & DeSaussure, 1869 (1 family)

Family **Sphaeriodesmidae** Humbert & DeSaussure, 1869 (1 genus)

Genus ***Cyclodesmus*** Humbert & De Saussure, 1869 (1 species)

***Cyclodesmus porcellanus*** Pocock, 1894 Superfamily **Xystodesmoidea** Cook, 1895 (1 family)

Family **Xystodesmidae** Cook, 1895

Suborder **Strongylosomatidea** Brölemann, 1916 (1 family)

Family **Paradoxosomatidae** Daday, 1889 (1 genus)

Genus ***Orthomorpha*** Bollman, 1893 (1 species)

***Orthomorpha coarctata*** (Saussure, 1860) Suborder **Polydesmidea** Pocock, 1887 (2 infraorders)

Infraorder *Incertae sedis*

Superfamily *Incertae sedis*

Family *Incertae sedis*

Genus ***Dasyodontus*** Loomis, 1936

Infraorder **Oniscodesmoides** Simonsen, 1990 (1 superfamily)

Superfamily **Pyrgodesmoidea** Silvestri, 1896 (1 family)

Family **Pyrgodesmidae** Silvestri, 1896 (5 genera)

Genus ***Docodesmus*** Cook, 1896 (1 species)

† ***Docodesmus brodzinskyi*** Shear, 1981 Genus ***Iomus*** Cook, 1911

Genus ***Lophodesmus*** Pocock, 1894

Genus ***Myrmecodesmus*** Silvestri, 1910 (1 species)

† ***Myrmecodesmus antiquus*** Riquelme & Hernández–Patricio, 2021

Genus ***Psochodesmus*** Cook, 1896

Infraorder **Polydesmoides** Pocock, 1887 (3 superfamilies)

Superfamily **Haplodesmoidea** Cook, 1895 (1 family)

Family **Haplodesmidae** Cook, 1895 (1 genus) Genus ***Inodesmus*** Cook, 1896

Superfamily **Polydesmoidea** Leach, 1815 (1 family)

Family **Polydesmidae** Leach, 1815 (1 genus)

Genus ***Polydesmus*** Latreille, 1802

Superfamily **Trichopolydesmoidea** Verhoeff 1910 (1 family)

Family **Trichopolydesmidae** Verhoeff, 1910 (1 genus)

Genus ***Monstrodesmus*** Golovatch, Geoffroy &VandenSpiegel, 2014 (1 species)

† *Monstrodesmus grimaldii* Su *et al*., 2022

***Incertae sedis***

Subclass *Incertae sedis*

Order *Incertae sedis*

† Family **Proiulidae** Fritsch, 1899 (1 genus)

† Genus ***Tomiulus*** Martynov, 1936 (1 species)

† ***Tomiulus angulatus*** Martynov, 1936 Family *Incertae sedis*

† Genus ***Decorotergum*** Jell, 1983 (1 species)

† ***Decorotergum warrenae*** Jell, 1983

## Acknowledgments

MAR was supported by the CONACYT fellowship as part of the MMRN postgraduate program at the UAEM. We thank Susana Guzmán-Gómez at the LMF2-LANABIO, Instituto de Biología-UNAM, for photomicrography assistance. We thank Luis Zúñiga for access to the MALM and Bibiano Luna for access to the MACH. Both museums in San Cristóbal de las Casas, Chiapas, have collections of Mexican amber inclusions. We also thank the Academic Editor, Fuqiang Chen, and another annonimous rewiever whose comments and corrections have improved the final published version of this paper.

## References

Almond, J.E. 1985. The Silurian-Devonian fossil record of the Myriapoda. Philosophical Transactions of the Royal Society of London B, 309: 227–237. doi: 10.1098/rstb.1985.0082

Anderson, L.I., Dunlop, J.A., Horrocks, C.A., Winkelmann, H.M., Eagar, R.M.C. 1997. Exceptionally preserved fossils from Bickershaw, Lancashire UK (Upper Carboniferous, Westphalian A (Langsettian)). Geological Journal, 32: 197–210. doi: 10.1002/(SICI)1099-1034(199709) 32:33.0.CO;2-6

Andrée, K. 1913. Weiteres über das carbonische Arthostraken-genus *Arthopleura* Jordan. Palaeontographica, 60: 295–310.

Baalbergen, E., Donovan, S.K. 2012. Terrestrial arthropods from the Upper Pleistocene of Jamaica: systematics, palaeoecology and taphonomy. Geological Journal, 48(6): 628–645. doi: 10.1002/gj.2477

Baird, D. 1958. New records of Paleozoic Diplopod Myriapoda. Journal of Paleontology, 32(1): 239–241.

Baldwin, W. 1911. Fossil Myriapods from the Middle Coal-measures of Sparth Bottoms, Rochdale, Lancashire. Geological Magazine, 2: 74–80.

Bertkau, P. 1878. Einige Spinnen und ein Myriapode aus der Braunkohle von Rott. *Verhandlungen des Naturhistorischen Vereins der Preussischen Rheinlande und Westfalens*, Bonn, 35: 346–360.

Boule, M. 1893. Sur les débris d’*Arthropleur*a trouvés en France. Bulletin de la Société de l’Industrie Minérale, 7: 619–638.

Brade-Birks, S.G. 1923. Notes on Myriapoda, xxxviii. Kampecaris tuberculata, n. sp., from the Old Red Sandstone of Ayrshire. Proceedings of the Royal Physical Society of Edinburgh, 20: 277–280.

Briggs, D.E.G., Rolfe, W.D.I., Brannan, J. 1979. A giant myriapod trail from the Namurian of Arran, Scotland. Palaeontology, 22: 273–291.

Briggs, D.E.G., Plint, A.G., Pickerill, R.K. 1984. *Arthropleura* trails from the Westphalian of eastern Canada. Palaeontology, 27: 843–855.

Briggs, D. E. G. 1986. Walking trails of the giant arthropod Arthropleura. Bulletin Trimestriel de la Société d’Histoire Naturelle et des Amis du Muséum d’Autun, 117: 141–147.

Briggs, D.E.G., Almond, J.E. 1994. The arthropleurids from the Stephanian (Late Carboniferous) of Montceau-les-Mines (Massif Central - France). Mémoires de la Section des Sciences, 12: 127–135.

Brookfield, M.E., Catlos, E.J., Suarez, S.E. 2020. Myriapod divergence times differ between molecular clock and fossil evidence: U/Pb zircon ages of the earliest fossil millipede-bearing sediments and their significance. Historical Biology, 33(10): 2009–2013. doi: 10.1080/08912963.2020.1761351

Calder, J., Rygel, M., Ryan, R., Falcon-Lang, H., Herbert, B. 2005. Stratigraphy and sedimentology of early Pennsylvanian red beds at Lower Cove, Nova Scotia, Canada: the Little River Formation with redefinition of the Joggins Formation. Atlantic Geology, 41: 143–167. doi: 10.4138/183

Calman, W.T. 1914. III.—On Arthropleura moyseyi, n. sp., from the Coal-Measures of Debyshire. Geological Magazine, 1: 541–544. doi: 10.1017/S0016756800153452

Castro, M.P. 1997. Hallazgos de “Arthropleura” en el Estefaniense de la Península Ibérica. Revista Española de Paleontología, 12: 15–22.

Chaney, D.S., Lucas, S.G., Elrick, S. 2013. New occurrence of an arthropleurid trackway from the lower Permian of Utah. *In*: Lucas, S.G., DiMichele, W.A., Barrick, J.E., Schneider, J.W., Spielmann, J.A. (eds.), The Carboniferous-Permian Transition. New Mexico Museum of Natural History and Science, Bulletin, 60. pp. 64–65.

Chia, L.P., Liu, H.T. 1950. Fossil Myriapods from Choukoutien. Bulletin of the Geological society of China, 30(1-4): 23–27.

Clarke, B.B. 1951. The geology of Dinmore Hill, Herefordshire, with a description of a new Myriapod from the Dittonian rocks there. Trans. Woolhope Naturalists Field Club, 33:222–236.

Cook, O. F. 1895. Introductory note on the families of Diplopoda, in The Craspedosomatidae of North America. Annals of the New York Academy of Sciences, 9:1–9.

Cockerell, T.D.A. 1907. Some fossil arthropods from Florissant, Colorado. Bulletin of the American Museum of Natural History, 23: 605–616.

Cockerell, T.D.A. 1917. Arthropods in Burmese amber. Psyche, 24: 40–45.

Conde, B. 1954. Les Diplopodes Penicillates de l’Ambre et de la faune actuelle. Bulletin de la Societ e zoologique de France, 79: 74–78.

Conde, B., Nguyen Duy-Jacquemin, M., 1963. Diplopodes pènicillates rècoltès à Bombay par P. A. Remy. Revue française d’Entomologie, 30 (1): 68–78.

Copeland, M.J. 1957. The arthropod fauna of the Upper Carboniferous rocks of the maritime provinces. Geological Survey of Canada Memoir, 286: 1–110.

Davies, N.S., Garwood, R., McMahon, W.J., Schneider, J.W., Shillito, A.P. 2021. The largest arthropod in Earth history: insights from newly discovered *Arthropleura* remains (Serpukhovian Stainmore Formation, Northumberland, England). Journal of the Geological Society, 179: 18. doi: 10.1144/jgs2021-115

Dawson, J.W. 1860. On a terrestrial mollusk, a chilognathous myriapod, and some new species of reptiles, from the Coal-Formation of Nova Scotia. Quarterly Journal of the Geological Society of London, 16: 268–277.

De La Comble, J. 1963. Un arthropode encore inconnu du Permien, *Arthropleura* spec. Bulletin de la Société d’Histoire Naturelle d’Autun, 28, 6.

Dernov, V. 2019. Taphonomy and paleoecology of fauna and flora from deltaic sandstones of Mospinka Formation (Middle Carboniferous) of Donets Basin. Geo&Bio, 18: 37–63. doi: 10.15407/gb1805

Dietlen, R. 1902. Nachtrag zu “*Julus* cfr. *antiquus* und sonstige Funde aus dem Böttinger Sprudelkalk.” Jahreshefte des Vereins für Vaterländische Naturkunde in Württemberg, (58): 83–85.

Dohrn, A., 1868. Julus brassii n. sp.: Ein Myriapode aus der Steinkohlenformation: Verhandlungen des naturhistorischen Vereines der preussischen Rheinlande und Westphalens, v. 3, p. 335–336.

Donovan, S.K., Veltkamp, C.J. 1994. Unusual preservation of late Quaternary millipedes from Jamaica. Lethaia, 27: 355–362.

Duncan, J.I., Briggs, D.E.G., Archer, M. 1998. Three-dimensionally mineralized insects and millipedes from the Tertiary of Riversleigh, Queensland, Australia. Palaeontology, 41(5): 835–851.

Dzik, J. 1975. Spiroboloid millipeds from the Upper Cretaceous of the Gobi Desert, Mongolia. Palaeontologia Polonica, 33: 17-23.

Edgecombe, G.D. 2015. Diplopoda: Fossils. In: Minelli, A. (ed.), Treatise on Zoology Anatomy, Taxonomy, Biology. The Myriapoda, Vol. 2. Brill, Netherlands. pp. 337–352.

Enghoff, H., Golovatch, S., Short, M., Stoev, P., Wesener, T. 2015. Diplopoda: Taxonomic verview. In: Minelli, A. (ed.), Treatise on Zoology Anatomy, Taxonomy, Biology. The Myriapoda, Vol. 2. Brill, Netherlands. pp. 363–453.

Falcon-Lang, H.J., Benton, M.J., Braddy, S.J., Davies, S.J. 2006. The Pennsylvanian tropical biome reconstructed from the Joggins Formation of Nova Scotia, Canada. Journal of the Geological Society, London, 163: 561–576. doi: 10.1144/0016-764905-063

Falcon-Lang, H.J., Miller, R.F. 2007. Palaeoenvironments and palaeoecology of the Early Pennsylvanian Lancaster Formation (‘Fern Ledges’) of Saint John, New Brunswick, Canada. *Journal of the Geological Society*, London, 164: 945–957. doi: 10.1144/0016-76492006-189

Falcon-Lang, H.J., Minter, N.J., Bashforth, A.R., Gibling, M.R., Miller, R.F. 2015. Mid-Carboniferous diversification of continental ecosystems inferred from trace fossil suites in the Tynemouth Creek Formation of New Brunswick, Canada. Palaeogeography, Palaeoclimatology, Palaeoecology, 440: 142–160. doi: 10.1016/j.palaeo.2015.09.002

Ferguson, L. 1966. The recovery of some large track-bearing slabs from Joggins, Nova Scotia. Atlantic Geology, 2: 128–130. doi: 10.4138/1501

Frič, A. 1875. Über die Fauna der Gaskohle des Pilsner und Rakonitzer Beckens. Sitzungsberichte der Königlichen Böhmischen Gesellschaft der Wissenschaften, Mathematisch-naturwissenschaftlichen Classe, 1875: 70–79.

Fritsch, A. 1895. Vorläufiger Bericht über die Arthropoden und Mollusken der böhmische Permformation. Sitzungsberichte der Königlich-Böhmischen Gesellschaft der Wissenschaften, Mathematisch-Naturwissenschaftliche Classe, 1894(36): 1–4.

Fritsch, A. 1899. Fauna der Gaskohle und der Kalkstein der Permformation Böhmens*, Vol.* 4. pp. 13-56.

Geinitz, H.B. 1855. Die Versteinerungen der Steinkohlenformation in Sachsen. W. Engelmann, Leipzig, 59 pp. 36 pls.

Getty, P., Sproule, R., Stimson, M., Lyons, P. 2017. Invertebrate trace fossils from the Pennsylvanian Rhode Island formation of Massachusetts, USA. Atlantic Geology, 53: 185–206. doi: 10.4138/atlgeol.2017.007

Goldenberg, F. 1873. Fauna Saraepontana Fossilis. Die Fossilen Thiere aus der Steinkohlenformation von Saarbrücken, 1tes Heft. Saarbrücken. pp. 1–26.

Golovatch, S.I. 2013. A reclassification of the millipede superfamily Trichopolydesmoidea, with descriptions of two new species from the Aegean region (Diplopoda, Polydesmida). ZooKeys, 340: 63–78. doi.org/10.3897/zookeys.340.6295

Grimaldi, D.A., Engel, M.S., Nascimbene, P.C. 2002. Fossiliferous amber from Myanmar (Burma): its rediscovery, biotic diversity, and paleontological significance. American Museum Novitates, 3361: 1–72. doi. 10.1206/0003-0082(2002)3612.0.CO;2

Grinnell, F. 1908. Quaternary myriopods and insects of California. Bulletin of the Department of Geology, 5(12): 207–215.

Guthörl, P. 1934. Die Arthropoden aus dem Carbon und Perm des Saar-Nahe-Pfalz-Gebietes. Abhandlungen der Preußischen Geologischen Landesanstalt Berlin, 164: 1–219.

Hahn, G., Hahn, R., Brauckmann, C. 1986. Zur Kenntnis von *Arthropleura* (Myriapoda; Ober-Karbon). Geologica et Palaeontologica, 20, 125–137.

Handschin, E. 1944. Insekten aus den Phosphoriten des Quercy. Schweizerische Paläontologische Abhandlungen, 64(4): 1–23.

Hannibal, J.T. 1997. Remains of *Arthropleura*, a gigantic myriapod arthropod, from the Pennsylvanian of Ohio and Pennsylvania. Kirtlandia, 50: 1–9.

Hannibal, J.T. Feldmann, R.M. 1981. Systematics and functional morphology of oniscomorph millipedes (Arthropoda: Diplopoda) from the Carboniferous of North America. Journal of Paleontology, 55: 730–746.

Hannibal, J.T. Krzemiński, W. 2005. A palaeosomatid millipede (Archipolypoda: Palaeosomatida) from the Carboniferous (Namurian A) of Silesia. Poland. Polskie Pismo Entomologiczne, 74: 205–217.

Hannibal, J.T. 2006. Millipedes (Diplopoda) from the Fort Sill fissures (Lower Permian) of southwestern Oklahoma: rare examples of Permian millipedes and of fossil millipedes from a Paleozoic fissure fill. Geological Society of America Abstracts with Programs, 38 (7): 553.

Hannibal, J.T., May, W.J. 2020. Permian millipedes from the Fort Sill fissures of southwestern Oklahoma, with comments on allied taxa and millipedes preserved in karstic environments. Journal of Paleontology, 95: 586–600. doi: 10.1017/jpa.2020.100

Hoffman, R.L. 1963. New genera and species of upper Paleozoic Diplopoda. Journal of Paleontology, 37: 167–174.

Hoffman, R. 1969. Myriapoda, exclusive of Insecta. *In*: Moore, R. (ed.), Treatise on Invertebrate Paleontology, Part R, Vol. 2. Geological Society of America and University of Kansas Press, Lawrence, KS. pp. R572–R606.

Huang, J., Hannibal, J., Feldmann, R., Zhang, Q, Hu, S., Schweitzer, C., Xie, T. 2018. A new millipede (Diplopoda, Helminthomorpha) from the Middle Triassic Luoping biota of Yunnan, Southwest China. Journal of Paleontology, 92(3): 478–487. doi: 10.1017/ jpa.2017.93

Jackson, J.W., Brade-Birks, H.K., Brade-Birks, S.G. 1919. Notes on Myriopoda. XIX. A revision of some fossil material from Sparth Bottoms, Lancs. *The Geological Magazine, New Series*, Decade, 6: 406–411.

Janossy, D. 1986. Pleistocene Vertebrate Faunas of Hungary. Developments in Palaeontology and Stratigraphy, 8. Elsevier, Amsterdam. 208pp.

Jell, P.A. 1983. An Lower Jurassic millipede from the Evergreen Formation in Queensland. Alcheringa, 7: 195–199.

Jiang, X., Shear, W. A., Hennen, D.A., Chen, H., Xie, Z. 2019. One hundred million years of stasis: *Siphonophora hui* sp. nov., the first Mesozoic sucking millipede (Diplopoda: Siphonophorida) from mid-Cretaceous Burmese amber. Cretaceous Research, 97: 34–39. doi: 10.1016/j.cretres.2019.01.011

Jordan, F.W.H., von Meyer, H. 1854. Über die Crustaceen der Steinkohlenformation von Saarbrücken. Palaeontographica, 4: 1–15.

Kjellesvig-Waering, E.N. 1986. A restudy of the fossil Scorpionida of the world. Palaeontographica Americana, 55: 1–287.

Kliver, M. 1883. Ueber einige neue Blattinarien-zwei *Dictyoneura*-und zwei *Arthropleura*-Arten aus der Saarbrücker Steinkohlenformation. Palaeontographica, 29: 249–267.

Koch, C. L., Berendt, G. C. 1854. Die im Bernstein befindlichen Crustaceen, Myriapoden, Arachniden und Apteren der Vorwelt. Die in Bernstein Befindlichen Organischen Reste der Vorwelt Gesammelt in Verbindung mit Mehreren Bearbeitetet und Herausgegeben, 1(2): 1–124.

Langiaux, J., Sotty, D. 1976. Première découverte d’un myriapode dans le Paléozoïque supérieur du Massif Central français. Revue Périodique de “La Physiophile,” 85: 42–46.

Langiaux, J., Sotty, D. 1977. Ichnologie 4: pistes et empreintes dans le Stéphanien de Blanzy-Montceau. La Physiophile, 86: 74–91.

Laurito, C.A., Valerio, A.L. 2018. Primer registro fósil de un Myriapoda (Polydesmida) para el Pleistoceno Tardío de la localidad de La Palmera de San Carlos, provincia de Alajuela, Costa Rica. Revista Geológica De América Central, 58: 179–187. doi: 10.15517/ rgac.v58i0.32848

Lheritier, M., Perroux, M., Vannier, J., Escarguel, G., Wesener, T., Moritz, L., Chabard, D., Adrien, J., Perrier, V. 2023. Fossils from the Montceau-les-Mines Lagerstätte (305 Ma) shed light on the anatomy, ecology and phylogeny of Carboniferous Millipedes. Journal of Systematic Palaeontology, 21(1): 2169891. doi: 10.1080/14772019.2023.2169891

Liu, W., Rühr, P. T., Wesener, T. 2017. A look with μCT technology into a treasure trove of fossils: The first two fossils of the millipede order Siphoniulida discovered in Cretaceous Burmese amber (Myriapoda, Diplopoda). Cretaceous Research, 74: 100–108.

Li, Y., Hu, S., Li, J., et al., 2021. Fossil *Orthomorpha* cf. *coarctata* (Arthropoda, Diplopoda, Paradoxosomatidae) from the Middle Pleistocene of Luonan, Shaanxi Province. Quaternary Sciences, 41(1): 304–307.

Lucas, S.G., Hunt, A.P., Calder, J.H., Reid, D., Hebert, B., Stimson, M. 2005a. Tetrapod footprints from Joggins, Nova Scotia: a template for understanding Carboniferous tetrapod footprints. Geological Association of Canada Annual Meeting, 30: 116–117.

Lucas, S.G., Lerner, A.J., Hannibal, J.T., Hunt, A.P., Schneider, J.W. 2005b. Trackway of a giant Arthropleura from the Upper Pennsylvanian of El Cobre Canyon, New Mexico. *In:* Geology of the Chama Basin, Lucas, S.G., Zeigler, K.E., Lueth, V.W., Owen, D.E. (eds), New Mexico Geological Society 56 th Annual Fall Field Conference Guidebook. pp. 279–282.

Mángano, M.G., Buatois, L.A., West, R.R., Maples, C.G. 2002. Ichnology of a Pennsylvanian Equatorial Tidal Flat: The Stull Shale Member at Waverly, Eastern Kansas. Kansas Geological Survey Bulletin, 245. The University of Kansas Printing Serverce, Lawrence, Kansas. 133pp.

Martino, R.L., Greb, S.F. 2009. Walking trails of the giant terrestrial arthropod *Arthropleura* from the Upper Carboniferous of Kentucky. Journal of Paleontology, 83: 140–146. doi: 10.1666/08-093R.1

Martynov, A.V. 1936. On some new materials of Arthropoda from Kuznetsk-Basin. Izvestiya Akademii Nauk SSSR: Seriya Biologicheskaya, 6: 1251–1264.

Mauries, J.P. 1992. Sur la vraie place due genre Protosilvestria Handschin dans la classification des diplopodes iuliformes (Iuliformia, Iulida, Cambalidea). Berichte des Naturwissenschaftlich-Medizinischen Vereins in Innsbruck, (Suppl)10: 23–31.

McClennen, M., Jenkins, J., Uhen, M., Bloom, D., Weiczorek, J. 2017. The Paleobiology Database (PBDB). Available from https://paleobiodb.org/#/ (accessed 14 February 2024).

Meek, F.B., Worthen, A.H. 1868. Preliminary notice of a scorpion, a Eurypterus?, and other fossils from the Coal-measures of Illinois. *The American Journal of Science and Arts*, Second Series, 46: 19–28.

Menge, A., 1854. Footnote. *In*: Koch, C.L., Berendt, G.C. (eds.), Die im Bernstein Befindlichen Crustaceen, Myriapoden, Arachniden, und Apteren der Vorwelt. Nicolaischen Buchhandlung, Berlin. p. 102.

Miner, R.W. 1926. A fossil myriapod of the genus Parajulus from Florissant, Colorado. American Museum Novitates, 219: 1–5.

Montoya, P., Alberdi, M.T., Barbadillo, L.J., van der Made, J., Morales, J., Murelaga, X., Peñalvera, E., Robles, F., Ruiz Bustos, A., Sánchez, A., Sanchizb, B., Soria, D., Szyndlar, Z. 2001. Une faune très diversifiée du Pléistocène inférieur de la Sierra de Quibas (province de Murcia, Espagne). Comptes Rendus de l’Académie Des Sciences - Series IIA - Earth and Planetary Science, 332(6): 387–393. doi: 10.1016/s1251-8050(01)01544-0

Moreau, J. D., Gand, G., Fara, E., Galtier, J., Aubert, N., Fouché, S. 2019. Trackways of *Arthropleura* from the Late Pennsylvanian of Graissessac (Hérault, southern France). Historical Biology, 33: 996–1007. doi: 10.1080/08912963.2019.1675055

Moritz, L., Wesener, T. 2019. The first known fossils of the Platydesmida an extant American genus in Cretaceous amber from Myanmar (Diplopoda: Platydesmida: Andrognathidae). Organisms Diversity & Evolution, 19(3): 423–433. doi: 10.1007/s13127-019-00408-0

Moritz, L., Wesener, T. 2021. Electrocambalidae fam. nov., a new family of Cambalidea from Cretaceous Burmese amber (Diplopoda, Spirostreptida). European Journal of Taxonomy, 755: 22–46. doi: 10.5852/ejt.2021.755.1397

Nelikhov, A. 2010. In the dungeons of Dzhezkazgan. PaleoMir, 1: 60–69.

Nguyen Duy-Jacquemin, M., Azar, D. 2004. The oldest records of Polyxenida (Myriapoda, Diplopoda): new discoveries from the Cretaceous ambers of Lebanon and France. Geodiversitas, 26: 631–641.

Novozhylov, N. 1962. Family Arthropleuridae Zittel, 1848. *In*: Rodendorf, B.B. (ed.), Fundamentals of Paleontology. Arthropods. Tracheata and Chelicerata. Publishing House of the Academy of Sciences of the USSR. pp. 37–63.

Pacyna, G., Florjan, S., Borzęcki, R. 2012. New morphological features of *Arthropleura* sp. (Myriapoda, Diplopoda) based on new specimens from the Upper Carboniferous of Lower Silesia (Poland). Annales Societatis Geologorum Poloniae, 82: 121–126.

Pavela, M. 2018. Fossil species from the Carboniferous: Arthropleura Jordan & Meyer 1854. PALEO, 2018: 74.

Peach, B.N. 1882. On some fossil myriapods from the Lower Old Red Sandstone of Forfarshire. Proceedings of the Royal Physical Society of Edinburgh, 7: 177–188.

Peach, B.N. 1899. On some new myriapods from the Palaeozoic rocks of Scotland. Proceedings of the Royal Physical Society of Edinburgh, 14: 113–126.

Pearson, P.N. 1992. Walking traces of the giant myriapod *Arthropleura* from the Strathclyde Group (Lower Carboniferous) of Fife. Scottish Journal of Geology, 28: 127–133. doi: 10.1144/sjg28020127

Perrier, V., Charbonnier, S. 2014. The Montceau-les-Mines Lagerstätte (Late Carboniferous, France). Comptes Rendus Palevol, 13: 353–367. doi: 10.1016/j.crpv.2014.03.002

Pickford, M., Andrews, P. 1981. The Tinderet Miocence sequence in Kenya. Journal of Human Evolution, 10(1): 11–33. doi: 10.1016/ S0047-2484(81)80023-1

Pierce, W.D. 1951. Fossil arthropods from Onyx-Marble. Bulletin of the Southern California Academy of Sciences, 50: 34–49.

Pillola, G.L., Zoboli, D. 2021. First occurrence of *Arthropleura armata* (Myriapoda) in the Moscovian (Carboniferous) of SW Sardinia (Italy). Bollettino della Società Paleontologica Italiana, 60(1): 49–54. doi: 10.4435/BSPI.2021.01

Proctor, C.J. 1998. Arthropleurids from the Westphalian D of Writhlington Geological Nature Reserve, Somerset. *Proceedings of the Geologists’* Association, 109: 93–98. doi: 10.1016/S0016-7878(98)80009-6

Pruvost, P. 1919. Introduction à l’Étude du Terrain Houiller du Nord et du Pas-de-Calais: la Faune Continentale du Terrain Houiller du Nord de la France. 584pp.

Pruvost, P. 1930. La faune continentale du terrain houiller de la Belgique. Memoires du Musee Royal d’Histoire Naturelle de Belgique, 44: 103–282.

Racheboeuf, P.R., Hannibal, J.T., Vannier, J.M.C. 2004. A new species of the diplopod *Amynilyspes* (Oniscomorpha) from the Stephanian lagerstätte of Montceau-les-Mines, France. Journal of Paleontology, 78: 221–229. doi: 10.1666/0022-3360(2004)078%3C0221:ANSOTD%3E2.0.CO;2

Rasnitsyn, A.P, Ross, A.J. 2000. A preliminary list of arthropod families present in the Burmese amber collection at The Natural History Museum, London. Bulletin of the Natural History Museum: Geological Series, 56(1): 21–24.

Richardson, E.S.Jr. 1956. Pennsylvanian invertebrates of the Mazon Creek area, Illinois: Trilobitomorpha, Arthropleurida. Fieldiana Geology, 12(4): 9–76.

Richardson, E.S.Jr. 1959. Pennsylvanian invertebrates of the Mazon Creek area, Illinois; Trilobitomorpha, Arthropleurida II. Fieldana Geology, 12: 79–82.

Riquelme, F., Alvarado-Ortega, J., Ramos-Arias, M., Hernández, M., le Dez, I., Lee-Whiting, T.A., Ruvalcaba-Sil, J.L. 2013. A fossil stemmiulid millipede (Diplopoda: Stemmiulida) from the Miocene amber of Simojovel, Chiapas, Mexico. Historical Biology, 26(4): 415–427. doi: 10.1080/08912963.2013.778843

Riquelme, F., Hernández-Patricio, M., Martínez-Dávalos, A., Rodríguez-Villafuerte, M., Montejo-Cruz, M., Alvarado-Ortega, J., Ruvalcaba-Sil, J.L., Zúñiga-Mijangos, L. 2014. Two flat-backed polydesmidan millipedes from the Miocene Chiapas-Amber Lagerstätte, Mexico. PLoS ONE, 9(8): e105877. doi: 10.1371/journal.pone.0105877

Riquelme, F., Hernández-Patricio, F. 2018. The millipedes and centipedes of Chiapas amber. Check List, 14(4): 637–646. doi: 10.15560/14.4.637

Riquelme, F., Hernández-Patricio, M., Álvarez-Rodríguez, M. 2021. A Miocene pyrgodesmid millipede (Polydesmida: Pyrgodesmidae) from Mexico. PeerJ, 9: e10574. doi: 10.7717/peerj.10574

Rolfe, W.D.I. 1980. Early invertebrate terrestrial faunas. *In*: Panchen, A.L. (ed.), The Terrestrial Environment and the Origin of Land Vertebrates. Academic Press, London and New York. pp. 117–157.

Ross, A., Sheridan, A. 2013. Amazing Amber. NMS Enterprises Limited-Publishing, Edinburgh. 64pp.

Ross, A.J., Mellish, C.J., Crighton, B., York, P.V. 2016. A catalogue of the collections of Mexican amber at the Natural History Museum, London and National Museums Scotland, Edinburgh, UK. Boletín de la Sociedad Geológica Mexicana, 68(1): 45–55. doi: 10.18268/BSGM2016v68n1a7

Ross, A. J. 2018. Burmese (Myanmar) Amber Taxa, On-line Checklist v. 2018.2. 104pp. Available from http://www.nms.ac.uk/explore/stories/natural-world/burmese-amber/ (accessed 30 December 2023)

Ross, J.A., Edgecombe, G.D., Clark, N.D.L. 2018. Taxonomic names, in A new terrestrial millipede fauna of earliest Carboniferous (Tournaisian) age from southeastern Scotland helps fill ‘Romer’s Gap’. Earth and Environmental Science Transactions of the Royal Society of Edinburgh, 108: 99–110.

Röβler, R., Schneider, J.W. 1997. Eine bemerkenswerte Paläobiocoenose im Unterkarbon Middeleuropastt Fossilführung und Paläoenvironment der Hainichen-Subgruppe (Erzgebirge-Becken). Veröffentlichungen des Museum für Naturkunde Chemnitz, 20: 5–44.

Röβler, R., Zierold, T., Feng, Z., Kretzschmar, R., Merbitz, M., Annacker, V., Schneider, J.W. 2012. A snapshot of an early Permian ecosystem preserved by explosive volcanism: new results from the Chemnitz petrified forest, Germany. Palaios, 27: 814–834. doi: 10.2110/palo.2011.p11-112r

Ryan, R. 1986. Fossil myriapod trails in the Permo-Carboniferous strata of northern Nova Scotia, Canada. Atlantic Geology, 22: 156–161. doi: 10.4138/1604

Ryan, R.J., Boehner, R.C. 1994. Geology of the Cumberland Basin, Cumberland, Colchester and Pictou Counties, Nova Scotia. Mines and Energy Branches, Memoir, 10. Department of Natural Resources, Nova Scotia. 222pp.

Salter, J.W. 1863. On some species of Eurypterus and allied forms. *Quarterly Journal of the Geological Society*, London, 19: 81–87. doi: 10.1144/GSL.JGS.1863.019.01-02.15

Santiago-Blay, J.A., Poinar, G.O. 1992. Millipedes from the Dominican amber, with the description of two new species (Diplopoda, Siphonophoridae) of Siphonophora. Annals of the Entomological Society of America, 85: 363–369.

Schneider, J., Barthel, M. 1997. Eine Taphocoenose mit *Arthropleura* (Arthropoda) aus dem Rotliegend (?Unterperm) des Döhlen-Becken (ElbeZone, Sachsen). Freiberger Forschungsheft C, 466: 183–223.

Schneider, J.W., Werneburg, R. 1998. Arthropleura und Diplopoda (Arthropoda) aus dem Unter-Rotliegend (Unter-Perm, Assel) des Thüringer Waldes (Südwest-Saale-Senke). Veröffentlichungen Naturhistorisches Museum Schleusingen, 13: 19–36.

Schneider, J.W., Lucas, S.G., Werneburg, R., Röβler, R. 2010. Euramerican Late Pennsylvanian/Early Permian arthropleurid/tetrapod associations-implications for the habitat and paleobiology of the largest terrestrial arthropod. Carboniferous-Permian transition in Canon del Cobre, northern New Mexico. *In*: Lucas, S.G., Schneider, J.W., Spielmann, J. (eds), New Mexico Museum of Natural History and Science, Bulletin*, 49*. pp.49-70.

Scudder, S.H. 1882. Archipolypoda, a subordinal type of spined myriapods from the Carboniferous formation. Memoirs of the Boston Society of Natural History, 3: 143–182.

Scudder, S. H. 1886. Systematic review of our present knowledge of fossil Insects, including Myriapods and Arachnids. Bull. U.S. Geol. Survey no. 31, 128 pp.

Scudder, S.H. 1890. New Carboniferous Myriapoda from Illinois. Memoirs of the Boston Society of Natural History, 4: 417–442.

Scudder, S.H. 1895. Canadian fossil insects, myriapods and arachnids, 3. Notes upon myriapods and arachnids found in sigillarian stumps in the Nova Scotia coal field. Geological Survey of Canada Contributions to Canadian Palaeontology, 2(1): 57–66.

Shear, W.A. 1981. Two fossil millipedes from the Dominican amber (Diplopoda: Chytodesmidae, Siphonophoridae). Myriapodologica, 8: 51–54.

Shear, W.A., Hannibal, J.T., Kukalová-Peck, J. 1992. Terrestrial arthropods from Upper Pennsylvanian rocks at the Kinney Brick Quarry, New Mexico. New Mexico Bureau of Mines & Mineral Resources, Bulletin, 138: 135–141.

Shear, A.W., Selden, P. 1995. *Eoarthropleura* (Arthropoda, Arthropleurida) from the Silurian of Britain and the Devonian of North America. *Neues Jahrbuch für Geologie und Paläontologie*, Abhandlungen, 196(3): 347–375.

Shear, W.A. 1997. The fossil record and evolution of the Myriapoda. *In:* Fortey, R.A., Thomas, R.H. (eds), Arthropod Relationships. Systematics Association Special Volume 55. Chapman & Hall, London. pp. 211–219.

Shear, W.A., Selden, P.A., Gall, J.C. 2009. Millipedes from the Grès à Voltzia, Triassic of France, with comments on Mesozoic millipedes (Diplopoda: Helminthomorpha: Eugnatha). International Journal of Myriapodology, 3: 7–19.

Shear, W.A., Edgecombe, G.D. 2010. The geological record and phylogeny of the Myriapoda. Arthropod Structure & Development, 39: 174–190.

Shear, W.A. 2011. Class Diplopoda de Blainville in Gervais, 1844. *In:* Zhang, Z.Q. (ed.), Animal Biodiversity: An Outline of Higher-level Classification and Survey of Taxonomic Richness. Zootaxa, 3148: 169–164.

Sierwald, P., Shear, W.A., Shelley, R.M., Bond, J.E. 2003. Millipede phylogeny revisited in the light of the enigmatic order Siphoniulida. Journal of Zoological Systematics and Evolutionary Research, 41: 87–99.

Sierwald, P., Bond, J.E. 2007. Current status of the myriapod class Diplopoda (millipedes): taxonomic diversity and phylogeny. Annual Review of Entomology, 52: 401–420.

Slaughter, B.H. 1966. The Moore Pit local fauna; Pleistocene of Texas. Journal of Paleontology, 40(1): 78–91.

Srivastava, G.P., Shukla, M., Kumar, P., Kumar, M., Prakash. A. 2006. Record of pillbug (*Armadillidium*) and millipede (*Polyxenus*) remains from the resin lumps of Warkalli Formation (upper Tertiary), Kerala coast. Journal of the Geological Society of India, 67: 715–719.

Štamberg, S., Zajíc, J. 2008. *Carboniferous and Permian Faunas and their Occurrence in the Limnic Basins of the Czech Republic*. Museum of Eastern Bohemia, Hradec Králové. 224 pp.

Sterzel, J.T. 1918. Die organischen Reste des Kulms und Rotliegenden der Gegend von Chemnitz. Abhandlungen der Mathematische-Physischen Klasse der Sächsischen Akademie der Wissenschaften, 53: 203–315.

Stoev, P., Moritz, L., Wesener, T. 2019. Dwarfs under dinosaur legs: a new millipede of the order Callipodida (Diplopoda) from Cretaceous amber of Burma. ZooKeys, 84(1): 79–96. doi: 10.3897/zookeys.841.34991

Størmer, L. 1976. Arthropods from the Lower Devonian (Lower Emsian) of Alken an der Mosel, Germany. Part 5: Myriapoda and additional forms, with general remarks on fauna and problems regarding invasion of land by arthropods. Senckenbergiana Lethaea, 57(2/3): 87–183.

Su, Y., Cai, C., Huang, D. 2019. Revision of *Phryssonotus burmiticus* (Diplopoda, Polyxenida, Synxenidae) in mid-Cretaceous amber from Myanmar. Cretaceous Research, 93: 216–224. doi: 10.1016/j.cretres.2018.09.002

Su, Y., Cai, C., Huang, D. 2020. Two new species of the bristle millipede genus *Pauropsxenus* (Diplopoda, Polyxenidae) in mid-Cretaceous Burmese amber. Cretaceous Research, 111(1): 104427. doi: 10.1016/j.cretres.2020.104427

Su, Y.T., Cai, C.Y., Huang, D.Y. 2021. Morphological revision of *Siphonophora hui* (Myriapoda: Diplopoda: Siphonophoridae) from the mid-Cretaceous Burmese amber. Palaeoentomology, 004(3): 279–288.

Su, Y.T., Cai, C.Y. Huang, D.Y. 2022. A new species of Trichopolydesmidae (Myriapoda, Diplopoda, Polydesmida) from the mid-Cretaceous Burmese amber. Palaeoentomology, 5: 606–622. doi: 10.11646/palaeoentomology.5.6.10

Verhoeff, C. 1897. Diplopoden Rheinpreussens und Beiträge zur Biologie und vergleichenden Faunistik europäischer Diplopoden, Vorläufer zu einer rheinischen Diplopodenfauna. Decheniana, 53:186–280.

Vernon, R.D. 1912. On the geology and palaeontology of the Warwickshire Coalfield. *Quarterly Journal of the Geological Society*, London, 68: 587–638. doi: 10.1144/GSL.JGS.1912.068.01-04.35

Walter, H., Gaitzsch, B. 1988. Beiträge zur Ichnologie limnisch-terrestrischer Sedimentationsräume. Teil II: *Diplichnites minimus* n. ichnosp. aus dem Permosiles des Flechtinger Höhenzuges. Freiberger Forschungshefter, C427: 73–84.

Wesener, T. 2019. The oldest pill millipede fossil: A species of the Asiatic pill millipede genus *Hyleoglomeris* in Baltic amber (Diplopoda: Glomerida: Glomeridae). Zoologischer Anzeiger, 283: 40–45. doi: 10.1016/j.jcz.2019.08.009

Wesener, T., Moritz, L. 2018. Checklist of the Myriapoda in Cretaceous Burmese amber and a correction of the Myriapoda identified by Zhang (2017). Check List, 14(6): 1131–1140. doi: 10.15560/14.6.1131

Whyte, M.A. 2018. Mating trackways of a fossil giant millipede. Scottish Journal of Geology, 54: 63–68. doi: 10.1144/sjg2017-013

Wilson, H.M., Shear, W.A. 2000. Microdecemplicida, a new order of minute Arthropleurideans (Arthropoda, Myriapoda) from the Devonian of New York State, USA. Transactions of the Royal Society of Edinburgh. Earth Sciences, 90: 351–375.

Wilson, H.M., Anderson, L.I. 2004. Morphology and taxonomy of Paleozoic millipedes (Diplopoda: Chilognatha: Archipolypoda) from Scotland. Journal of Paleontology, 78: 169–184.

Wilson, H.M., Daeschler, E.B., Desbiens, S. 2005. New flat-backed archipolypodan millipedes from the Upper Devonian of North America. Journal of Paleontology, 79: 738–744.

Wilson, H.M. 2005a. A new genus of archipolypodan millipede from Coseley lagerstätte, Upper Carboniferous, UK. Palaeontology, 48: 1097–1100.

Wilson, H.M. 2005b. Zosterogrammida, a new order of millipedes from the Middle Silurian of Scotland and the Upper Carboniferous of Euramerica. Palaeontology, 48: 1101–1110.

Wilson, H.M., 2006. Juliformian millipedes from the Lower Devonian of Euramerica: implications for the timing of millipede cladogenesis in the Paleozoic. Journal of Paleontology, 80: 638–649.

Woodward, H. 1871. On *Euphoberia brownii*, H. Woodw., a new species of myriapod from the Coal-measures of the west of Scotland. The Geological Magazine, 8: 102–104.

Woodward, H. 1907. Further notes on the Arthropoda of the British CoalMeasures. Geological Magazine, 4: 539–549. doi: 10.1017/S0016756800134120

Zhang, W.W. 2017. Frozen Dimensions of the Fossil Insects and Other Invertebrates in Amber. Chonqing University Press, Chonqing. 697pp.

